# Myosin 5A mediates membrane-associated periodic skeleton reassembly during axon regeneration in response to ROCK-2 inhibition

**DOI:** 10.64898/2026.07.09.737468

**Authors:** Atrayee Basu, Elisa M. Howard, Tanina Arab, Nabab Khan, Jean Kanyo, TuKiet Lam, Stephen M. Strittmatter

**Author notes:** Department of Neuroscience, Stanford University School of Medicine, Stanford, CA, USA.

## Abstract

The membrane-associated periodic skeleton (MPS) is a submembrane lattice composed of actin rings and spectrin tetramers that repeats every 190 nm along axons and maintains mechanical stability. Loss of the MPS precedes axon fragmentation during degeneration, but during axon regrowth after injury the extent and timing of MPS reformation are not clear. We used stimulated emission depletion (STED) microscopy to track βII-spectrin periodicity in regenerating axons from mouse cortical neurons, human iPSC-derived cortical neurons, and human iPSC-derived motor neurons following mechanical axotomy. Regrowing axons initially lack periodic βII-spectrin organization, particularly near the growth cone. Over 8 to 15 days, periodicity is partially restored in intermediate axonal regions, while distal segments remain disorganized. We found that reducing Rho kinase ROCK-2 activity either pharmacologically or by CRISPRi promotes axon regrowth and accelerates MPS recovery rate five-fold, reaching near-normal levels by 3 days post-injury. To identify the key effectors, we performed co-immunoprecipitation mass spectrometry of the βII-spectrin complex under injury and ROCK-2-inhibited conditions. Myosin 5A (MYO5A) association with spectrin rose sharply upon injury and further increased when ROCK-2 was absent. Functional experiments positioned MYO5A downstream of ROCK-2. Knocking down MYO5A abolished the enhanced regrowth of ROCK-2-deficient neurons, while overexpressing MYO5A increased regrowth in wild-type neurons. Depleting MYO5A also partially disrupted βII-spectrin periodicity in healthy, uninjured axons, indicating a requirement for MYO5A in MPS maintenance under physiological conditions. STED imaging demonstrated that ROCK-2 is arranged periodically along the axon at 190 nm intervals, suggesting it regulates the local lattice. These findings define a ROCK-2/MYO5A pathway linking a targetable kinase to nanoscale cytoskeletal repair and axon regeneration.

## INTRODUCTION

The application of super-resolution fluorescence microscopy to neuronal cell biology uncovered a previously unexpected level of subcortical cytoskeletal organization. Using stochastic optical reconstruction microscopy (STORM), Xu, Zhong, and Zhuang identified the membrane-associated periodic skeleton (MPS). This submembrane lattice consists of short actin filaments organized into circumferential ring-like structures spaced at a characteristic ∼190 nm periodicity, bridged by αII/βII-spectrin tetramers and capped by adducin ^1^. Further studies employing STORM and stimulated emission depletion (STED) super-resolution microscopy revealed that this periodic lattice is not limited to a single neuronal subtype. Instead, it exists in excitatory granule cells, inhibitory parvalbumin-positive interneurons, dopaminergic neurons, motor neurons, and dorsal root ganglion sensory neurons^2,3^. The MPS is also phylogenetically conserved, appearing in species such as *Caenorhabditis elegans*, *Drosophila melanogaster*, and *Gallus gallus*, as well as in mammals such as rodents and humans^2–4^. This level of conservation indicates a fundamental and indispensable function of MPS in neuronal biology.

Since it was first described, the roles of the MPS have grown substantially. The spectrin-dependent skeleton provides mechanical support to the long, thin structure of axons. In *C. elegans* β-spectrin mutants, axons break spontaneously during normal locomotion^5^. The actin-spectrin lattice acts as a tension-buffering shock absorber, safeguarding axons from mechanical stress^6^. In addition to providing structural support, the MPS arranges transmembrane proteins such as ion channels, cell adhesion molecules, and receptor tyrosine kinases into regular, periodic groups. This organization forms signaling platforms that facilitate receptor transactivation and activate downstream ERK signaling pathways^7,8^. It also interacts with non-muscle myosin II to form actomyosin complexes that control axon diameter, affecting action potential conduction velocity and axonal transport efficiency^9,10^. The MPS is involved in the assembly and function of the axon initial segment^11^. Additionally, the MPS affects the stability of axonal microtubules^12^. Co-immunoprecipitation and mass spectrometry studies have uncovered hundreds of potential MPS-interacting proteins, expanding the list of related molecules and their functions^13^.

During neuronal development, the MPS assembles in a proximal-to-distal fashion. In cultured hippocampal neurons, periodic βII-spectrin organization initially appears in the proximal axon as early as 2 days *in vitro* (DIV), then proceeds distally over time, eventually covering the entire axon shaft by about DIV10^14^. This process involves distal transport of spectrin tetramers by kinesin motors^15^ and requires actin nucleation to form new MPS patches, which then coalesce with the existing proximal lattice^16,17^. Once established, the MPS remains stable with a slow turnover of βII-spectrin, as measured by fluorescence recovery after photobleaching^14^. Although progress has been made, the molecular mechanisms that regulate MPS maintenance in mature axons, particularly under pathological conditions, remain poorly understood.

There is growing evidence that MPS remodeling is involved in axon degeneration. MPS disassembly, marked by loss of βII-spectrin periodicity and reduced F-actin, occurs independently of caspase-mediated apoptosis and precedes axonal fragmentation during trophic factor withdrawal in sensory neurons^18,19^. Chemical stabilization of F-actin reduced both MPS loss and axonal fragmentation, indicating that MPS integrity is essential for axonal survival and that its disruption is a necessary step leading to degeneration^18^ and a potential target for neuroprotection. In contrast, the contribution of the MPS to axon regeneration is not well defined.

After injury, surviving neurons can extend new axonal processes from their severed stumps to re-establish connections under certain circumstances. Detailed studies have examined the cytoskeletal dynamics involved, focusing on microtubules, neurofilaments, and the active actin structures present within the growth cone^20^. The status of the submembrane periodic skeleton during regeneration remains unclear, though there are several reasons to believe the MPS is relevant. First, it offers the mechanical support that maintains axonal integrity^5,6^; regrowing axons navigating a hostile tissue environment would presumably benefit from the early reconstitution of this scaffold. Second, the MPS organizes signaling receptors and adhesion molecules that may support growth cone guidance and interactions between axons and their environment^7,8^. Third, the assembly of the MPS during development occurs in a proximal-to-distal pattern, which may be mirrored during the maturation of regrowing axonal segments. Open questions remain regarding the degree to which the MPS is disrupted during axon regeneration, whether it can recover, and whether pharmacological interventions can accelerate its restoration to enhance regeneration.

Rho-associated coiled-coil kinase ROCK-2 is a strong candidate for regulating MPS dynamics during regeneration. ROCK-2 serves as a key downstream effector of RhoA GTPase, phosphorylating substrates that regulate actin–myosin contractility, such as myosin light chain (MLC), LIM kinases (LIMKs), and collapsin response mediator protein 2 (CRMP2)^21^. The RhoA-ROCK pathway is potently activated by brain and spinal cord injury^22,23^ and by myelin-associated neurite growth inhibitors such as Nogo-A, MAG, and OMgp, which signal to growth cone collapse and block regenerative growth^21,24–27^. Pharmacological ROCK inhibition using small molecules such as Y-27632 and Fasudil promotes axon regeneration and functional recovery in multiple rodent spinal cord injury models^26,28–30^. Until now, the understanding of the pro-regenerative effects of ROCK inhibition has mainly focused on growth cone dynamics and the reduction of inhibitory signaling factors^21^. It has not been investigated whether ROCK-2 affects the nanoscale organization of the MPS or its downstream effectors.

Myosin 5A (MYO5A) is an unconventional class V myosin motor protein responsible for transporting various cargoes along actin filaments. It plays important roles in organelle trafficking, membrane dynamics, and neuronal morphogenesis^31,32^. In neurons, MYO5A is essential for shaping dendritic spines, moving synaptic vesicles, and transporting endoplasmic reticulum into dendritic spines^33^. Mutations in MYO5A lead to Griscelli syndrome type 1, which is marked by neurological impairment^34^. However, the role of MYO5A in the formation or dynamics of the spectrin-based periodic skeleton has not been described.

Here, we use STED super-resolution microscopy to demonstrate that the MPS is disrupted in a distance-dependent manner in regenerating axons of mouse cortical, human iPSC-derived cortical, and human iPSC-derived motor neurons after mechanical injury. MPS periodicity is partially restored within 8–15 days after injury, though this recovery is limited and varies spatially in the regrowing axons. Both pharmacological inhibition and CRISPR interference (CRISPRi)-mediated knockdown of ROCK-2 boost overall axon regrowth and significantly speed MPS reassembly in regenerating axons at 3 days post-injury. This is a key time point when vehicle-treated controls exhibit minimal recovery of periodic organization. An unbiased proteomic analysis of the βII-spectrin interactome reveals MYO5A as a downstream effector. Its expression and physical association with the spectrin complex increase after injury and with ROCK-2 inhibition. Epistasis experiments indicate that MYO5A is both necessary and sufficient for axon regeneration. Knocking down MYO5A eliminates the regenerative benefits observed in ROCK-2-deficient neurons, whereas overexpressing it promotes regrowth in wild-type neurons without altering ROCK-2. MYO5A knockdown also disrupts βII-spectrin periodicity in intact, uninjured axons, indicating its essential role in maintaining the MPS. These findings identify a ROCK-2/MYO5A signaling pathway that connects a targetable kinase to nanoscale cytoskeletal organization and regeneration potential, offering insights for enhancing axon repair after nervous system injury.

## RESULTS

### βII-spectrin periodicity is initially lost in regenerating axons of mouse and human neurons

To understand MPS behavior during axon regeneration, we established an *in vitro* scrape injury model using primary mouse cortical neurons and human iPSC-derived cortical neurons and employed stimulated emission depletion (STED) super-resolution microscopy to examine βII-spectrin periodicity in regrowing axons at the nanoscale. We first confirmed that βII-spectrin exhibited robust periodic organization in intact, uninjured axons across multiple neuronal types. In primary mouse cortical cultures at DIV14 and human iPSC-derived cortical neurons at DIV43, extensive axonal networks co-expressing βII-spectrin and βIII-tubulin were observed (Supplementary Fig. 1a, b). To extend these observations to a disease-relevant cell type, we characterized human iPSC-derived motor neurons at DIV32, confirming their identity by co-expression of βIII-tubulin with the motor neuron markers Islet-1, FOXP1, and ChAT (Supplementary Fig. 1c). STED imaging of intact motor neuron axons revealed clear periodic βII-spectrin organization, and autocorrelation analysis confirmed oscillations with a spacing of ∼190 nm, consistent with the canonical MPS periodicity described previously in rodent neurons ^1,7^ (Supplementary Fig. 1d, e). After establishing a baseline MPS organization in intact neurons, we investigated whether the periodic arrangement of βII-spectrin persists in axons that regenerate following mechanical injury. We performed scrape injury at DIV10 (mouse) or DIV40 (human) and fixed cultures 3 days post-injury for STED imaging (Fig. 1a). For intact axonal regions proximal to the cell body, we defined two regions of interest: R1a (∼50 μm from the soma) and R1b (∼100 μm from the soma). In mouse cortical neurons, STED imaging revealed that βII-spectrin retained its characteristic periodic pattern in R1a, whereas this organization appeared modestly reduced in R1b (Fig. 1b). A similar pattern was observed in human iPSC-derived cortical neurons, with robust periodicity in R1a and diminished periodic organization in R1b (Fig. 1c). Fourier transformation analysis confirmed that both R1a and R1b displayed prominent spectral peaks at the frequency corresponding to ∼190 nm periodicity in both mouse and human neurons (Fig. 1d, e). Autocorrelation analysis corroborated these findings, with both R1a and R1b exhibiting strong periodic patterns in the autocorrelation function across both species (Fig. 1f-g).

**Figure 1:**
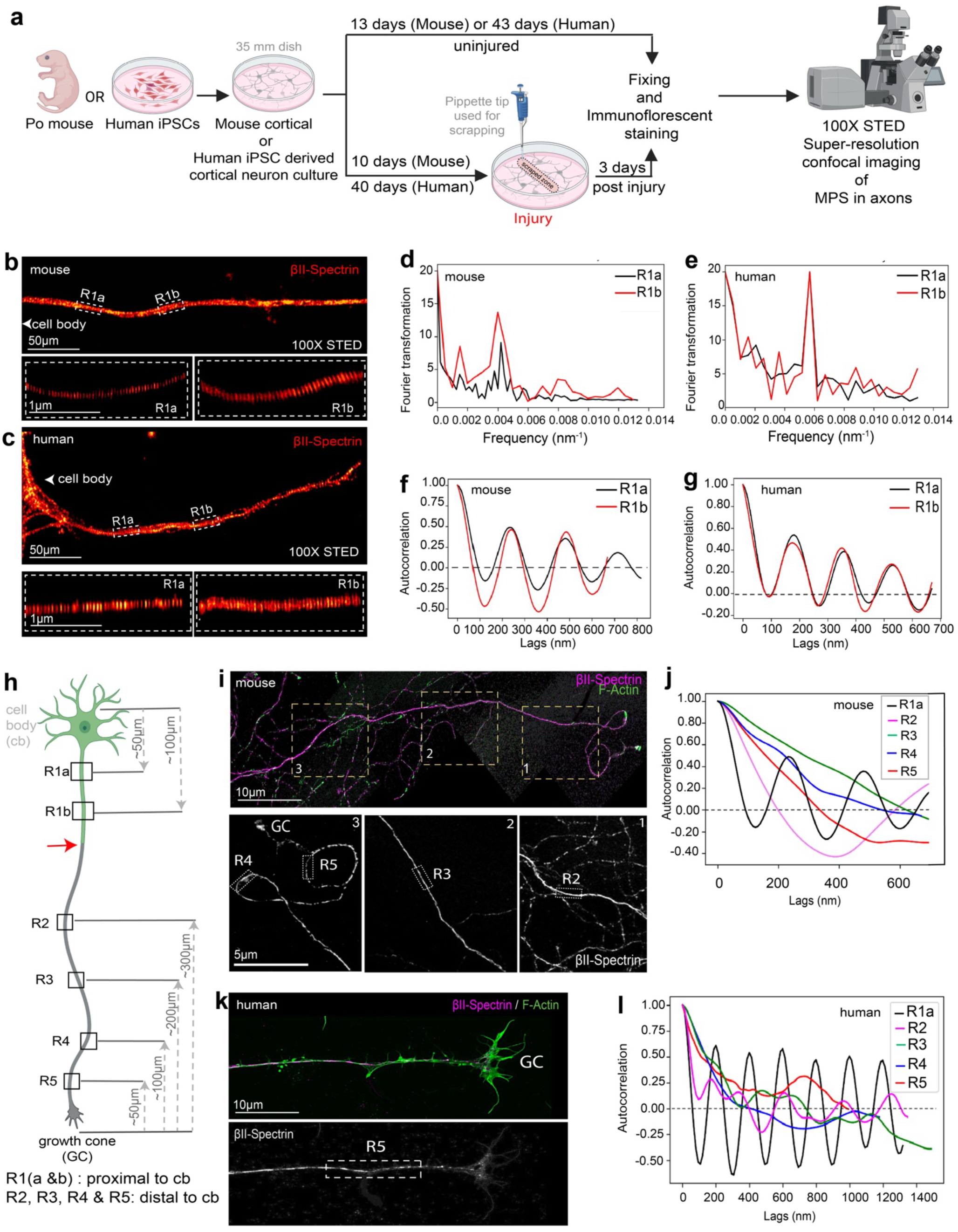
Periodic βII-spectrin arrangement is lost in regenerating mouse and human cortical axons. **a.** Schematic of the experimental workflow. Primary cortical neurons from P0 mice or human iPSC-derived cortical and motor neurons were cultured in 35-mm dishes. Uninjured controls were fixed at 13 days in culture (mouse) or 43 days in culture (human). For injury conditions, scrape injury was performed using a 20 μl pipette tip at DIV10 (mouse) or DIV40 (human). Cultures were fixed at 3 days post-injury, then immunostained, and subsequently imaged by STED super-resolution confocal microscopy at 100X magnification to assess the membrane-associated periodic skeleton (MPS) localization in axons. **b-c**. Representative STED super-resolution image of an axon of a mouse cortical neuron (b) and a human cortical neuron (c). The axon was immunostained for βII-spectrin (Red hot). The top panel shows the intact axon proximal to the cell body, with regions of interest, R1a and R1b, indicated by dashed boxes. Bottom panels show magnified views of R1a and R1b, revealing periodic βII-spectrin organization in both the ROIs. Scale bars: top panel, 10 μm; bottom panels, 1 μm. **d-e**. Fourier transformation analysis plots of βII-spectrin fluorescence intensity profiles along mouse (d) and human (e) axons at regions R1a (black) and R1b (red). Both R1a and R1b display prominent spectral peaks consistent with ∼190 nm periodicity. **f-g**. Autocorrelation analysis plots of βII-spectrin distribution in mouse (f) and human (g) axons. The autocorrelation function (ACF) is plotted as a function of spatial lag (nm). Both R1a (black) and R1b (red) exhibit periodic oscillations characteristic of intact MPS. The dashed lines in the plots indicate zero correlation. **h.** Schematic illustrating the regional sampling strategy used to assess MPS periodicity along regrowing axons. Regions R1a and R1b are located proximal to the cell body (cb), at approximately 50 μm and 100 μm from the soma, respectively. Regions R2–R5 are positioned along the regrowing axonal segment distal to the injury site, at progressively decreasing distances from the growth cone (GC): R2 at approximately 300 μm, R3 at approximately 200 μm, R4 at approximately 100 μm, and R5 at approximately 50 μm from the GC. The red arrow indicates the site of scrape injury. **i.** The top panel is a STED image of a regrowing axon of a mouse cortical neuron co-immunostained for βII-spectrin (magenta) and F-actin (green), with numbered boxes (1,2, 3) corresponding to magnified regions. Bottom panels showed higher-magnification views of axonal segments in regions R2–R5 and the growth cone (GC), highlighting the βII-spectrin (Grey) signal. The periodic organization of βII-spectrin is progressively lost toward the growth cone. Scale bars: top panels 10 μm; bottom panels 5 μm. **j.** Autocorrelation analysis plot of βII-spectrin periodicity at regions R1a (black), R2 (pink), R3 (green), R4 (blue), and R5 (red) along a single regrowing axon of an injured mouse cortical neuron. Periodicity is strongest in R1a (proximal, intact) and progressively diminishes toward the growth cone (R5). The dashed line indicates zero autocorrelation. **k.** Top panel is a STED image of a regrowing axon of a human iPSC-derived cortical neuron co-immunostained for βII-spectrin (magenta) and F-actin (green), with the growth cone (GC) indicated. The bottom panels showed a magnified view of the βII-spectrin signal in region R5 near the growth cone. Scale bar, 5 μm. **l.** Autocorrelation analysis plot of βII-spectrin periodicity at regions R1a (black), R2 (pink), R3 (green), R4 (orange), and R5 (blue) along a single regrowing axon of an injured human cortical neuron. Consistent with the mouse data, MPS periodicity is most prominent in the proximal intact region (R1a) and is progressively lost in distal regions approaching the growth cone. The dashed line indicates zero autocorrelation.

We then examined the MPS along the full length of regrowing axons extending from the injury site into the scraped zone. Confocal imaging of both mouse and human cortical neurons following injury confirmed that regrowing axons co-expressed βII-spectrin and F-actin, with growth cones enriched in F-actin and βII-spectrin distributing along the axon shaft (Supplementary Fig. 2a–c). To systematically assess MPS periodicity as a function of distance from the growth cone, we defined a series of regions along regrowing axons: R1a and R1b proximal to the cell body, and R2–R5 at progressively decreasing distances from the growth cone (300, 200, 100 and 50 μm, respectively) (Fig. 1h). STED imaging of regrowing mouse cortical neuron axons co-stained for βII-spectrin and F-actin revealed a striking proximal-to-distal gradient. The patterned arrangement of βII-spectrin seen in the proximal intact areas disappeared in the more distal segments near the growth cone (Fig. 1i). Autocorrelation analysis across regions R1a–R5 confirmed this gradient quantitatively, with R1a exhibiting the highest periodicity and R5 (nearest the growth cone) showing essentially no periodic oscillations (Fig. 1j).

This spatial pattern of MPS disruption was conserved in human iPSC-derived cortical neurons. STED imaging of regrowing human axons similarly showed robust βII-spectrin periodicity in proximal regions that was progressively lost toward the distal growth cone (Fig. 1k). Autocorrelation analysis across the same regional framework demonstrated that R1a retained the highest MPS density, while periodicity declined through R2–R4 and was largely absent at R5 (Fig. 1l). A similar proximal-to-distal reduction in periodicity was seen in regenerating human iPSC-derived motor neuron axons at DIV32 after injury. Autocorrelation analysis across regions R1–R4 showed that the middle region (R3) had the most recovery of periodic oscillations, while areas closest to the growth cone stayed mostly aperiodic or non-periodic (Supplementary Fig. 2d, e).

Together, these data indicate that the characteristic periodic pattern of βII-spectrin, crucial for the mature MPS, is largely absent from regenerating axons following mechanical injury. This disturbance depends on the distance, with the most distal axonal segments near the growth cone showing a significant loss of periodicity. This pattern is consistent across species, including mouse cortical neurons and human iPSC-derived cortical and motor neurons, implying that MPS disassembly and incomplete reassembly are intrinsic components of the regenerative process, distinct from its known role in axon degeneration. These findings raise questions about whether and to what extent MPS periodicity can regenerate after longer post-injury intervals, and whether pharmacological or genetic treatments can accelerate this process to support functional axon regrowth.

### βII-spectrin periodicity in regenerating axons recovers slowly and partially

We considered whether the absence of βII-spectrin periodicity from regrowing axons at 3 days post-injury (dpi) is permanent or whether the MPS can recover over longer post-injury intervals. During normal axon development, MPS assembly follows a proximal-to-distal gradient, with periodic organization first appearing near the cell body and progressively extending toward the axon tip over the course of days to weeks^1,14^. To quantify MPS or βII-spectrin periodic localization recovery, we used the amplitude of the autocorrelation function (ACF) of βII-spectrin fluorescence intensity profiles as a metric of MPS or βII-spectrin density. In an intact axon with well-organized MPS, the autocorrelation function displays prominent periodic oscillations, and the amplitude of the first peak at ∼190 nm lag (typically ∼0.9 in mature axons) provides a robust single-value readout of periodic organization (Fig. 2a). We defined the ACF calculation range as the window between 190 nm and 400 nm range denoting the autocorrelation function equals to 0 indicating the baseline of no periodic correlation (Fig. 2b). This metric allowed us to compare MPS density across multiple axonal regions, time points, and neuronal types in a standardized manner. In mouse cortical neurons at 3 days post-injury (dpi), a substantial proportion of distal axonal segments (R3–R5) lacked periodic peaks within the expected range, confirming the widespread loss of MPS organization observed in our initial analysis. By 8 dpi, however, the fraction of segments exhibiting periodicity within the 190–400 nm range increased appreciably, particularly at regions R2 and R3 (Supplementary Fig. 3a). A similar temporal trend was observed in human iPSC-derived cortical neurons: the proportion of axonal segments displaying MPS-range periodicity increased between 3 dpi and 15 dpi, most notably at R2 and R3 (Supplementary Fig. 3b). These categorical analyses provided an initial indication that MPS periodicity partially recovers in regrowing axons over time.

**Fig. 2:**
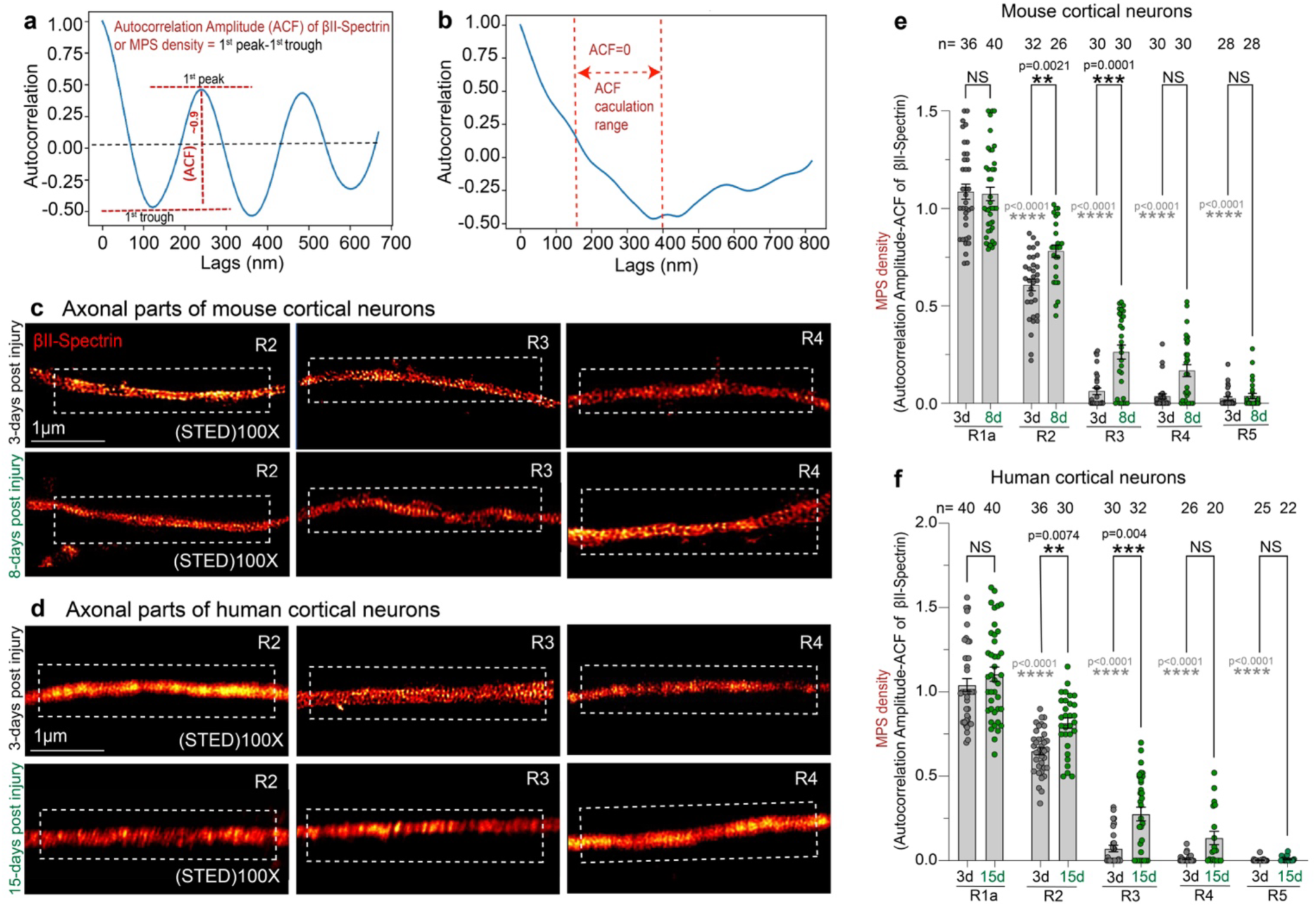
βII-spectrin periodicity recovers partially in regenerating mouse and human axons at 8–15 days post-injury. **a.** Representative autocorrelation functional plot of βII-spectrin fluorescence intensity along an intact human axon, illustrating the derivation of the autocorrelation amplitude (ACF) as a quantitative measure of MPS density. The ACF is calculated as the difference between the first peak (at ∼250 nm lag, corresponding to the fundamental MPS periodicity; red dashed line, ∼0.50) and the first trough (at ∼120 nm lag, corresponding to the half-period between adjacent spectrin positions; red dashed line, ∼ −0.50), yielding an ACF value of ∼0.9 in this example of a well-organized intact axon (red vertical dashed line). This value serves as a single-number readout of the MPS periodic organization. The black dashed line denotes zero autocorrelation. **b.** Representative autocorrelation functional plot illustrating the ACF calculation range. The red dashed lines indicate the two zero-crossing points of the ACF, defining the ACF calculation range used to quantify the degree of periodicity. ACF = 0 indicates the baseline of no periodic correlation. **c.** Representative STED super-resolution images of βII-spectrin (Red hot) along regrowing axons of mouse cortical neurons at regions R2, R3, and R4 (From Fig 1h) at 3 days post-injury 3 days post injury (top row) and 8 dpi (bottom row). At 3 dpi, periodic organization is largely absent across all distal regions. By 8 dpi, periodic βII-spectrin arrangement becomes more apparent, particularly at R2 and R3, indicating partial MPS recovery. Scale bar, 1 μm. **d.** Representative STED super-resolution images of βII-spectrin (Red hot) along regrowing axons of human iPSC-derived cortical neurons at regions R2, R3, and R4 at 3 dpi (top row) and 15 dpi (bottom row). Consistent with findings in mouse neurons, periodic βII-spectrin organization is disrupted at 3 dpi and partially recovers by 15 dpi, most notably at R2 and R3. Scale bar, 1 μm. **e.** Quantification of MPS density (autocorrelation amplitude of βII-spectrin) across five regrown axonal regions (R1a, R2, R3, R4, R5) in mouse cortical neurons at 3 days (3d, black dots) and 8 days (8d, green dots) post-injury. Sample sizes (number of axonal segments, n): R1a (n = 36, 40), R2 (n = 32, 26), R3 (n = 30, 30), R4 (n = 30, 30), R5 (n = 28, 28). MPS density significantly increased between 3d and 8d at R2 (P = 0.0021; black asterisks) and R3 (P = 0.0001; black asterisks), but not at R1a, R4 and R5. Grey asterisks denote comparisons of each region (R2–R5) against the intact region R1a at 3 days post-injury; all regions showed a significant difference, *p* < 0.0001. **f.** Quantification of MPS density across five regrown axonal regions in human cortical neurons at 3 days (3d, black dots) and 15 days (15d, green dots) post-injury. Sample sizes or n: R1a (n = 40, 40), R2 (n = 36, 30), R3 (n = 30, 32), R4 (n = 26, 20), R5 (n = 25, 22). MPS density significantly increased between 3d and 15d at R2 (*P* = 0.0074) and R3 (*P* < 0.0001), but not at R1a, R4, or R5. Grey asterisks denote comparisons of each region (R2–R5) against the intact region R1a at 3 days post-injury; all regions showed a significant difference, *p* < 0.0001. In both **e** and **f**, data are presented as mean ± SEM with individual data points shown. Statistics: One-way ANOVA with Tukey’s post hoc test for multiple comparisons. NS, not significant; \*\**P* < 0.01; \*\*\**P* < 0.001; **** i < 0.0001. P-values are indicated on the graphs.

To visualize this recovery, we performed STED super-resolution imaging of βII-spectrin in regrowing axons of mouse cortical neurons at 3 dpi and 8 dpi. At 3 dpi, the βII-spectrin signal at regions R2, R3, and R4 was largely continuous and lacked periodic organization (Fig. 2c, top row). By 8 dpi, however, the same regions displayed a markedly different appearance at R2 and R3; periodic ring structures of βII-spectrin became partially visible, indicating substantial re-establishment of the MPS lattice. At R4, periodic features of βII-spectrin remained consistent with 3dpi (Fig. 2c, bottom row). We observed a similar pattern in human iPSC-derived cortical neurons, although over a longer time period. At 3 dpi, βII-spectrin at regions R2, R3, and R4 was largely aperiodic (Fig. 2d, top row). By 15 dpi, periodic organization became evident at R2 and R3, where the characteristic ∼190 nm spacing of βII-spectrin rings was partially restored. At R4, some periodicity was observed, but the pattern remained less well-defined (Fig. 2d, bottom row). The longer recovery time in human cortical neurons (15 dpi versus 8 dpi in mice) aligns with the slower maturation of human iPSC-derived neurons compared with primary rodent cultures^35^. Quantification of MPS density (ACF of βII-spectrin) across all sampled regions confirmed these visual observations (Fig. 2e and f). In mouse cortical neurons, the density of MPS at the proximal intact region R1a remained constant from 3 dpi to 8 dpi, indicating that the recovery process did not impact the proximal axon. In contrast, MPS density at R2 increased significantly by 8 dpi (*P* < 0.01), and the recovery at R3 was even more pronounced (*P* < 0.001). R4 and R5, the regions nearest the growth cone, showed no statistically significant improvement in MPS density by 8 dpi, indicating that recovery at this time point was restricted to intermediate axonal regions (Fig. 2e). In human iPSC-derived cortical neurons, the same regional pattern of recovery was observed over the 3-to-15 dpi interval (Fig. 2f). The R1a periodicity stayed consistent, confirming the proximal axon’s structural integrity at both time points. MPS density at R2 and R3 increased significantly by 15 dpi (*P* < 0.01 for both), whereas R4 and R5 did not show significant improvement. In both mouse and human cortical neurons, MPS recovery was primarily observed in intermediate axonal segments (R2 and R3), located about 200–300 μm from the growth cone. In contrast, the most distal segments (R4 and R5) remained largely aperiodic even at the latest time points examined.

We then examined whether this partial MPS recovery also occurs in human iPSC-derived motor neurons, a cell type of particular significance for peripheral nerve damage^36,37^ and amyotrophic lateral sclerosis^38^. The classification of autocorrelation peaks showed a progressive increase in the percentage of motor neuron axonal segments exhibiting periodicity in the 190–400 nm range from 3 dpi to 8 dpi, with the largest increases observed at R1–R4 (Supplementary Fig. 3c). R5, the most distal region, remained largely aperiodic at both time points. STED imaging at region R3 confirmed the emergence of periodic βII-spectrin organization by 8 dpi, with clearly resolved periodic rings visible along the motor neuron axon (Supplementary Fig. 3d). Quantification of ACF across R1–R5 demonstrated significant recovery at R1 (*P* < 0.05) and R3 (*P* < 0.001) by 8 dpi (Supplementary Fig. 3e). R2 showed a trend toward increased periodicity that did not reach statistical significance, while R4 and R5 remained near-zero at both time points. These findings demonstrate that partial MPS recovery is a common characteristic of axon regeneration in both cortical and motor neuron subtypes. The spatial pattern of this recovery is primarily observed in the intermediate regions R2 and R3, whereas the most distal segments remain aperiodic. This matches a model in which MPS assembly occurs behind the moving growth cone, progressing from the proximal to the distal end. This process mirrors the developmental assembly seen in maturing axons^1,14,16^. During axonal development, spectrin tetramers are transported distally and assembled into the periodic lattice over several days, ultimately reaching the axon tip^14,15^. Our data suggest that regrowing axons go through a similar maturation process, but it remains incomplete at 8–15 dpi, with the most distal segments lacking a periodic spectrin lattice. The partial and region-specific MPS recovery suggests there are inherent constraints limiting the speed at which the MPS can be reformed during axon regeneration^12,17^. Furthermore, there is an implication that cytoskeletal regulators might accelerate MPS restoration and support axon regrowth.

### Loss of ROCK-2 function supports rapid recovery of βII-spectrin periodicity

We considered molecular interventions that might accelerate MPS recovery. Rho-associated coiled-coil kinase-2 (ROCK-2) is a major effector in the RhoA signaling pathway. It is a recognized regulator of actin–myosin contractility, growth cone behavior, and neurite extension^21,26–28^. ROCK-2 phosphorylates various substrates, including myosin light chain (MLC), LIM kinases, and collapsin response mediator protein 2 (CRMP2), all of which regulate actin filament dynamics and cause growth cone collapse^21^. In rodent models of spinal cord injury, inhibiting ROCK pharmacologically has been shown to promote axon regeneration^26–29^. However, it remains unclear whether inhibiting ROCK-2 affects the nanoscale organization of the MPS during regeneration. We therefore investigated the effects of both pharmacological ROCK-2 inhibition and CRISPR-mediated ROCK-2 knockdown on axon regrowth and MPS reassembly.

To assess the effect of ROCK-2 inhibition on axon regrowth, we cultured primary mouse (P0) cortical neurons or human iPSC-derived cortical neurons in 96-well plates, then performed mechanical scrape injury at DIV11 (mouse) or DIV40 (human). Next, we administered the ROCK-2 inhibitor Y-27632, Fasudil Hydrochloride, or vehicle (0.1% DMSO) 6 hours post-injury (Supplementary Fig. 4a). The neuronal cultures were fixed 3 days post-injury (3 dpi), immunostained for βIII-tubulin, and imaged by confocal microscopy at 10X magnification. First, we performed a dose response analysis of Y-27632 in mouse cortical neurons. Representative confocal images revealed a visible increase in neurite density within the scraped zone at 5 and 10 μM Y-27632 compared to vehicle (Supplementary Fig. 4b). Quantifying the percentage area of regrowth across concentrations from 0 to 40 μM showed a dose-dependent increase, with significant rises at 5, 10, and 20 μM (*P* < 0.001). However, there was a decline at 40 μM, indicating a biphasic response that may involve toxicity at the highest dose (Supplementary Fig. 4c). Based on this dose-response profile, we chose 10 μM Y-27632 for all subsequent experiments. Next, we examined the time course of regrowth under treatment with 10 μM Y-27632 and 10 μM Fasudil Hydrochloride. Comparison of vehicle and Y-27632- or Fasudil Hydrochloride-treated mouse cortical neurons at 3, 5, and 8 dpi showed that ROCK-2 inhibition by both drugs significantly promoted axon regrowth from 3 dpi through 8 dpi (*P* < 0.001 for both; Supplementary Fig. 4d).

To determine whether these effects apply to human neurons, we treated injured human iPSC-derived cortical neurons with 10 μM Y-27632 or a vehicle control and evaluated regrowth at 3 days post-injury. Confocal imaging revealed a significant increase in neurite extension into the scraped area in Y-27632-treated cultures compared to controls, as confirmed by thresholded image analysis (Supplementary Fig. 4e). Quantitative analysis revealed a significant rise in the percentage of regrowth area (*P* < 0.0001; Supplementary Fig. 4f), indicating that pharmacological inhibition of ROCK-2 promotes axon regrowth in human cortical neurons like mouse neurons. Further, we extended this analysis to human iPSC-derived motor neurons. Confocal imaging of motor neuron cultures immunostained for βIII-tubulin, Islet-1 (motor neuron marker), and DAPI showed significantly enhanced axon regrowth in Y-27632-treated cultures compared to vehicle control (0.1% DMSO) (Supplementary Fig. 4g). Quantifying the percentage area of regrowing axons revealed a significant increase with ROCK-2 inhibition (*P* < 0.0001; Supplementary Fig. 4h), confirming that Y-27632’s pro-regenerative effects are consistent across cortical and motor neuron subtypes.

To confirm that ROCK-2 is the primary kinase responsible, and to minimize off-target effects of pharmacological inhibition, we created CRISPRi-mediated ROCK-2 knockdown (KD) human iPSC lines using two different sgRNAs (sgRNA1, L1; sgRNA2, L2). Confocal imaging verified the knockdown efficiency at the protein level by showing reduced ROCK-2 immunofluorescence in ROCK-2-CRISPR-KD sgRNA1 (L1) neurons compared to wild-type (WT-Scramble2-CRISPR) controls, with βII-spectrin levels remaining unchanged (Supplementary Fig. 5a). Quantifying the ROCK-2/βII-spectrin fluorescence intensity ratio revealed a significant decrease in ROCK-2 protein levels in both cell bodies (*P* < 0.001) and axons (*P* < 0.001) of knockdown neurons compared to the WT-Scramble-2-CRISPR control (Supplementary Fig. 5b). Further, quantitative RT-PCR confirmed that both L1 and L2 lines showed significantly lower ROCK-2 mRNA levels compared to the control Wild Type line (*P* < 0.01 and *P* < 0.001, respectively; Supplementary Fig. 5c). WT-Scramble control and two ROCK-2-CRISPR-KD iPSC lines were differentiated into cortical neurons, cultured in 96-well plates, subjected to scrape injury at DIV40, and then fixed and immunostained after 3 days post-injury for axon regrowth analysis (Supplementary Fig. 5d). Confocal imaging of βIII-tubulin revealed significantly enhanced neurite regrowth into the scraped zone in both L1 and L2 knockdown lines compared to WT controls (Supplementary Fig. 5e). Quantification confirmed that both ROCK-2-CRISPR (KD) lines exhibited significantly greater regrowth compared to the WT-Scramble-2 control (*P* < 0.0001 for both), with no significant difference observed between the two knockdown lines. (Supplementary Fig. 5f). These results indicate that genetic suppression of ROCK-2 can enhance axon regrowth, phenocopying the effects observed with pharmacological inhibition.

After demonstrating that inhibiting ROCK-2 alone can enhance overall axon regeneration, we investigated whether it also accelerate MPS reassembly in the regrowing axons. This directly addresses whether the improved cytoskeletal organization accompanies and potentially contributes to the improved regenerative capacity. To evaluate this, we cultured mouse cortical or human iPSC-derived cortical neurons and introduced a scrape injury. We then treated the neuron cultures with 10 μM Y-27632 or vehicle 6 hours post injury and fixed them 3 days later for STED super-resolution imaging (Fig. 3a). STED imaging at region R3, positioned at an intermediate distance from the growth cone of a mouse neuron (Fig. 1h)—where partial MPS recovery was previously seen at later time points (Fig. 2e and Fig. 3b, top panel)—revealed organized periodic βII-spectrin along the axon shaft as early as 3 dpi in Y-27632-treated neurons (Fig. 3b, bottom). Autocorrelation analysis at R3 in WT mouse neurons treated with 10 µM Y-27632 confirmed a periodic organization, with an ACF value of 1.036, indicating a well-structured MPS (Fig. 3c). Quantifying MPS density across regions R1–R5 (Fig. 1h) in Y-27632-treated versus vehicle-treated neurons showed that ROCK-2 inhibition significantly increased βII-spectrin periodicity at R3 (*P* < 0.0001) and R4 (*P* < 0.05) at 3 dpi (Fig. 3d). The proximal region R1a, which already had a significant MPS organization in vehicle-treated controls, was not notably affected by Y-27632 treatment. Additionally, regions R2 and R5 — the farthest and closest from the growth cone, respectively exhibited no changes in MPS organization and were unaffected by Y-27632 treatment (NS, Fig. 3d). This regional pattern indicates that inhibiting ROCK-2 specifically accelerates MPS recovery at intermediate axonal distances, where the periodic lattice is actively assembling. It does not alter MPS organization in pre-existing proximal segments or in the most immature distal segments. To validate these results using a genetic approach, we employed ROCK-2-CRISPR-knockdown iPSC lines (L1 and L2) that were introduced into human cortical neurons. After 40 days in culture, a scrape injury was induced, and cultures were fixed at 3 days post-injury (dpi) for STED imaging (Fig. 3e). STED confocal imaging of βII-spectrin in the R3 region of neurons with ROCK-2-CRISPR (KD) sgRNA1 and sgRNA2 (L1 and L2) at 3 dpi revealed regenerating axons with clearly defined periodic βII-spectrin organization (Fig. 3f, middle and lower panels), in contrast to the WT-Scramble-2 control (Fig. 3f, top panel). Quantification of MPS density in regions R1–R5 of WT, L1, and L2 neurons at 3 dpi revealed that both ROCK-2 knockdown lines had significantly higher βII-spectrin periodicity compared to WT (Fig. 3g). The most prominent effects were seen at R3, where both L1 and L2 showed significantly higher MPS density compared to WT (*P* < 0.0001). Significant enhancements were also detected at R2 (*P* < 0.001 and 0.003) and at R4 (*P* < 0.05), but not at R5. The wider regional effect of genetic knockdown, as opposed to acute pharmacological inhibition, probably resulted from the sustained, constitutive reduction of ROCK-2 during neuronal development and throughout the post-injury period in the CRISPR lines.

**Fig. 3:**
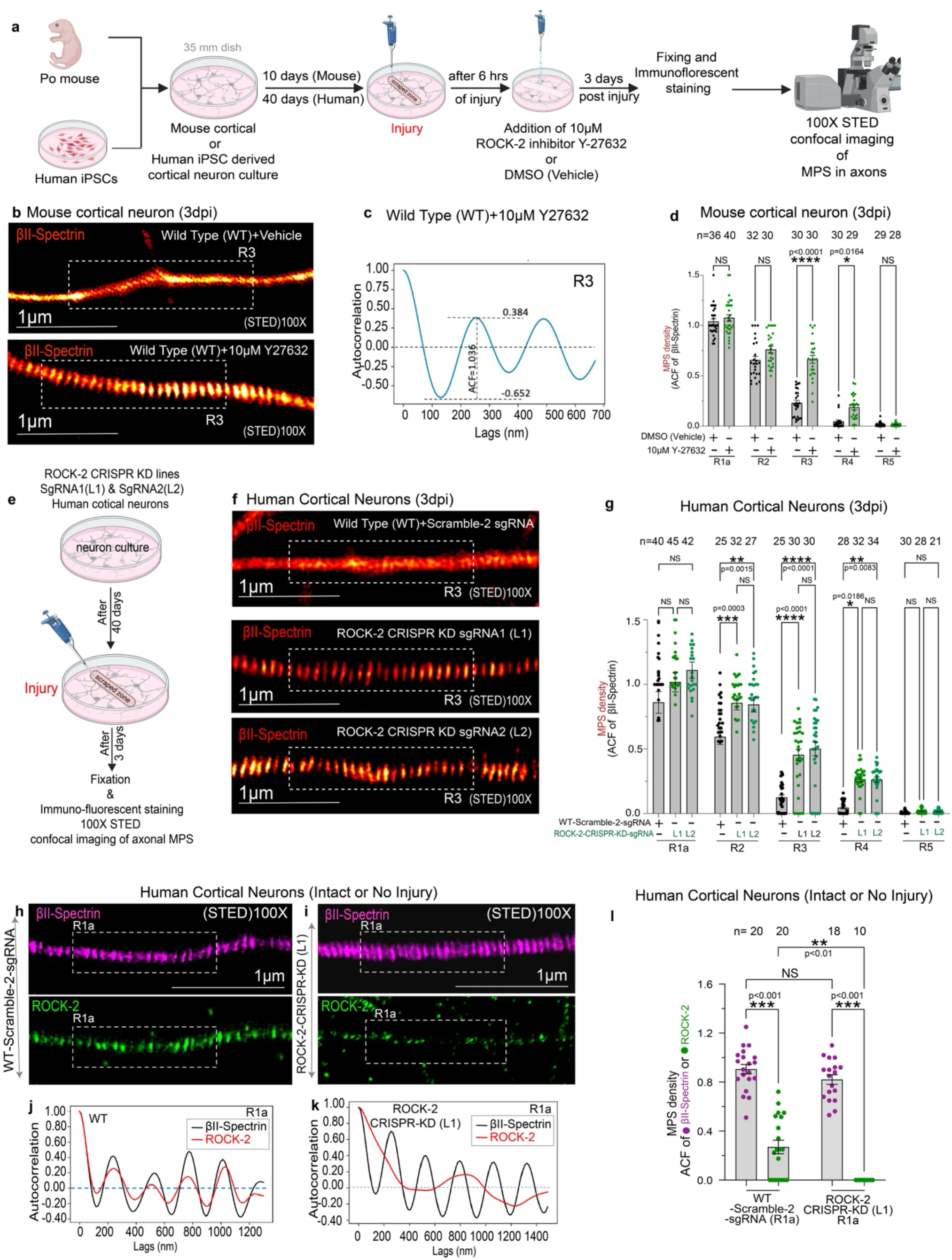
Pharmacological inhibition and CRISPRi-mediated knockdown of ROCK-2 accelerate MPS reassembly in regenerating axons. **a.** The schematic of the experimental workflow for pharmacological inhibition of ROCK-2 (RHO- associated protein kinase 2) in both mouse and human cortical neurons. Primary cortical neurons from P0 mice or human iPSC-derived cortical neurons were cultured in 35-mm dishes. Mechanical scrape injury was performed at DIV10 (mouse) or DIV40 (human). Six hours post-injury, cultures were treated with 10 μM Y-27632, a ROCK-2 inhibitor, or vehicle (0.1% DMSO). Cultures were fixed 3 days post-injury, immunostained, and imaged by STED super-resolution confocal microscopy at 100X magnification to assess MPS organization in axons. **b.** Representative 100X magnification of STED super-resolution images of βII-spectrin (Red hot) at axonal region R3 in regrowing mouse cortical neuron axons at 3 days post-injury (3 dpi). Top: Wild-type (WT) neurons treated with vehicle (DMSO), showing largely aperiodic βII-spectrin signal with no clearly discernible periodic organization at R3. Bottom: Wild-type neurons treated with 10 μM Y-27632 (ROCK-2 inhibitor), displaying markedly enhanced periodic βII-spectrin organization at R3, with clearly resolved periodic puncta spaced at ∼190 nm intervals along the regrowing axon shaft. Dashed boxes indicate representative segments used for autocorrelation analysis. ROCK-2 inhibition accelerates the recovery of MPS periodicity at intermediate axonal regions as early as 3 dpi. Scale bars, 1 μm. **c.** Representative autocorrelation function of βII-spectrin fluorescence intensity at region R3 of a regrowing axon. The ACF is calculated as the difference between the first peak (0.384) and the first trough (−0.652), yielding 1.036. Dashed lines indicate measurement parameters. **d.** Quantification of MPS density (ACF of βII-spectrin) across five axonal regions (R1a, R2, R3, R4, R5) in mouse cortical neurons at 3 days post-injury, treated with vehicle (−, black dots) or 10 μM Y-27632 (+, green dots). Sample sizes or n (number of axonal segments): R1a (*n* = 36, 40), R2 (*n* = 32, 30), R3 (*n* = 30, 30), R4 (*n* = 30, 29), R5 (*n* = 29, 28). Y-27632 treatment significantly increased MPS density at R3 (*P* < 0.0001) and R4 (*P* = 0.0164) only at 3 days post-injury. **e.** A schematic of the experimental workflow for CRISPR-mediated ROCK-2 gene interference knockdown. Human iPSC lines with ROCK-2 knockdown via two independent sgRNAs (sgRNA1, L1; sgRNA2, L2) were differentiated into cortical neurons, cultured for 40 days, subjected to injury, and fixed 3 days post-injury for STED imaging of axonal MPS. **f.** Representative STED super-resolution images of βII-spectrin (red hot) at axonal region R3 in regrowing human iPSC-derived cortical neuron axons at 3 days post-injury (3 dpi). Top: wild-type (WT) neurons expressing Scramble-2 sgRNA, showing largely aperiodic βII-spectrin signal at R3, consistent with incomplete MPS reassembly at this early time point. Middle: ROCK-2 CRISPR KD sgRNA1 (L1) neurons, displaying markedly enhanced periodic βII-spectrin organization at R3, with clearly resolved periodic puncta along the regrowing axon shaft. Bottom: ROCK-2 CRISPR KD sgRNA2 (L2) neurons, showing a similar enhancement of βII-spectrin periodicity at R3, independently confirming the effect observed with sgRNA1. Dashed boxes indicate representative segments used for autocorrelation analysis. Scale bars, 1 μm. **g.** Quantification of MPS density across five axonal regions in human cortical neurons from wild-type (WT, black dots), ROCK-2 knockdown sgRNA1 (L1, green dots), and sgRNA2 (L2, green dots) lines at 3 days post-injury. Sample sizes: R1a (*n* = 40, 45, 42), R2 (*n* = 25, 32, 27), R3 (*n* = 25, 30, 30), R4 (*n* = 28, 32, 34), R5 (*n* = 30, 28, 21). ROCK-2 knockdown significantly increased MPS density at R2 (L1: *P* = 0.0003; L2: *P*< 0.0001), R3 (L1: *P* < 0.0001; L2: *P* = 0.0015), and R4 (L1: *P* = 0.0186; L2: NS), with L1 also showing a significant increase at R4 (*P* = 0.0083 vs WT). No significant changes were observed at R1a or R5. In **d** and **g**, data are presented as mean ± SEM with individual data points shown. Statistics: One-way ANOVA with Tukey’s post hoc test for multiple comparisons. NS, not significant; \*\**P* < 0.01; \*\*\**P* < 0.001; ****P < 0.0001. P-values are indicated on the graphs. N = 3 independent experiments. **h-i.** STED super-resolution images of βII-spectrin (magenta) and ROCK-2 (green) at region R1a in intact, uninjured axons of wild-type (WT + Scramble-2 sgRNA) (**h**) and ROCK-2-CRISPR-KD (L1) (**i**) human iPSC-derived cortical neurons. In WT axons, both βII-spectrin and ROCK-2 display periodic organization. In ROCK-2-KD axons, βII-spectrin periodicity is maintained while ROCK-2 signal is reduced and aperiodic, confirming specificity. Dashed boxes indicate segments used for autocorrelation analysis. Scale bars, 1 μm. **j-k.** Autocorrelation analysis of βII-spectrin (black) and ROCK-2 (red) intensity profiles at R1a in WT (**j**) and ROCK-2-CRISPR-KD (L1) (**k**) neurons. ROCK-2 exhibits ∼190 nm periodicity in WT axons but not in KD axons, whereas βII-spectrin periodicity is preserved in both genotypes. Dashed lines indicate a zero-correlation baseline. **l.** MPS density (ACF) of βII-spectrin (magenta) and ROCK-2 (green) at R1a in intact WT and ROCK-2-KD (L1) axons. ROCK-2 periodicity is significantly reduced in KD neurons (P < 0.001), while βII-spectrin periodicity is unaffected (NS). Data are mean ± SEM with individual data points; *N* = 3 independent replicates. One-way ANOVA with Tukey’s *post hoc* test. NS, not significant; \*\**P* < 0.01; \*\*\**P* < 0.001.

To provide independent genetic validation that ROCK-2 constrains axon regrowth, we extended our analysis to ROCK-2 knockout (ROCK-2−/−) mice. Primary cortical neurons were isolated from individual P0 pups of ROCK-2(−/−) homozygous mutant or wild-type (WT) littermates and cultured in 35-mm dishes with one pup per plate, while tail biopsies were collected for genotyping (Supplementary Fig. 6a and 6b). A mechanical scrape injury was performed at DIV11, and the cultures were fixed three days afterward (Supplementary Fig. 6a). Confocal imaging of βIII-tubulin immunostained scraped cultures revealed significantly increased neurite regrowth into the injury zone in ROCK-2−/− neurons compared to WT controls (Supplementary Fig. 6c). Quantification showed that genetic deletion of ROCK-2 in mice resulted in a significant rise in the percentage of regrowth area (*P* < 0.01, two-tailed Student’s t-test; Supplementary Fig. 6 d). These findings, validated by pharmacological and genetic data from three approaches, identify ROCK-2 as the key negative regulator of overall axon regrowth and post-injury MPS reformation.

The specific function of ROCK-2 in regulating MPS dynamics raised the question of whether ROCK-2 is spatially associated with the periodic skeleton. To address this, we used dual-color confocal and STED super-resolution imaging on control (WT) human iPSC-derived cortical neuron axons co-immunostained for βII-spectrin and ROCK-2. Confocal images of wild-type neurons revealed that both βII-spectrin and ROCK-2 are in neuron cell bodies and axons (Supplementary Fig. 7a and b). From the merged images, we calculated Pearson’s correlation coefficient to evaluate ROCK-2/βII-spectrin co-localization in both cell bodies and axons (Supplementary Fig. 7c). The Pearson’s coefficient was significantly higher in cell bodies than in axons (*P* < 0.0001, unpaired two-tailed Student’s t-test), indicating that while ROCK-2 co-localizes with βII-spectrin in both regions, the overlap is more pronounced in the soma than in the axon. STED imaging provided a more detailed view of the nanoscale organization of ROCK-2 along intact axons (Fig. 3h–k). In wild-type (WT-Scramble-2 control) neurons, dual-color STED imaging of region R1a revealed that βII-spectrin displayed its typical periodic pattern along the axon shaft, while ROCK-2 also showed a partially periodic pattern (Fig. 3h). ROCK-2 also showed a partially periodic arrangement (Fig. 3h). Autocorrelation analysis of βII-spectrin and ROCK-2 showed periodic oscillations with similar spacing, with peaks approximately 190 nm apart, characteristic of MPS periodicity (Fig. 3j). In ROCK-2-CRISPR-KD (L1) neurons, STED imaging showed that βII-spectrin preserved its periodic pattern at R1a, while the ROCK-2 signal was notably reduced and did not display a periodic distribution (Fig. 3i). Autocorrelation analysis indicated that the periodic pattern of βII-spectrin remained in ROCK-2-KD neurons, while the periodicity in ROCK-2 was lost (Fig. 3k). This suggests that ROCK-2 does not play a role in βII-spectrin spacing in healthy axons. Quantifying the MPS density using the autocorrelation amplitude (ACF) of βII-spectrin and ROCK-2 in intact axons at region R1a (Fig. 3l) confirmed these results. In wild-type neurons, βII-spectrin exhibited high ACF values, and ROCK-2 showed notable periodicity, though with slightly lower amplitude (*P* < 0.001). In ROCK-2-KD (L1) neurons, βII-spectrin periodicity was fully preserved (not significantly different from WT βII-spectrin), while ROCK-2 periodicity was notably decreased (*P* < 0.01 compared with WT ROCK-2; *P* < 0.001 compared with WT βII-spectrin).

To determine whether the improved MPS recovery seen with ROCK-2 inhibition is due to a mechanistic connection between ROCK-2 and the periodic skeleton, or if it is merely a secondary consequence of enhanced axon regrowth, we investigated an alternative pro-regenerative pathway. PTEN deletion is known to promote axon regeneration by activating the PI3K/Akt/mTOR signaling pathway^39,40^. We generated two independent human iPSC lines with PTEN knockdown via CRISPRi (L1 and L2) and assessed axon regrowth and MPS periodicity after scrape injury. Confocal images taken three days post-injury reveal increased axon regrowth into the scraped area in both PTEN-CRISPR-KD lines (L1 and L2) compared to wild-type (WT- Scramble-2 sgRNA) controls, as shown by βIII-tubulin immunostaining and similar in magnitude to ROCK2 inhibition (Supplementary Fig. 8a). Thresholded images of the selected regions confirm that PTEN-KD cultures have a higher density of regrowing axons. Quantifying the percentage of regrowth area shows both PTEN-KD lines exhibit significantly more regrowth than WT (L1, *P* = 0.004; L2, *P* = 0.003), with no significant difference between L1 and L2 (NS; Supplementary Fig. 8b). Although βII-spectrin shows more regrowth, super-resolution imaging of region R3 in regrowing axons at 3 days post-injury revealed no enhancement in the periodic pattern in PTEN-KD neurons compared to WT controls (Supplementary Fig. 8c). Both WT-Scramble-2 sgRNA and PTEN-CRISPR-KD (L1) neurons showed mostly disorganized βII-spectrin signal at R3, indicating that PTEN knockdown did not enhance MPS reassembly despite acceleration of axon regrowth. Quantification of MPS density, assessed via the autocorrelation amplitude of βII-spectrin across five axonal regions (R1a, R2, R3, R4, R5) in WT, PTEN-CRISPR-KD (L1), and PTEN-CRISPR-KD (L2) neurons at 3 days post-injury, validated these results (Supplementary Fig. 8d). At R1a, MPS density was high and comparable across all three genotypes, indicating that the MPS organization remained intact in the proximal uninjured segments. At R2, all genotypes showed a significant reduction in MPS density relative to R1a levels (P < 0.0001 for WT, L1, and L2), but there were no differences among genotypes within R2 (NS). At R3, R4, and R5, MPS density was almost zero in all conditions, with no notable differences among genotypes at any of these distal regions (NS). Thus, PTEN knockdown improves regrowth to a level equal to ROCK-2 inhibition but fails to restore MPS periodicity in regenerating axons. This demonstrates that increased axon regrowth alone is insufficient to accelerate MPS reassembly, ruling out the simple explanation that ROCK2-induced MPS recovery is a byproduct of enhanced regeneration. Instead, rapid MPS recovery following ROCK-2 inhibition reflects ROCK-2 interaction with the spectrin periodic structure. Thus, pharmacological and genetic evidence indicate that ROCK-2 activity constrains both the speed of axon regrowth and the reconstruction of the MPS periodic lattice in regenerating axons.

### βII-spectrin interactome contains MYO5A as an injury- and ROCK-2-regulated effector

After confirming that ROCK-2 inhibition promotes axon regrowth and MPS reassembly, we sought to identify the underlying molecular mechanisms. The MPS is a complex network of macromolecules whose composition and interactions can vary dynamically during axon injury and repair. Therefore, we conducted a comprehensive proteomic analysis of the βII-spectrin interactome under various injury and ROCK-2 inhibition conditions to identify candidate proteins linking ROCK-2 signaling to MPS remodeling. We performed a co-immunoprecipitation (CoIP) mass spectrometry experiment across six conditions, systematically varying the genotype of iPSC-derived human cortical neurons (WT versus ROCK-2-CRISPR-KD), injury status (−/+), and Y-27632 treatment (−/+), as shown in Supplementary Fig. 9a. Cell lysates from each condition (S1–S6) were subjected to CoIP using antibodies against βII-spectrin, αII-spectrin, and IgG (control) as bait. The eluates were separated by SDS-PAGE, and the excised gel fragments were analyzed by mass spectrometry to identify proteins and determine their relative abundance (Supplementary Fig. 9a and Supplementary Excel 1). This method identified a total of 717 proteins that co-immunoprecipitated with the spectrin complex across conditions (Supplementary Data 1). To assess the specificity of the CoIP, we analyzed the overlap of proteins pulled down by βII-spectrin and αII-spectrin as bait proteins, compared to IgG as a control bait across various conditions (Fig. 4a). In each condition, a core set of proteins was co-precipitated by both spectrin baits but not by the IgG control, confirming their specific interaction with the spectrin complex. The number of shared spectrin-specific interactors varied across conditions. The WT uninjured untreated cultures showed 57 shared proteins, whereas injury combined with Y-27632 treatment resulted in 35, and ROCK-2 CRISPRi KD injured cultures yielded 32 (Fig. 4a, panels i–vi). These condition-dependent variations in the spectrin interactome suggest a dynamic remodeling of the MPS-associated protein network caused by injury and ROCK-2 activity. This method identified 121 proteins that co-immunoprecipitated with the spectrin complex across conditions (Supplementary Excel 2). Unsupervised hierarchical clustering of the protein abundance data, presented as a z-score heatmap, revealed distinct proteomic signatures associated with injury, ROCK-2 inhibition, and their combination (Supplementary Fig. 9b and Supplementary Data 2). Notably, WT uninjured samples clustered separately from injured and Y-27632-treated samples. Meanwhile, ROCK-2 CRISPRi KD samples formed their own distinct cluster, suggesting that both injury and ROCK-2 inhibition substantially affect the spectrin-associated proteome. Gene Ontology (GO) functional enrichment analysis of all 121 identified proteins revealed a prominent emphasis on cytoskeletal and actin-related processes (Supplementary Fig. 9c). The most enriched terms included cytoskeleton organization (GO:0007010), cadherin binding (GO:0045296), and the actin cytoskeleton (GO:0015629). Other highly significant terms were actin filament binding (GO:0051015) and actin filament-based process (GO:0030029). Importantly, terms directly associated with the MPS, such as spectrin (GO:0008091), spectrin-associated cytoskeleton (GO:0014731), and actomyosin (GO:0042641), also showed high enrichment. Additionally, categories related to microtubules (GO:0032970) and cytoskeleton-dependent intracellular transport (GO:0030705) were prominently represented (Supplementary Fig. 9c). This enrichment pattern verifies that our CoIP method successfully identified the spectrin-associated cytoskeletal network and its regulators.

**Fig. 4:**
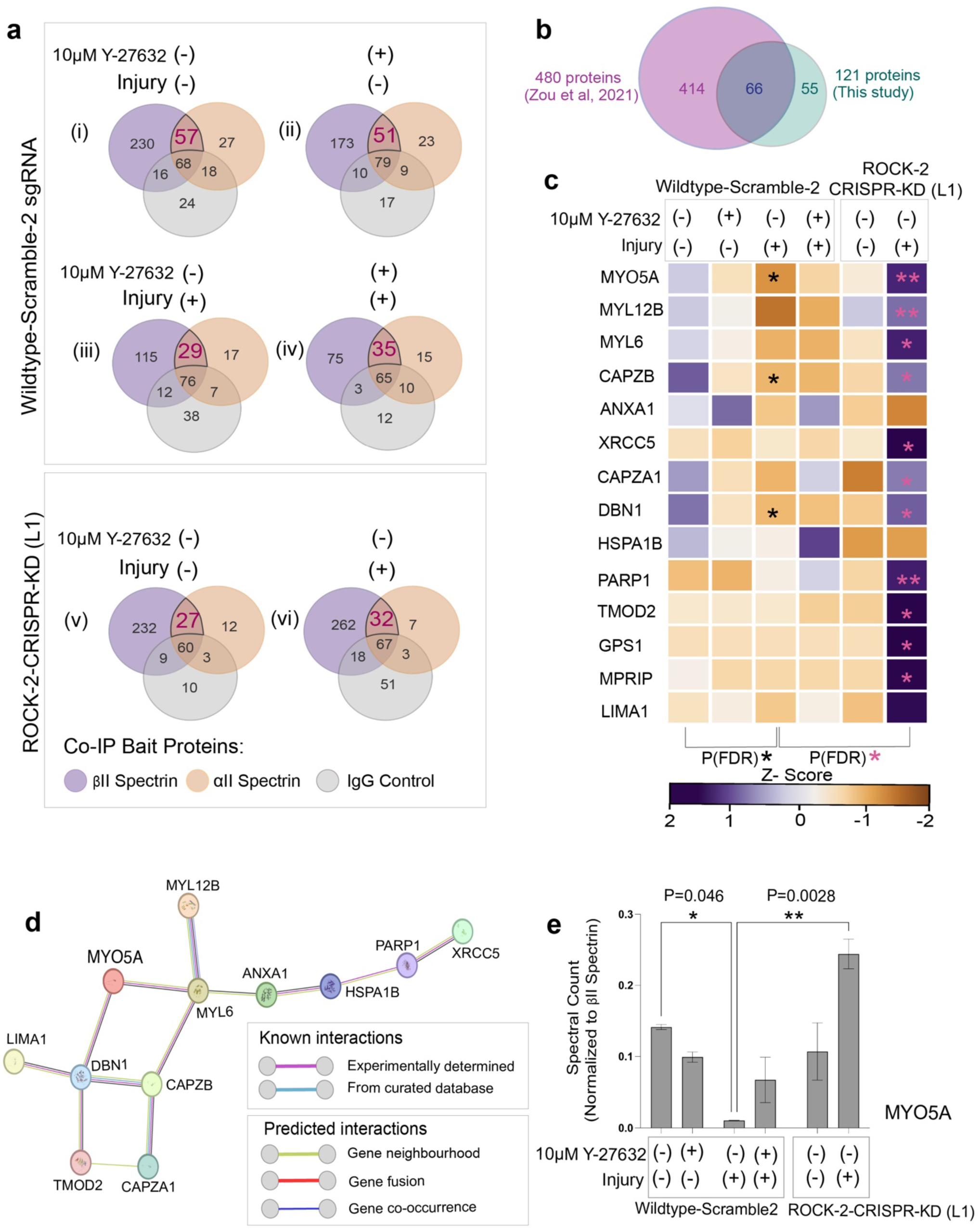
Proteomic mass spectrometry analysis identifies MYO5A as an injury- and ROCK-2-dependent interactor of the βII-spectrin complex. **a.** Venn diagrams showing the overlap of the proteins co-immunoprecipitated with βII-spectrin (purple), αII-spectrin (orange), and IgG control (grey) across six experimental conditions in wildtype neurons (panels i–iv) and ROCK-2 CRISPR knockdown (KD) line 1 neurons (panels v–vi), varying injury status (−, without or +, with) and 10 μM Y-27632 treatment (−, vehicle/+, with drug). Numbers indicate unique and shared protein counts identified per condition. Highlighted numbers (red, underlined) denote proteins co-precipitated by both βII and αII-spectrin baits but not the IgG control. **b.** Venn diagram comparing the βII-spectrin interactome identified in this study (121 proteins, teal) with the previously published spectrin interactome (480 proteins, purple; Zou et al., 2021). An overlap of 87 proteins is indicated between the two studies. **c.** Heatmap of z-score normalized spectral counts for 14 candidate interacting proteins across six experimental conditions (columns) in wildtype and ROCK-2 CRISPR KD neurons. Conditions vary by injury (−/+) and Y-27632 treatment (−/+). Color scale ranges from enriched (dark purple, z = 2) to depleted (dark orange, z = −2). Asterisks indicate statistically significant changes [\**P*(FDR) < 0.05; \*\**P*(FDR) < 0.01] within each genotype group. **d.** STRING protein–protein interaction network of the 14 candidate interactors identified in **c**. **e.** Quantification of MYO5A spectral counts normalized to βII-spectrin spectral counts across injury and Y-27632 treatment conditions in wildtype and ROCK-2 CRISPR KD neurons. Data are presented as mean ± SEM. P values were determined by the two-tailed unpaired *t*-test (*P* = 0.046; \**P* = 0.046, \*\**P* = 0.0028).

To evaluate the novelty and reliability of this interactome, we compared the 121 proteins identified here with a previously published spectrin interactome comprising 480 proteins^13^ (Fig. 4b). This comparison revealed 66 proteins in common, independently confirming many of our identifications. Notably, 55 proteins were unique to our dataset, possibly representing condition-specific interactors that occur only during injury or ROCK-2 inhibition. These were not observed in the previous study, which utilized uninjured mouse hippocampal neurons^13^. Out of the 121 proteins, we identified 14 candidates with the most significant changes in abundance across various conditions (Fig. 4c and Supplementary Data 3). A heatmap illustrating z-score normalized spectral counts for bait proteins across six conditions revealed that several proteins, including MYO5A, MYL12B, MYL6, CAPZB, and DBN1, were significantly upregulated by Y-27632 treatment and/or by CRISPRi of ROCK-2, particularly in injured states. Others, including TMOD2, GPS1, and MPRIP, showed notable changes, especially in the ROCK-2 CRISPR-KD genotype (Fig. 4c). GO functional enrichment analysis of these 14 differentially abundant proteins revealed a strong enrichment in processes related to the actin cytoskeleton, including actin cytoskeleton (GO:0015629), actin filament-based process (GO:0030029), actin cytoskeleton organization (GO:0030036), actin binding (GO:0003779), negative regulation of actin filament depolymerization (GO:0030835), and actin filament capping (GO:0051693) (Supplementary Fig. 9d). STRING in-silico analysis of the protein–protein interaction network for 14 candidate proteins identified a dense hub centered on MYO5A, with known or predicted connections to MYL12B, MYL6, DBN1, CAPZB, CAPZA1, TMOD2, and LIMA1 (Fig. 4d). Several of these interactors are known regulators of actin filament dynamics, including CAPZB and CAPZA1, which function as actin filament capping proteins. TMOD2 is a tropomodulin that caps the pointed end of actin filaments, while DBN1 (Drebin) is an actin-binding protein involved in neuronal cytoskeletal remodeling^41–43^.

The presence of these actin-regulatory proteins near MYO5A in the spectrin interactome suggests that MYO5A may serve as a molecular link between ROCK-2 signaling and MPS-related actin dynamics. Among the 14 candidates, MYO5A itself is an unconventional myosin motor protein involved in organelle transport and actin-based motility^31^. Quantifying MYO5A spectral counts normalized to βII-spectrin across various conditions revealed a striking pattern (Fig. 4e). MYO5A’s association with the spectrin complex was slightly reduced by Y-27632 treatment alone (*P* = 0.046) and was most significantly increased in injured ROCK-2-CRISPR-KD cultures (*P* = 0.0028; Fig. 4e). This pattern shows that MYO5A is downregulated after injury but increases further when ROCK-2 is inhibited in injured conditions. This implies that MYO5A may serve as a downstream effector, with its recruitment to the spectrin complex normally suppressed by ROCK-2 activity and released upon ROCK-2 inhibition. Together, these proteomic results demonstrate that injury and ROCK-2 inhibition significantly alter the βII-spectrin interactome, especially affecting actin cytoskeletal regulators. Among 14 interactors with differential regulation, MYO5A was the most prominently and consistently associated with the spectrin complex, with its association strongly enhanced by ROCK-2 inhibition in the injured condition.

### MYO5A protein expression is upregulated by injury and ROCK-2 inhibition

Our proteomic analysis revealed MYO5A as a strongly regulated interactor of the βII-spectrin complex in response to injury and ROCK-2. MYO5A is an unconventional class V myosin motor protein that transports diverse cargoes along actin filaments and plays key roles in organelle transport, membrane dynamics, and neuronal morphogenesis^31,33,44^. To validate the proteomic findings and explore MYO5A regulation in detail, we performed immunoblot analyses under different conditions in both human iPSC-derived and mouse cortical neurons. These analyses of total cell lysates from human iPSC-derived cortical neurons confirmed the presence of MYO5A (∼215 kDa), βII-spectrin (∼250 kDa), and βII-actin (used as a loading control) across all tested conditions (Fig. 5a). Quantifying MYO5A band intensity, normalized to βII-spectrin revealed a condition-dependent expression pattern (Fig. 5c). In wild-type (WT) neurons, MYO5A levels were low in uninjured, untreated controls and remained unchanged with Y-27632 treatment alone or injury alone. Notably, the combination of injury and Y-27632 treatment significantly increased MYO5A expression (*P* < 0.0001; Fig. 5c). In ROCK-2 CRISPRi knockdown neurons, MYO5A levels were consistently elevated in both uninjured and injured states, significantly exceeding those in WT uninjured controls (*P* < 0.0001; Fig. 5c). This indicates that ROCK-2 activity normally suppresses MYO5A expression. Blocking this inhibition, whether by drug treatment with injury or genetic means, results in elevated MYO5A levels.

**Fig. 5:**
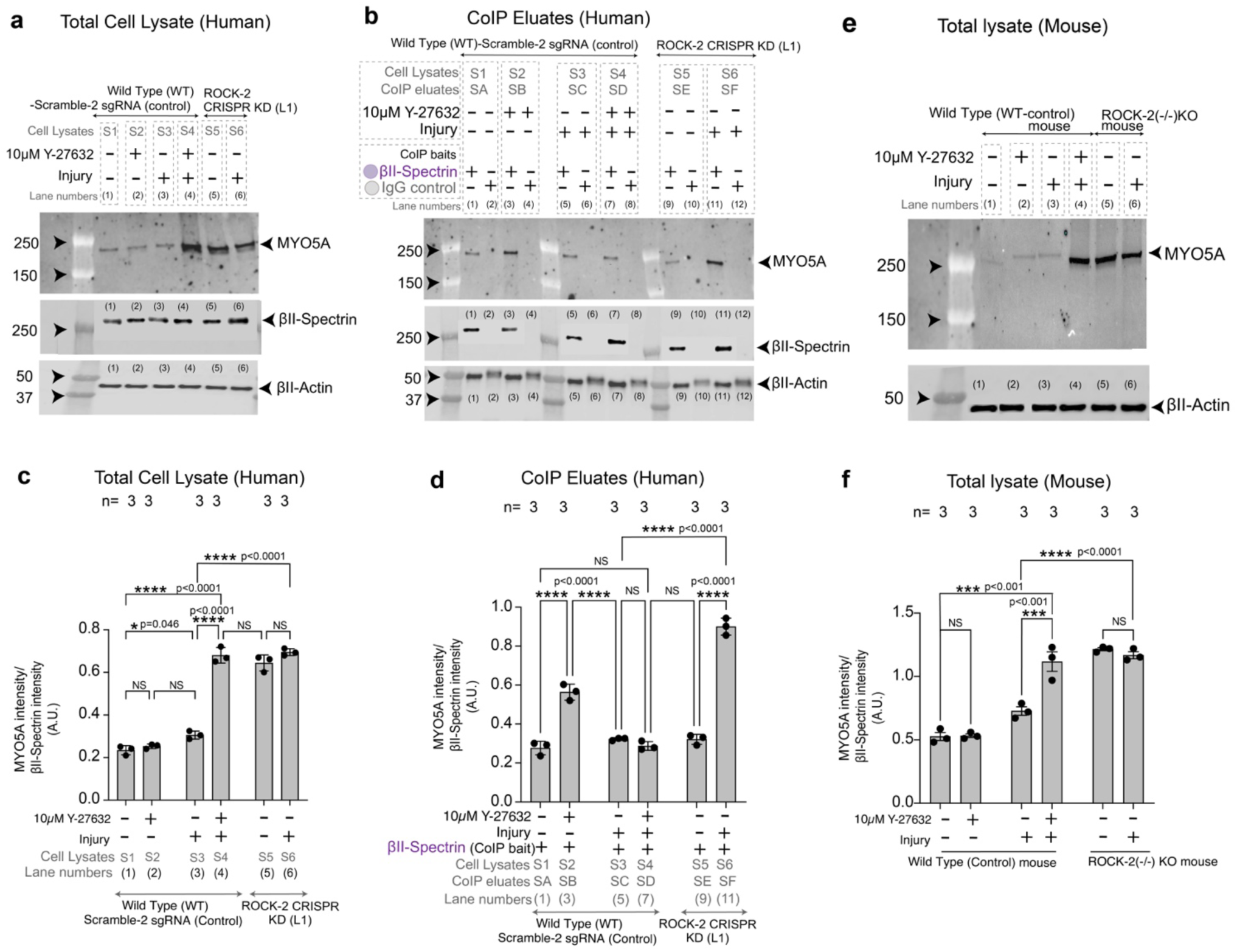
MYO5A upregulation during axon regeneration is enhanced by pharmacological and genetic inhibition of ROCK-2. **a.** The western blot of total cell lysates from wildtype-Scramble-2 sgRNA control and ROCK-2-CRISPR-KD (Line or L1) neurons probed for MYO5A (∼215 kDa), βII-spectrin (∼250 kDa), and βII-actin (loading control, ∼42 kDa) across six experimental conditions varying injury status [−, without /+, with] and treatment with 10 μM Y-27632 [−, vehicle (DMSO) /+, with drug]. **b.** The western blot of co-immunoprecipitation (Co-IP) eluates from wildtype and ROCK-2-KD neurons, probed for MYO5A, βII-spectrin, and βII-actin under the same injury and Y-27632 treatment conditions as in (a). Co-IP was performed using βII-spectrin and IgG (as a control) as baits. **c.** Quantification of MYO5A band intensity normalized to βII-spectrin intensity from total cell lysates in human cortical neurons. In wild-type neurons, Y-27632 treatment increased MYO5A levels in the uninjured condition (*P* = 0.046), and injury with Y-27632 further elevated MYO5A compared to uninjured vehicle (*P* < 0.0001), injured vehicle (*P* < 0.0001), and uninjured Y-27632-treated (P < 0.0001) conditions. A highly significant (*P* < 0.0001) difference was observed between the WT control injured condition without the drug and the ROCK-2-CRISPR KD injured condition. No significant differences were observed between ROCK-2-KD- L1 uninjured and injured conditions. **d.** Quantification of MYO5A band intensity normalized to βII-spectrin intensity from CoIP eluates in human cortical neurons. In wild-type neurons, injury with Y-27632 significantly increased MYO5A–βII-spectrin co-precipitation compared to both injured vehicle (*P* < 0.0001) and uninjured Y-27632-treated (*P* < 0.0001) conditions. In ROCK-2-KD L1 neurons, injury significantly increased MYO5A co-precipitation compared to the uninjured condition (*P* < 0.0001). No significant differences were observed between uninjured wild-type conditions or between injured wild-type vehicle and uninjured conditions. **e.** Western blots of total cell lysates from mouse cortical neurons (wild-type and ROCK-2−/−KO) probed for MYO5A and βII-actin. Conditions include wild-type uninjured without drug (−/−), wild-type uninjured with 10 μM Y-27632 (−/+), wild-type injured without drug (+/−), wild-type injured with 10 μM Y-27632 (+/+), ROCK-2−/− uninjured (−/−), and ROCK-2−/− injured (−/+). **f.** Quantification of MYO5A band intensity normalized to βII-actin intensity from mouse cortical neuron total cell lysates. In wild-type neurons, injury with Y-27632 treatment significantly increased MYO5A expression compared to injured vehicle (P < 0.001). In **c**, **d**, and **f**, data are presented as mean ± SEM with individual data points shown. Statistics: one-way ANOVA with Tukey’s *post hoc* test for multiple comparisons. NS, not significant; \**P* < 0.05; \*\*\**P* < 0.001; \*\*\*\**P* < 0.0001. *P*-values are indicated on the graphs.

To determine whether increased MYO5A levels resulted in a stronger physical interaction with the spectrin complex, we performed western blot analysis on co-immunoprecipitation (CoIP) eluates, using βII-spectrin as bait (Fig. 5b). Quantifying MYO5A band intensity normalized to βII-spectrin in the elutes revealed that Y-27632 treatment significantly enhanced MYO5A’s association with the spectrin complex in WT neurons (*P* < 0.0001). In ROCK-2 CRISPRi KD neurons, MYO5A consistently associated with the spectrin complex, unaffected by injury. The peak MYO5A levels occurred in injured ROCK-2-CRISPR-KD neurons (*P* < 0.0001; Fig. 5d). Notably, the MYO5A enrichment pattern observed in CoIP elutes closely resembled the data from the total cell lysate, confirming that the increased MYO5A protein is not only cytoplasmic but also actively recruited to and physically associated with the βII-spectrin–based MPS complex. To verify these findings in mice, we performed immunoblot analysis on total cell lysates from primary mouse cortical neurons (Fig. 5e). Consistent with human data, MYO5A expression was low in uninjured WT controls and increased significantly after Y-27632 treatment in WT neurons (*P* < 0.001; Fig. 5f). In ROCK-2−/− (KO) mouse cortical neurons, MYO5A levels increased in both uninjured and injured states (*P* < 0.0001; Fig. 5f). The MYO5A increase caused by Y-27632 in both human and mouse neurons demonstrate a conserved axon regulating pathway. These findings suggest a model wherein ROCK-2, positioned at the MPS, locally inhibits MYO5A recruitment to prevent actin-dependent reassembly of the periodic lattice. When ROCK-2 activity is suppressed, MYO5A is de-repressed and recruited to the spectrin complex to support restoration of the MPS during axon regeneration.

### MYO5A drives axon regeneration downstream of ROCK-2 inhibition and maintains βII-spectrin periodicity in intact axons

The biochemical observations suggest MYO5A is a downstream effector of ROCK-2. We sought to determine whether MYO5A is essential for axon regrowth, and whether the improved regenerative ability conferred by ROCK-2 inhibition relies on MYO5A. We developed an AAV-based MYO5A knockdown strategy in the CRISPRi-competent neurons and evaluated its impact on axon regrowth and MPS arrangement in intact axons of both wild-type and ROCK-2-deficient human iPSC-derived cortical neurons. Human cortical neurons were transduced with AAV vectors co-expressing MYO5A-targeting sgRNAs and BFP (MYO5A-sgRNA1-BFP or MYO5A-sgRNA2-BFP) (Supplementary Fig. 10a-b). Phase-contrast and fluorescence imaging verified successful and extensive viral transduction, showing strong BFP expression throughout neuronal cultures for both sgRNA constructs (Supplementary Fig. 10e). Confocal imaging was performed on wild-type (WT) human cortical neurons to verify the efficiency and specificity of the knockdown, including samples without AAV transduction and those transduced with Myo5a-sgRNA1 AAV or Myo5a-sgRNA2 AAV. The neurons were immunostained for βII-spectrin and MYO5A (Supplementary Fig. 10f). SgRNA-mediated knockdown significantly reduced the MYO5A immunofluorescence signal compared to non-transduced controls. Neuronal morphology and βII-spectrin expression remained intact, indicating that MYO5A reduction was effective and specific, without alterations in spectrin levels or neuronal health. Quantitative analysis of MYO5A fluorescence, normalized to βII-spectrin, revealed a significant decrease in MYO5A protein levels in both R1 and R2 knockdown groups compared to the wild type without AAV addition (*P* < 0.0001 for both). There was no significant difference between the two sgRNA constructs (Supplementary Fig. 10g).

To evaluate the role of MYO5A in axon regeneration, we conducted a combined experiment with wild-type and ROCK-2 CRISPR knockdown (KD) human iPSC lines (L1 and L2). These cells were plated in 96-well plates and transduced with MYO5A-sgRNA-AAVs at DIV14, allowing sufficient time for MYO5A knockdown before injury (Fig. 6a). At DIV40, mechanical axotomy was performed, and cultures were fixed three days later (DIV43) for immunostaining and confocal imaging at 100X magnification. Representative confocal images of βIII-tubulin immunostaining at the injury site showed notable differences in axon regrowth among the conditions (Fig. 6b). In wild-type neurons not transduced with AAV, only limited axon regrowth was observed in the scraped area, consistent with our earlier findings (Fig. 6b, panel i). In ROCK-2(KD) neurons from L1 and L2 without AAV, regrowth was notably enhanced (Fig. 6b, panels ii and iii), confirming the pro-regenerative effect of ROCK-2 knockdown previously reported (Fig. 3; Supplementary Fig. 5). Importantly, transduction with either MYO5A-sgRNA1 or MYO5A-sgRNA2 AAV significantly decreased axon regrowth across all three genetic backgrounds—wild-type, ROCK-2(KD) L1, and ROCK-2(KD) L2 (Fig. 6b). Binary images with thresholding revealed a significant decrease in βIII-tubulin-positive neurites entering the injury zone following MYO5A knockdown, even in ROCK-2-deficient lines that typically show robust regenerative capacity. To determine the optimal timing and strength of MYO5A knockdown effects, we examined axon regrowth in wild-type neurons transduced with MYO5A-sgRNA AAV at four different time points: DIV7, DIV14, DIV21, and DIV33 (Fig. 6c). Transduction at DIV7 and DIV14 resulted in consistent, significant reductions in axon regrowth for both sgRNA constructs (*P* < 0.01 and *P* < 0.001). Conversely, transduction at later time points (DIV21 and DIV33) produced more variable outcomes, likely due to insufficient time for MYO5A protein suppression before the DIV40 axon injury. These results indicate that early MYO5A knockdown substantially impairs the regenerative capacity of human cortical neurons and highlight MYO5A’s crucial role during axon regrowth, beyond its function in initial neuronal differentiation and maturation.

**Fig. 6:**
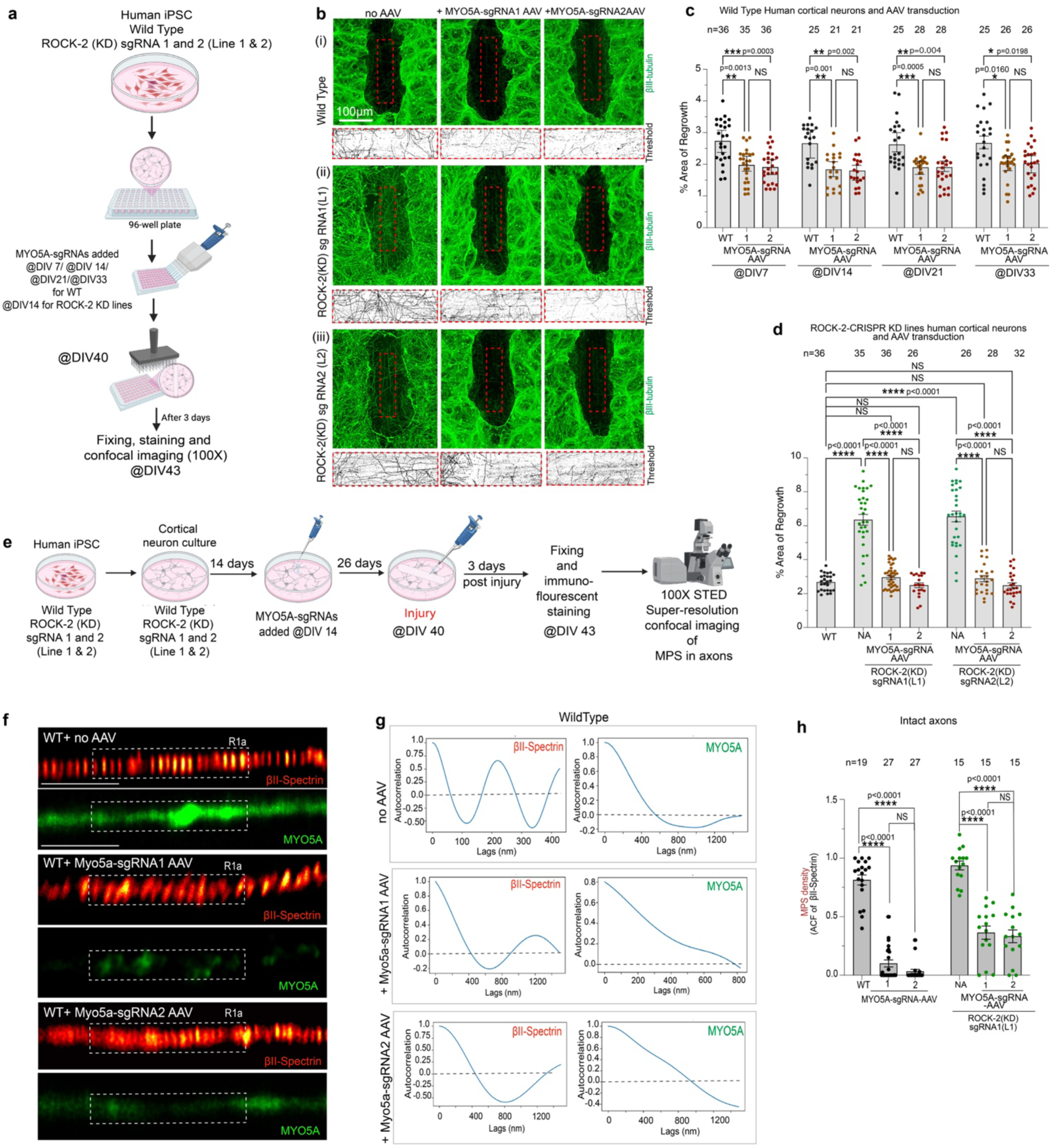
MYO5A knockdown impairs axon regrowth and βII Spectrin periodicity in intact axons for both wild-type and ROCK-2-deficient iPSC-derived neurons. a. A schematic of the experimental workflow. Human iPSCs (wild-type and ROCK-2 CRISPR KD sgRNA1 and sgRNA2 lines) were differentiated into cortical neurons and plated in 96-well plates. MYO5A-sgRNA1 and sgRNA2 AAVs were added at DIV 7, DIV 14, DIV 21, or DIV 33 for wild-type neurons, and only at DIV 14 for ROCK-2 KD lines. At DIV 40, neurons were subjected to a scrape injury and fixed 3 days later (DIV 43) for immunostaining and confocal imaging (10X). b. Representative confocal images of βIII-tubulin immunostaining (green) and corresponding thresholded binary images in regenerating axons at 3 days post-injury. (i) Wild-type neurons treated with no AAV, MYO5A-sgRNA1 AAV, or MYO5A-sgRNA2 AAV. (ii) ROCK-2 KD sgRNA1 (L1) neurons under the same three AAV conditions. (iii) ROCK-2 KD sgRNA2 (L2) neurons under the same three AAV conditions. Red dashed boxes indicate regions of interest used to quantify the percentage of regrowth area. Scale bar: 100 μm. c. Quantification of the percent area of regrowth in the injury or scraped zone in wild-type human cortical neurons treated with MYO5A-sgRNA AAVs (sgRNA1 and sgRNA2) at different time points. MYO5A-sgRNA1 AAV (Brown dots) significantly reduced regrowth compared to wild-type non-transduced controls at all time points tested: DIV 7 (P = 0.0013), DIV 14 (*P* = 0.001), DIV 21 (*P* = 0.0005), and DIV 33 (*P* = 0.0160). MYO5A-sgRNA2 AAV (Red dots) showed a similar trend to MYO5A-sgRNA1 AAV. No significant difference was observed between sgRNA1 and sgRNA2 at any time point. Sample sizes (number of individual wells of the 96-well plate analyzed): DIV 7 (*n* = 36, 35, 36), DIV 14 (*n* = 25, 21, 21), DIV 21 (*n* = 25, 28, 28), DIV 33 (*n* = 25, 26, 26). d. Quantification of the percent area of regrowth in ROCK-2-CRISPRi-KD human cortical neurons. In ROCK-2 KD sgRNA1 (L1) neurons, both no AAV (NA) and wild-type controls showed significantly greater regrowth compared to MYO5A-sgRNA1 and sgRNA2 AAV-treated conditions (P < 0.0001). Similarly, in ROCK-2 KD sgRNA2 (L2) neurons, MYO5A-sgRNA AAV treatment significantly reduced regrowth (P < 0.0001). No significant difference was observed between MYO5A-sgRNA1 and sgRNA2 treatments within each ROCK-2 KD line. Wild-type and ROCK-2 KD NA conditions showed comparable regrowth levels (NS). Sample sizes or number of individual wells of the 96-well plate analyzed: WT (n = 36), ROCK-2 KD L1 NA (n = 35), ROCK-2 KD L1 + sgRNA1 (n = 36), ROCK-2 KD L1 + sgRNA2 (n = 26), ROCK-2 KD L2 NA (n = 26), ROCK-2 KD L2 + sgRNA1 (n = 28), ROCK-2 KD L2 + sgRNA2 (n = 32). In **c** and **d**, data are presented as mean ± SEM with individual data points shown. All experiments were conducted in *N* = 3 independent replicates. Statistics: one-way ANOVA with Tukey’s *post hoc* test for multiple comparisons. NS, not significant; \**P* < 0.05; \*\**P* < 0.01; \*\*\**P* < 0.001; \*\*\*\**P* < 0.0001. P-values are indicated on the graphs. e. Schematic of the experimental workflow. Human iPSCs (wild-type and ROCK-2 KD sgRNA1 and sgRNA2 lines) were differentiated into cortical neurons for 14 days in 35 mm dishes. MYO5A-sgRNA1 and sgRNA2 AAVs were added for transduction at DIV 14. At DIV 40, neurons were subjected to injury, fixed at 3 days post-injury (DIV 43), immunostained, and imaged by 100X STED confocal microscopy to assess MPS organization in axons. f. Representative STED images of βII-spectrin (red hot) and MYO5A (green) in the uninjured axons of wild-type human cortical neurons either treated with MYO5A-sgRNA1 AAV (middle), MYO5A-sgRNA2 AAV (bottom), or no AAV (top), un-transduced control. Dashed boxes indicate regions of interest (R1b or 100 μm proximal to cell body). In wild-type neurons without AAV treatment, βII-spectrin exhibits a clear periodic distribution characteristic of the MPS, which is disrupted upon MYO5A knockdown with both sgRNAs. The MYO5A signal is correspondingly reduced under sgRNA treatment. Scale bars, 5 μm. g. Representative autocorrelation plots of βII-spectrin (left) and MYO5A (right) distributions along the axons for each condition. In the no AAV condition (top), βII-spectrin shows a characteristic periodic autocorrelation pattern with peaks at around ∼190 nm intervals, and MYO5A shows no periodic distribution. Upon treatment with MYO5A-sgRNA1 AAV (middle) or MYO5A-sgRNA2 AAV (bottom), the periodic autocorrelation pattern of βII-spectrin gets abolished, and MYO5A periodicity is lost. Dashed lines indicate the zero-correlation baseline. h. Quantification of MPS density (autocorrelation amplitude of βII-spectrin) in uninjured axons. In wild-type neurons, both MYO5A-sgRNA1 (1) and MYO5A-sgRNA2 (2) AAV treatment significantly reduced MPS density compared to untreated wild-type (WT) (P < 0.0001 for both). No significant difference was observed between sgRNA1 and sgRNA2 groups. In ROCK-2 KD sgRNA1 (L1) neurons, untreated (NA) neurons exhibited high MPS density, which was significantly reduced by both MYO5A-sgRNA1 (*P* < 0.0001) and MYO5A-sgRNA2 (*P* < 0.0001) AAV treatment. No significant difference was observed between MYO5A-sgRNA1 and MYO5A-sgRNA2 treatments within the ROCK-2 KD background. Data are presented as mean ± SEM with individual data points shown as the axonal segments. Sample number (number of intact axons) n= from left to right, 19, 27, 27, 15, 15, and 15 axonal segments. All experiments were conducted in *N* = 3 independent replicates. Statistics: one-way ANOVA with Tukey’s *post hoc* test for multiple comparisons. NS, not significant; \*\*\*\**P* < 0.0001. *P*-values are indicated on the graph.

The key element of our experimental design was epistatic analysis, which tested whether MYO5A knockdown could suppress the increased regrowth phenotype seen in ROCK-2-deficient neurons. Quantifying the axon regrowth area in WT, ROCK-2(KD) L1, and ROCK-2(KD) L2 neurons confirmed that, without AAV transduction (NA), both ROCK-2 KD lines showed significantly greater regrowth than WT, consistent with previous results (Fig. 6d). Notably, transduction with either MYO5A-sgRNA1 or MYO5A-sgRNA2 AAV abolished this increase, bringing axon regrowth levels in both ROCK-2(KD) L1 and L2 lines down to those of non-transduced wild-type controls or lower (*P* < 0.0001; Fig. 6d). In wild-type neurons, reducing MYO5A levels also lowered regrowth below normal levels. The inhibitory impact of MYO5A knockdown was consistently seen across various ROCK-2 KD lines and different MYO5A-targeting sgRNAs, offering strong genetic evidence of their epistatic relationship. This finding has significant mechanistic implications. If inhibiting ROCK-2 promotes regrowth through a mechanism independent of MYO5A, then reducing MYO5A levels shouldn’t reverse the effects of ROCK-2 deficiency. However, because the loss of MYO5A fully suppresses the increased regeneration seen with ROCK-2 deficiency, we conclude that MYO5A acts downstream of ROCK-2 in the signaling pathway promoting axon regrowth. These loss-of-function experiments collectively validate MYO5A as a crucial effector for axon regrowth in human iPSC-derived cortical neurons. These data support a model in which ROCK-2 functions as a molecular brake on MYO5A-driven cytoskeletal remodeling at the MPS which in turn promotes greater axon regeneration.

A crucial unresolved question is whether MYO5A directly contributes to organizing or maintaining the MPS itself. If MYO5A contributes to the βII-spectrin periodic lattice’s integrity, its absence may disrupt MPS organization in healthy, uninjured axons where the MPS is usually well established. To test this prediction, we assessed how MYO5A knockdown influences βII-spectrin periodicity in intact axons, using STED super-resolution microscopy. Human iPSC-derived cortical neurons from wild-type and two separate ROCK-2 CRISPR knockdown lines (sgRNA1, L1; sgRNA2, L2) were cultured and then transduced with AAV vectors carrying MYO5A-targeting sgRNAs (either MYO5A-sgRNA1 or MYO5A-sgRNA2) at DIV14 (Fig. 6e). This timing allowed sufficient depletion of MYO5A protein to evaluate MPS integrity in mature axons. After mechanical injury at DIV40, the neurons were fixed at DIV43 (Fig. 6e). Cultures were stained for βII-spectrin and MYO5A, and intact axonal segments close to the cell body (R1a) that were uninjured were imaged using STED super-resolution confocal microscopy at 100X magnification (Fig. 6e). This experimental design enabled us to examine MPS organization in axonal segments that were stable and not regenerating. It isolates the effect of MYO5A loss on MPS stability from injury-related reformation. Dual-color STED imaging of intact wild-type axons without AAV showed the expected co-distribution of βII-spectrin and MYO5A. The βII-spectrin protein displayed regularly spaced puncta along the axon shaft, indicating a periodic organization roughly every 190 nm (Fig. 6f, WT + no AAV). In these MPS-intact axons, MYO5A itself exhibited a more diffuse, less periodic pattern that only partially overlapped with the spectrin lattice, consistent with limited MYO5A interaction with spectrin under basal conditions (Figs. 4, 5). The axons transduced with MYO5A-sgRNA1 AAV or MYO5A-sgRNA2 AAV showed a significant decrease in MYO5A immunofluorescence, confirming effective knockdown coupled with a clear disruption of the periodic βII-spectrin pattern (Fig. 6g). The typical periodic pattern of the MPS was replaced by a more diffuse, continuous, and irregular distribution of βII-spectrin, indicating that the periodic lattice lost stability after MYO5A suppression. Autocorrelation analysis of βII-spectrin and MYO5A fluorescence intensity profiles along intact axons quantitatively verified these qualitative observations (Fig. 6h). Transduction with either MYO5A-sgRNA1 or MYO5A-sgRNA2 AAV significantly reduced ACF amplitude (*P* < 0.0001 for both), with similar effects from both sgRNAs. The disruption of this stable structure upon MYO5A depletion implies that even in mature axons, the MPS requires active maintenance by MYO5A. This dependence is striking in light of the documented slow turnover of MPS spectrin^14^ and MPS resistance to pharmacological perturbation of actin dynamics ^12^. As an actin-based motor protein^31^, MYO5A may contribute to MPS maintenance through several non-mutually exclusive mechanisms such as transporting spectrin tetramers or other MPS components to sites of lattice turnover, generating mechanical forces that stabilize the actin ring-spectrin tetramer interface, or facilitating the incorporation of newly synthesized spectrin into the existing periodic framework.

To determine whether the disruption of the MPS caused by MYO5A loss is epistatic to ROCK-2 status, we conducted the same analysis on ROCK-2-CRISPR-(KD) sgRNA1 (L1) neurons (Fig. 6h). Non-transduced ROCK-2 KD neurons showed βII-spectrin ACF values comparable to wild-type controls, confirming that ROCK-2 knockdown alone does not affect MPS periodicity in intact axons. This aligns with our earlier observation that βII-spectrin periodicity remains intact in ROCK-2 KD neurons (Fig. 3i-l). However, transducing ROCK-2 KD neurons with either MYO5A-sgRNA1 or MYO5A-sgRNA2 AAV caused a significant reduction in βII-spectrin ACF amplitude (*P* < 0.0001; Fig. 6f), indicating disruption similar to that seen in wild-type neurons. This result indicates that MYO5A is crucial for maintaining MPS, regardless of ROCK-2 status, supporting the hypothesis that MYO5A acts downstream of ROCK-2 to provide essential support for the periodic skeleton. Our STED imaging (Fig. 6g) shows partial colocalization of MYO5A with βII-spectrin. ROCK-2, as a partly periodic component of the axonal MPS (Fig. 3), limits MYO5A expression and binding to the spectrin complex (Figs. 4, 5). Conversely, MYO5A is essential for maintaining MPS periodicity in intact axons (Fig. 6f-h) and for axon regrowth after injury (Fig. 6b-c). This ROCK-2/MYO5A axis is a newly identified signaling module that links nanoscale cytoskeletal organization with overall regenerative capacity.

### MYO5A overexpression is sufficient to enhance axon regrowth and accelerate MPS recovery

Our loss-of-function experiments show that MYO5A is essential for rapid axon regeneration, and its removal abolishes the pro-regenerative effect observed in ROCK-2-deficient neurons (Fig. 6). Furthermore, reducing MYO5A levels disrupted the periodic pattern of βII-spectrin even in stable uninjured axon segments (Fig. 6), suggesting MYO5A is vital for maintaining the MPS structure. Based on these findings, we considered whether boosting MYO5A activity might encourage axon regeneration, effectively bypassing the need to inhibit ROCK-2 by directly activating its downstream target. To evaluate this prediction, we overexpressed MYO5A in wild-type human iPSC-derived cortical neurons and observed its effect on axon regeneration following mechanical injury. Wild-type human iPSC-derived cortical neurons were cultured in 96-well plates and transduced at DIV14 with either a control lentivirus expressing mCherry (pLV(Exp)-EGFP/Puro-EF1A-mCherry) (Fig. 7a; Supplementary Fig. 10d), or a MYO5A-overexpressing lentivirus encoding MYO5A isoform 1 fused to an HA tag (pLV[Exp]-SYN1-(MYO5A-Iso-1)HA tag) (Fig. 7a; Supplementary Fig. 10c). A third group did not receive any lentivirus, serving as a baseline control (Fig. 7a). At DIV40, a mechanical scrape injury was performed, and cultures were fixed 3 days later. They were then immunostained for βIII-tubulin, anti-HA tag (to detect MYO5A-overexpression), mCherry (to identify the control viral transduction), and DAPI, followed by confocal imaging (Fig. 7a). The HA tag enabled clear detection of the exogenous protein. Confocal images of the scrape injury site demonstrated notable differences in axon regrowth across the three conditions (Fig. 7b). In the no-lentivirus group, some neurite regrowth was observed in the scraped area, with sparse βIII-tubulin-positive processes extending into the injury zone and no anti-HA tag signal, as expected. In the control lentivirus group, mCherry expression confirmed successful viral transduction, but the extent of axon regrowth into the scraped area was indistinguishable from the no-virus group, indicating that lentiviral transduction alone did not affect regenerative capacity. Neurons transduced with MYO5A-overexpressing lentivirus exhibited anti-HA tag immunoreactivity, confirming effective expression of the exogenous MYO5A and significantly enhanced neurite regeneration into the injury zone (Fig. 7b). Thresholded binary images of βIII-tubulin in the injury zones revealed a significantly greater density of regrowing axons in MYO5A-OE cultures compared to both control groups (Fig. 7b). Quantifying axon regrowth as the percentage of βIII-tubulin-positive area within the injury zone statistically confirmed these findings (Fig. 7c). Overexpression of MYO5A markedly improved regrowth, surpassing the no-virus group (*P* < 0.0001) and the control lentivirus group (*P* < 0.0001), which did not show a significant difference from one another (Fig. 7c). MYO5A-OE neurons achieved regrowth levels approximately three-fold higher than controls, comparable to the levels seen with pharmacological ROCK-2 inhibition (Supplementary Fig. 4e-f) and genetic ROCK-2 knockdown (Supplementary Fig. 5b-c). This gain-of-function finding supports the loss-of-function data from Fig. 6, providing strong evidence that MYO5A is both necessary and sufficient for promoting axon regrowth in human cortical neurons.

**Fig. 7:**
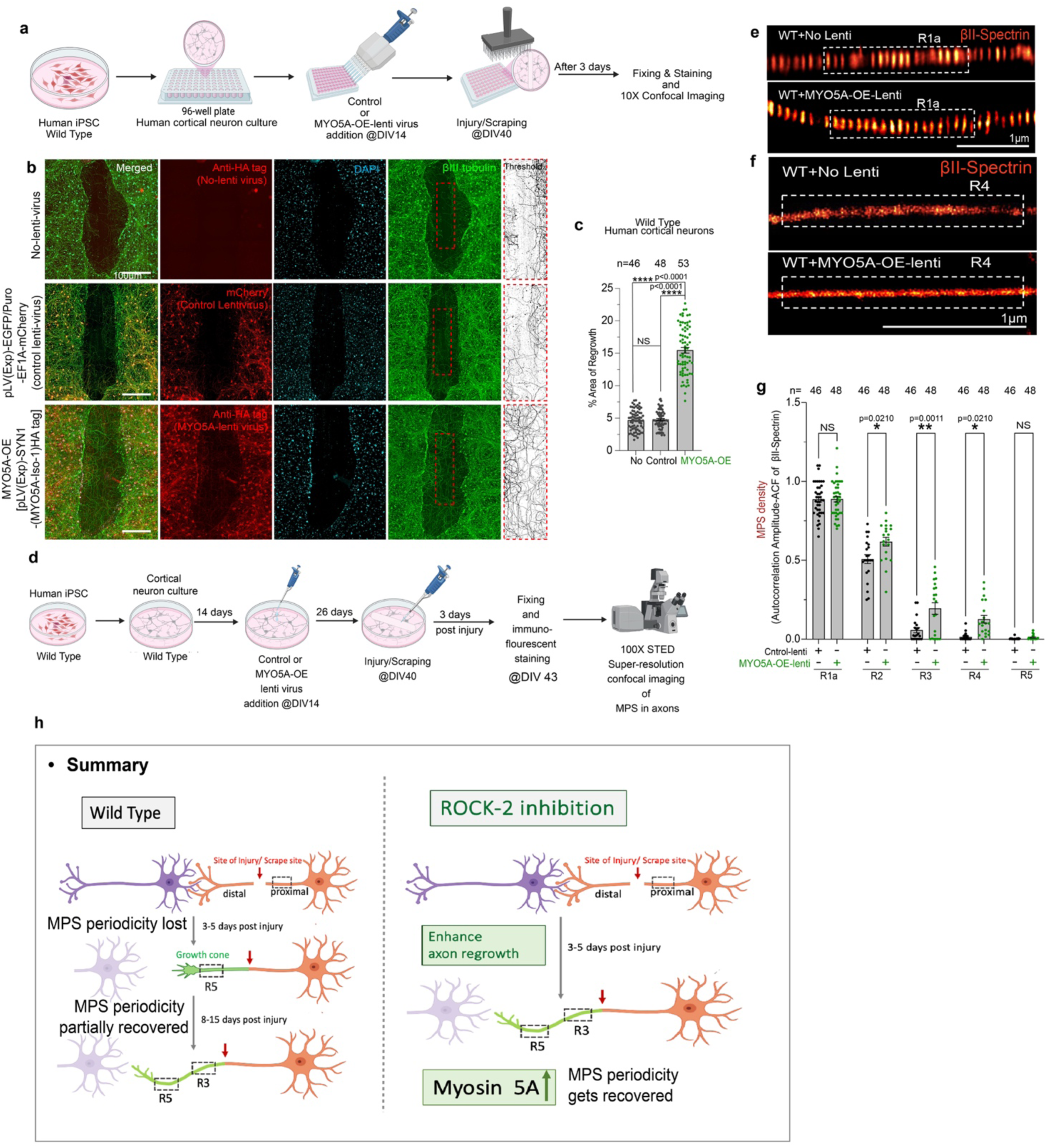
Overexpression of MYO5A shows a significant enhancement in axon regrowth. a. Schematic of the experimental workflow. Human iPSC-derived wild-type cortical neurons were cultured in 96-well plates. At DIV 14, neurons were transduced with either a control lentivirus (pLV(Exp)-EGFP/Puro-EF1A-mCherry) or a MYO5A-overexpression (OE) lentivirus [pLV(Exp)-SYN1-MYO5A-Iso-1-3X-HA tag] or left untreated (no lentivirus). At DIV 40, neurons were subjected to a scrape injury and fixed 3 days post-injury for immunostaining and 10X confocal imaging. b. Representative confocal images of regenerating axons at 3 days post-injury across three conditions. Top row: no lentivirus control, showing merged image, anti-HA tag (no signal, confirming absence of MYO5A-OE construct), DAPI, βIII-tubulin, and corresponding βIII-tubulin thresholded binary image of the regrowing axons in the indicated ROIs. Red dashed boxes indicate regions of interest. Middle row: control lentivirus-transduced neurons, showing merged image, mCherry (confirming successful control virus transduction), DAPI, βIII-tubulin, and thresholded image. Bottom row: MYO5A-OE lentivirus-transduced neurons, showing merged image, anti-HA tag (confirming MYO5A overexpression, red), DAPI, βIII-tubulin, and thresholded image. MYO5A-OE neurons display visibly enhanced axonal regrowth compared to both control conditions. Scale bar: 100 μm. c. Quantification of the percent area of regrowth in wild-type human cortical neurons. MYO5A-OE significantly increased the area of regrowth compared to both the no lentivirus condition (*P* < 0.0001) and the control lentivirus condition (*P* < 0.0001). No significant difference was observed between no lentivirus and control lentivirus conditions. Sample sizes (number of analyzed wells in 96-well plate): no lentivirus (n = 46), control lentivirus (n = 48), MYO5A-OE (n = 53). Data are presented as mean ± SEM with individual data points shown. All experiments were conducted in N = 3 independent replicates. Statistics: one-way ANOVA with Tukey’s POST HOC test for multiple comparisons. NS, not significant; \*\*\*\**P* < 0.0001. *P*-values are indicated on the graph. d. Schematic of the experimental workflow for STED super-resolution imaging. Wild-type human iPSC-derived cortical neurons were transduced at DIV14 with control or MYO5A-OE lentivirus, subjected to scrape injury at DIV40, fixed at DIV43, and imaged at 100X using STED super-resolution confocal microscopy for βII-spectrin periodicity in axons. **e-f.** STED super-resolution images of βII-spectrin (red hot) at regions R1a (**e**) and R4 (**f**) in regrowing axons of WT neurons without lentivirus (top) and with MYO5A-OE lentivirus (bottom) at 3 dpi. At R1a, both conditions display comparable periodic organization. At R4, MYO5A-OE neurons show enhanced βII-spectrin periodicity compared with controls. Dashed boxes indicate segments used for autocorrelation analysis. Scale bars, 1 μm. **g.** Quantification of MPS density (autocorrelation amplitude of βII-spectrin) across five axonal regions (R1a–R5) in WT neurons with control lentivirus (grey) or MYO5A-OE lentivirus (green). MYO5A overexpression significantly enhanced periodicity at R2 (*P* = 0.0210), R3 (*P* = 0.0011), and R4 (*P* = 0.0210), with no significant changes at R1a or R5. Sample sizes (number of axonal segments) from left to right, 46, 48, 46, 48, 46, 48, 46, 48, 46, and 48. Data are mean ± SEM with individual data points; *N* = 3 independent replicates. One-way ANOVA with Tukey’s *post hoc* test. NS, not significant; \**P* < 0.05; \*\**P* < 0.01. **h.** Summary model. In wild-type neurons (left), MPS periodicity is lost at 3–5 dpi and partially recovers by 8–15 days at intermediate regions (R3), while distal segments remain aperiodic. Upon ROCK-2 inhibition (right), axon regrowth is enhanced, MYO5A expression is upregulated, and MPS periodicity recovery is accelerated to 3 dpi. MYO5A acts downstream of ROCK-2 as a necessary and sufficient effector linking nanoscale cytoskeletal reassembly to axon regeneration.

To study the subcellular localization of MYO5A in regenerating axons, we used high-magnification confocal imaging of growth cones stained for MYO5A and F-actin (Supplementary Fig. 10h). In addition to its presence along the axon shaft, MYO5A was concentrated at the growth cone and co-localized with F-actin in both filopodia and lamellipodia, indicating its involvement in actin-dependent processes at the leading edge of regenerating axons (Supplementary Fig. 10h). This pattern supports MYO5A’s established role as an actin-based motor protein and suggests it modifies growth cone dynamics during axon regeneration ^45^, in addition to its function in MPS organization.

The neuronal MPS is more than just a static scaffold; it acts as an actomyosin network where non-muscle myosin II (NMII) bipolar filaments attach to the periodic actin rings and generate radial contractile forces that regulate axonal diameter^13^. Supporting this, acute inhibition of NMII with blebbistatin, depletion of NMIIA and NMIIB, or knocking down βII-spectrin to dismantle the MPS all resulted in a 20–30% increase in average axon diameter. Similar phenotypes in βII-spectrin and NMII disruptions suggest that the MPS is the structural basis by which actomyosin contractility controls axon size. Loss of α-adducin, a component that caps actin, also enlarges axon diameter, further confirming that the periodic skeleton’s integrity, rather than any single motor protein, governs this structural constraint^13^. Since MYO5A is crucial for preserving βII-spectrin’s periodic structure in intact axons, we hypothesized that axon diameter might be a marker for MPS integrity after MYO5A disruption. If depleting MYO5A destabilizes the periodic skeleton, it resembles MPS disruption and reduces radial restriction on axonal caliber. We therefore investigated whether altering MYO5A altered axon diameter. Measuring axon diameter approximately 50 µm from the cell body in healthy, uninjured axons. We found that MYO5A knockdown (MYO5A-sgRNA1-AAV) significantly increased axon diameter relative to controls (*P* < 0.0001), whereas MYO5A overexpression produced no significant change (NS; Supplementary Fig. 10i). These findings indicate that MYO5A helps regulate axon size, presumably by maintaining the actomyosin network linked to the MPS. The absence of MYO5A might compromise the structural constraints that the periodic skeleton exerts on axonal shape.

Having established that MYO5A overexpression promotes macroscale axon regrowth, we then examined whether it also accelerates the nanoscale reassembly of the MPS in regenerating axons—a feature previously associated with ROCK-2 inhibition (Fig. 3). To investigate this, we conducted a parallel experiment using 35-mm culture dishes for STED super-resolution imaging (Fig. 7d). Wild-type human iPSC-derived cortical neurons were cultured, transduced at DIV14 with either control or MYO5A-OE lentivirus, injured with a scrape at DIV40, and fixed at DIV43 for immunostaining. We then performed 100X STED super-resolution confocal imaging to observe βII-spectrin periodicity along the regenerating axons (Fig. 7d). Proximal intact region R1a STED images demonstrated that βII-spectrin preserved its typical periodic pattern in both control (WT + No Lenti) and MYO5A-OE conditions (Fig. 7e). This indicates that the MPS structure was similar across groups in uninjured axonal segments. In contrast, STED images of the distal regenerating region R4 showed significant differences (Fig. 7f). In control neurons without lentivirus, βII-spectrin at R4 appeared mostly aperiodic, confirming earlier observations of lost MPS periodicity in distal regenerating segments (Fig. 1). Conversely, in MYO5A-OE neurons, βII-spectrin at R4 exhibited increased periodicity, with more defined periodic puncta compared to the diffuse signal in controls (Fig. 7f). Quantifying MPS density, expressed as the autocorrelation amplitude (ACF) of βII-spectrin across five axonal regions (R1a, R2, R3, R4, R5) confirmed these results statistically (Fig. 7g). At R1a, MPS density was high and similar between control and MYO5A-OE conditions (NS), indicating intact MPS in uninjured proximal segments. MYO5A overexpression significantly increased MPS density at R2 (*P* = 0.0210), R3 (*P* = 0.0011), and R4 (*P* = 0.0210) relative to control neurons, indicating faster βII-spectrin periodicity recovery during regeneration in these regions. No significant difference was observed at R5 (NS), near the growth cone, where MPS periodicity remained low in both groups (Fig. 7g). These findings show that MYO5A overexpression can accelerate MPS reassembly in regenerating axons at 3 days post-injury, recapitulating the effects of pharmacological and genetic ROCK-2 inhibition (Fig. 3).

## DISCUSSION

We studied how membrane-associated periodic skeleton (MPS) dynamics change during axon regeneration and identified a ROCK-2/MYO5A pathway that connects the periodic spectrin lattice to the regenerative process. We used to STED microscopy after *in vitro* scrape injury of primary mouse cortical neurons as well as human iPSC-derived cortical neurons and motor neurons. We observed that the ∼190 nm βII-spectrin periodicity of the mature MPS is initially absent from regenerating axons but slowly and partially recovers over 8–15 days. Reformation of the MPS is greatly accelerated by ROCK-2 inhibition either pharmacologically or genetically. Proteomic analysis identified MYO5A as an interactor of the βII-spectrin complex regulated by injury and ROCK-2. Loss- and gain-of-function studies demonstrated that MYO5A is both necessary and sufficient for axon regrowth and MPS reformation after injury, as well as being essential for maintaining MPS periodicity in uninjured axons.

Based on these data, we propose a mechanistic model describing MPS dynamics during axon regeneration (Fig. 7h). In wild-type neurons, mechanical injury damages the membrane-associated periodic skeleton at the injury site. In the first 3–5 days post-injury, MPS periodicity is absent from regenerating axon segments, limiting regrowth, especially at the distal growth cone where disruption is essentially complete. Between 8–15 days, some MPS recovery occurs in intermediate axonal regions (R3), but the most distal areas near the growth cone remain aperiodic. ROCK-2 inhibition, through drugs or genetics, markedly enhances axon regrowth. It also raises MYO5A expression and its recruitment to the βII-spectrin complex, promoting early restoration of MPS periodicity in regenerating axons (Fig. 7g). Recognizing MYO5A as the key downstream effector explains how ROCK-2 inhibition boosts both overall axon regrowth and nanoscale cytoskeletal reassembly. MYO5A, an actin-based motor protein, is recruited to the spectrin complex after ROCK-2 inhibition, likely facilitating transport, integration, and stabilization of MPS components into the growing lattice, consistent with its known roles in actin-dependent cargo transport and membrane organization in neurons.

A previous βII-spectrin nanoscale study imaged distal MPS re-assembly following axotomy in patches that expand and merge^16^. However, the signaling mechanisms driving MPS reconstruction during axon regrowth remained undefined. Our findings uncover a previously unrecognized signaling pathway involving ROCK-2 and MYO5A that organizes the MPS and promotes overall axon regrowth following injury. This pathway links a clinically relevant kinase target^21,26^ to nanoscale cytoskeletal arrangement and successful regeneration. Since ROCK inhibitors are already used or tested in cardiovascular, ocular, and neurological conditions^21^, our findings suggest that their pro-regenerative effects may partly result from relieving MYO5A repression and restoring the periodic membrane skeleton. The conservation of this pathway in both mouse and human neurons, as well as across cortical and motor neuron types, underscores its crucial role and promising therapeutic potential for improving axon repair after traumatic injury and in neurodegenerative diseases.

Existing knowledge of MPS function primarily concerns mechanical stability^5,6^, signaling^7^, axon diameter and conduction^9^, and degeneration^18,19^, in which MPS disassembly precedes axonal fragmentation and acts as a structural checkpoint for axon integrity. Our data present another scenario, in which the MPS is disrupted during regeneration, but in this case, the disruption reflects the immature state of newly extended axon segments rather than a terminal collapse. Regrowing axons reestablish periodic βII-spectrin organization in a proximal-to-distal gradient that mimics developmental assembly^14,16^, and suggests that regeneration may partially recapitulate development. During development, kinesin-dependent transport moves spectrin tetramers along the axon, where they are integrated into actin ring–spectrin lattice units. These units nucleate locally and merge into a continuous periodic structure^14–16^. The incomplete MPS recovery of regenerating axons observed at 8–15 dpi, especially the persistent absence of periodicity near the growth cone, aligns with spectrin delivery and local lattice nucleation being limiting factors for regrowing axons. This fits recent evidence that proper MPS assembly in *C. elegans* requires a tightly balanced supply of spectrin via kinesin-dependent transport^15^, and that actin nucleation is a prerequisite for MPS initiation^12^.

ROCK-2 functions as a negative regulator of MPS reassembly. The Rho/ROCK pathway is a recognized inhibitor of axon regeneration, primarily by promoting growth cone collapse and increasing actin–myosin contractility^21,26–29^. Y-27632 and Fasudil facilitate regeneration in animal models of spinal cord injury^26–29^, and Fasudil has been administered to more than 30,000 patients for cerebrovascular conditions, demonstrating a favorable safety profile^28^. However, the downstream mechanisms are still unclear and are mostly thought to involve growth cone actin dynamics. We observe that ROCK-2 also restricts the reassembly of the periodic spectrin lattice in regenerating axons. Therefore, inhibiting ROCK-2 not only promotes axon extension but also allows them to develop a more mature cytoskeletal structure.

The structural foundation for local control is that ROCK-2 exhibits approximately 190 nm periodicity along the axon and partially overlaps with βII-spectrin (Supplementary Fig. 5d–g). Consequently, ROCK-2 becomes part of the group of proteins that have a periodic arrangement at or near the MPS, which includes spectrin, actin, adducin, ankyrin, sodium channels, and receptor tyrosine kinases^1,2,7^. Its periodic localization enables it to phosphorylate nearby substrates at each repeat, influencing lattice stability or turnover. ROCK-2 knockdown did not affect βII-spectrin periodicity under normal conditions, indicating ROCK-2 regulates MPS dynamics rather than being a structural element. This distinction may be important for therapeutic targeting.

To our knowledge, identifying MYO5A as the key downstream effector represents a previously unrecognized role for this motor. MYO5A, a class V unconventional myosin that transports cargoes such as organelles, mRNA, and signaling molecules along actin filaments^31,32^. In neurons, it has been linked to ER transport into Purkinje cell dendritic spines^33^, synaptic vesicle trafficking, and growth cone motility. Our data indicate that MYO5A physically interacts with the βII-spectrin complex. Its recruitment to the spectrin interactome relies on injury and ROCK-2 activity. Additionally, MYO5A is essential for MPS periodicity in healthy axons and for regrowth following injury. The epistasis result shown in Fig. 6 provide clear evidence that reducing MYO5A levels reverses the increased regrowth observed in ROCK-2-deficient cells. This suggests that ROCK-2 inhibition promotes regeneration via MYO5A.

The details of MYO5A action in maintaining and assembling the MPS requires further, and multiple parallel mechanisms might be involved. As an actin-based motor protein, MYO5A may transport spectrin tetramers or other MPS components to areas where the lattice is assembled or turned over. Although the mature MPS is highly stable, there is ongoing low-level turnover. Our observation that MYO5A knockdown disrupts periodicity even in uninjured axons (Fig. 6) suggests this turnover depends on active MYO5A. Additionally, MYO5A could exert forces that stabilize the actin ring–spectrin tetramer interface or assist in inserting new spectrin into the lattice. Moreover, since MYO5A co-immunoprecipitated with actin-regulatory proteins such as CAPZA1, CAPZB, TMOD2, and DBN1 (Fig. 4c-d), it might coordinate a multi-protein actin-regulatory complex involved in actin ring nucleation and filament capping within the MPS.

The ROCK-2/MYO5A axis appears to be mechanistically specific. PTEN knockdown promotes regrowth to a similar extent via the PI3K/Akt/mTOR pathway^39,40^. However, it does not restore MPS periodicity in axons that are regrowing (Supplementary Fig. 8a-d). Faster regrowth alone is insufficient to restore the MPS, indicating that MPS recovery is not just a passive consequence of axon extension. Instead, this suggests that ROCK-2 acts as a specific regulator of spectrin periodicity. It also indicates that MPS reassembly and growth cone movement are driven by at least partially separate molecular mechanisms during regeneration, and that treatments engaging both, such as ROCK-2 inhibition, may be more effective than those focusing solely on promoting growth.

The ROCK-2/MYO5A/MPS axis is conserved between mouse and human neurons, as well as across cortical and motor neuron subtypes. The human iPSC results are relevant to translation, while the motor neuron data are directly pertinent to specific clinical settings. Motor neurons have exceptionally long axons, making them especially susceptible to cytoskeletal disruption^4^. Furthermore, inhibiting ROCK-2 improves both axon regrowth and MPS recovery (Supplementary Figs. 5e-f and Fig 3f–g). This pathway presents a potential target for degenerative conditions such as amyotrophic lateral sclerosis, as well as for repair after traumatic spinal cord injury^8^.

The translational relevance is evident. ROCK inhibitors are currently approved or in advanced trials for conditions such as cerebral vasospasm (Fasudil, in Japan and China), glaucoma (ripasudil, netarsudil), and various cardiovascular diseases^21^. Our findings suggest that pro-regenerative effects might partly occur by alleviating MYO5A repression and restoring the periodic skeleton. These supports combining ROCK inhibitors with strategies that increase MYO5A expression or activity. Since overexpressing MYO5A alone can stimulate regrowth in wild-type neurons (Fig. 7b-c), it represents a potential target separate from ROCK-2.

Myosin motor proteins are essential in regulating axon regeneration, with MYO5A and NMII serving different functions. NMII was shown to generate radial forces that determine axon diameter, and inhibiting NMII results in wider axons without affecting the periodic lattice^13^. Blebbistatin has been demonstrated to promote neurite outgrowth, supporting regeneration following nerve injury, particularly when applied locally^37^. MYH9 knockdown mimics NMII inhibition effects. In contrast, MYO5A functions differently by being recruited after injury to support the reassembly of the spectrin lattice, which is crucial for MPS maintenance, even in uninjured axons. NMII inhibition rapidly diminishes barriers to growth cone progression, whereas MYO5A contributes to the structural maturation of regrowing axons. Our proteomic data indicate that multiple myosins interact with the MPS, coordinating both mechanical and transport functions. Combining NMII inhibition with approaches to elevate MYO5A could improve regeneration, as they act on distinct yet complementary cytoskeletal mechanisms.

Several questions may emerge from these studies. Does MPS restoration in regenerating axons correlate with functional recovery, such as action potential propagation, axonal transport efficiency, and synaptogenesis? The MPS structures help organize axonal ion channels^1^ and sets axon diameter and conduction velocity^9^. Therefore, faster MPS reassembly may potentially result in the earlier functional maturation of regenerated axons. Does MYO5A work together with other MPS regulators like adducin, non-muscle myosin II, and ankyrin^2,13^? The presence of myosin light chains (MYL12B, MYL6) in our screen suggests a potential interaction between class V and class II myosins at the MPS, warranting further investigation.

This study has limitations. While our scrape injury model is reproducible and manageable, it does not recapitulate the inhibitory environment of the injured CNS, including myelin-associated inhibitors, chondroitin sulfate proteoglycans, and the glial scar^46^. It remains untested whether ROCK-2 inhibition accelerates MPS reassembly in regrowing axons *in vivo*, especially in the presence of these barriers. Our proteomic screen identified 14 candidate interactors influenced by injury and ROCK-2 status. The functions of the other 13, including actin capping proteins, Tropomodulin, and Drebin, in MPS dynamics and regeneration, remain to be tested. Although our results are consistent across mouse and human neurons, gaining a full understanding of how MPS restoration influences axon maturation, electrophysiology, and synapse formation during axon regeneration will necessitate long-term *in vivo* studies.

Our findings indicate that inhibiting ROCK-2 upregulates MYO5A expression, which plays a role in the MPS/regrowth axis by connecting a kinase to the organization and regeneration of the axonal skeleton. MYO5A is both essential and enough on its own to produce this effect, providing a new target for axon repair approaches and revealing an unforeseen function for a class V myosin in maintaining a highly conserved neuronal structure.

## ON-LINE METHODS

## METHODS

### Mice

All animal experiments strictly followed protocols approved by Yale University’s IACUC. Animals were kept in a pathogen-free facility with a 12-hour light/dark cycle and had unlimited access to food and water at 21–23 °C with 50 ± 20% humidity. For primary cortical neuron cultures, neurons were harvested from wild-type (WT) Po pups. Female C57BL/6 mice with P0 pups (LMWP PUPS 0006E) were delivered from Charles River Laboratories. ROCK-2 knockout studies used heterozygous [ROCK-2 (+/−)] breeders to produce homozygous ROCK-2(−/−) pups. Cortical neurons were obtained from the anterior cortex of P0 pups, as described in the primary mouse cortical neuron culture section below. Because homozygous Rock-2 deletion causes placental issues, intrauterine growth retardation (IUGR), and high perinatal mortality, ROCK-2(−/−) mice cannot be maintained as a stable colony and are generated via interbreeding heterozygous ROCK-2(+/−) parents ^47^. Surviving P0 ROCK-2(−/−) pups can be distinguished from WT littermates by two physical features: smaller body size (runt phenotype) due to IUGR, and hematomas in the hind limbs appearing as reddish discoloration. These features are initial indicators; for definitive genotypic identification, each pup was cultured separately in a 35-mm dish, and tail biopsies were collected for PCR genotyping or through Transnetyx (https://www.transnetyx.com/). WT littermates from the same heterozygous cross served as controls.

### Primary mouse cortical neuron culture

Primary cortical neurons were isolated from P0 pups of WT or ROCK-2(−/−) KO mice. Bilateral cortices were dissected in ice-cold, calcium-free Hibernate E medium (BrainBits, cat. no. HACA500), with meninges and blood vessels carefully removed under a dissection microscope. The tissues then underwent enzymatic digestion at 37 °C for 30 minutes in a freshly prepared solution comprising 4 mL Mg2+/Ca2+-free Hanks’ balanced salt solution (HBSS: Gibco^TM^, cat. no. 14170-112), 144 μL papain (20 U/mL; Worthington, cat. no. LK003178, reconstituted in 5 mL HBSS), 25 μL EDTA (0.5 M: Sigma Aldrich, cat. no. 03690), 7.5 μL CaCl2 (1 M: ThermoFisher Scientific, cat. no. J63122.AD), and 500 μL DNase I (10 mg/mL; Sigma, cat. no. DN25), filtered through a 0.2 μm syringe filter. After digestion, the tissue was rinsed twice with Neurobasal medium and then triturated 10–12 times using a P1000 pipette. The cell suspension was subsequently filtered through a 40 μm cell strainer (Corning, cat. no. 352340), and cell viability was assessed using Trypan Blue solution, yielding typical yields of 60,000–80,000 cells per pup. Cells were seeded at 10 × 10^3^ cells per 100 μL onto poly-D-lysine (PDL)-coated 96-well plates (Corning, cat. no. 356640), or at 100,000–200,000 cells per 1,000 μL onto PDL/laminin-coated 35-mm glass-bottom dishes (iBidi, cat. no. 81148) for STED imaging. Culture medium consisted of Neurobasal-A (Gibco) supplemented with 1 mM sodium pyruvate (Gibco, cat. no. 11360-070), 2 mM GlutaMAX (Gibco, cat. no. 35050061), and 1× B-27 (Gibco, cat. no. 17504044). Cultures were maintained at 37 °C with 5% CO2 and left undisturbed for 7 days after plating. Half-medium changes were performed at DIV7 and then every 7 days thereafter. Mechanical scrape injury was performed at DIV10–11 using a floating pin tool with FP1-WP pins (V&P Scientific) for 96-well plates or a sterile 200 μL pipette tip for 35-mm dishes. Cultures were fixed at 3 or 8 days post-injury (dpi).

### Human iPSC lines and maintenance

Human-induced pluripotent stem cells (iPSCs) containing a doxycycline-inducible neurogenin-2 (NGN2) transgene inserted at the AAVS1 safe harbor locus (WT line, also stably expressing CRISPRi machinery; parental line known as CRISPRi-i3N iPSCs) were cultured in Essential 8 Medium (Gibco, cat. no. A1517001) on Vitronectin-coated plates (Gibco, cat. no. A31804; diluted 1:100 in sterile DPBS) at 37 °C with 5% CO2. The medium was replaced daily once the cells reached 50% confluency. When cells reached 80–90% confluency, they were passaged as clusters using Gentle Dissociation Reagent and replated onto fresh Vitronectin-coated wells in Essential 8 Medium with 2 nM Thiazovivin (STEMCELL Technologies, cat. no. 72254) ^48^.

### Differentiation of human iPSCs into human cortical neurons

After three days of pre-differentiation (Day 0), cells were dissociated with Accutase, resuspended in Classic Neuronal Medium, counted, and replated. Bio Coat Poly-D-Lysine-coated 96-well plates (Corning, cat. no. 356640) were seeded with 10,000 cells per well in 100 μL, on PDL-coated 35-mm dishes (iBidi, cat. no. 81148) with 100,000 cells in 1 mL, or on PDL-coated 6-well plates (Corning, cat. no. 351146) with 400,000 cells per well in 1 mL of medium. The medium consisted of a 1:1 mix of DMEM/F12 (Gibco, cat. no. 11320-033) and Neurobasal-A (Gibco, no. 10888-022), supplemented with 1× MEM Non-Essential Amino Acids, 0.5× GlutaMAX (Gibco, no. 35050-061), 0.5× N2 and B27 Supplements (Gibco, no. 17504-044), plus 10 ng/mL of each NT-3 and BDNF, 1 μg/mL mouse laminin, and 2 μg/mL doxycycline hydrochloride ^48^. On Day 7, half of the medium was replaced with fresh medium without doxycycline. From Day 14 onward, half-medium changes with fresh medium without doxycycline were performed weekly. Neurons were cultured for 40 days in vitro before use in experiments. At DIV40, all cultures were scraped, then fixed for regrowth measurements or MPS imaging at either 3 or 15 days post-injury (dpi). All cultures were regularly tested for mycoplasma contamination using the Lookout Mycoplasma PCR Detection Kit (Sigma-Aldrich, cat. no. MP0035).

### Generation of CRISPR interference-mediated knockdown of human iPSC lines

Human iPSC lines with ROCK-2 and PTEN CRISPRi knockdowns were generated using the CRISPRi-i3N parental line (WT), which stably expresses dCas9-KRAB under a doxycycline-inducible promoter ^48^. Two independent sgRNAs were designed to target the transcription start site of ROCK2 (sgRNA1, L1; sgRNA2, L2) or PTEN (sgRNA1, L1; sgRNA2, L2). The ROCK-2 sgRNA sequences are sgRNA1 (5’GCTGGCTGAGGGACTGCAAGG3’) and sgRNA2 (5’GGTGCGCCGGGAGCTGGCTGA3’). The PTEN sgRNA sequences are sgRNA1 (5’GTTGAGAGTTGAGCCGCTGTG3’) and sgRNA2 (5’GCAACTCTCAAACTTCCATCA3’). Both sgRNAs of ROCK-2 and PTEN were chosen from the CRISPick portal (https://portals.broadinstitute.org/gppx/crispick/public). These sgRNA sequences were cloned into a lentiviral backbone (see viral section below), lentiviral particles were produced, and iPSCs were transduced with puromycin selection. Knockdown efficiency was confirmed at both the mRNA level (via quantitative RT-PCR normalized to GAPDH) and the protein level (via immunofluorescence and western blot). WT-Scramble-2 iPSCs transduced with a non-targeting sgRNA served as controls. All CRISPRi lines were differentiated into cortical neurons using the previously described protocol.

### Differentiation of iPSCs into motor neurons

Human iPSC-derived motor neurons were generated using an NGN2-driven directed differentiation protocol ^49^. iPSCs harboring a doxycycline-inducible NGN2 transgene at the AAVS1 safe harbor locus (CRISPRi-i3N line, WTC11 background) were cultured in mTeSR1 medium (STEMCELL Technologies, cat. no. 85850) on Matrigel-coated plates (Corning, cat. no. 354277) at 37 °C with 5% CO2, with daily medium changes once cultures reached 50% confluence. On Day 0, iPSCs at 70–80% confluency was rinsed with DPBS, dissociated using StemPro Accutase (Thermo Fisher Scientific, cat. no. A1110501), centrifuged at 200 × g for 5 minutes at room temperature, and seeded at 4 × 10^6^ cells per 10-cm dish in mTeSR1 medium with 10 μM Y-27632 dihydrochloride (Tocris, cat. no. 1254) on Matrigel-coated plates. On Day 1, the medium was replaced with Induction Medium, which includes DMEM/F12 (Life Technologies, cat. no. 11320-033), 0.5× N2 supplement, 1× GlutaMAX, 0.1 mM MEM Non-Essential Amino Acids, 0.5% glucose, and 2 μg/mL doxycycline hydrochloride, supplemented with 10 μM SB431542 (Tocris, cat. no. 1614), 200 nM LDN193189 (Tocris, cat. no. 6053), 2 μM retinoic acid (RA; Tocris, cat. no. 0695), and 2 μM Smoothened agonist (SAG; Tocris, cat. no. 4366) to promote dual SMAD inhibition and motor neuron patterning. On Day 2, the medium was replaced with Induction Medium containing 5 μg/mL puromycin, 10 μM SB431542, 100 nM LDN193189, 1 μM RA, and 1 μM SAG. On Day 4, the medium was changed to Induction Medium with only 1 μM RA and 1 μM SAG.

On the 7^th^ day, cells were rinsed with DPBS, dissociated using Accutase, and replated at specified densities onto plates coated with poly-D-lysine and 10 μg/mL laminin. For 96-well plates, BioCoat PDL-coated plates (Corning, cat. no. 356640) were used. For STED imaging, 35-mm glass-bottom dishes were coated with 100 μg/mL poly-D-lysine for at least 1 hour at 37 °C, rinsed four times with sterile distilled water, and then coated with 10 μg/mL mouse laminin (Thermo Fisher Scientific, cat. no. 23017-015) in cold PBS for 1 hour at 37 °C. From Day 7 onward, cells were cultured in Neuronally Supportive Medium (NSM) containing Neurobasal medium (Life Technologies, cat. no. 21103-049), 2% (v/v) B27, 0.5% (v/v) N2, 1X GlutaMAX, 0.1 mM Non-Essential Amino Acids, and 0.5% glucose, freshly supplemented with 10 ng/mL BDNF (PeproTech, cat. no. 450-02), 10 ng/mL CNTF (PeproTech, cat. no. 450-13), and 10 ng/mL GDNF (PeproTech, cat. no. 450-10). During Days 7–8, 1 μg/mL laminin was added. Starting on Day 9, half-medium changes occurred three times weekly. Motor neurons were seeded at the same density as that of the human cortical neurons. Human motor neuron identity was confirmed via immunofluorescence co-labeling of βIII-tubulin with Islet-1, FOXP1, and ChAT at DIV32 ^49^. Cultures were maintained for 32 days before experiments; mechanical scrape injury was performed at DIV32, then fixed at either 3 or 8 days post-injury.

### Mechanical scrape injury model

Mechanical axon injury was induced using three methods, depending on the culture vessel and the experiments performed. In neuronal cultures on 35-mm glass-bottom dishes (used for STED MPS imaging), a sterile 200 μL pipette tip was drawn in a straight line across the monolayer to form a cell-free scraped zone about 200–500 μm wide. For cultures in 96-well plates (used for micro confocal imaging of axonal regrowth), a floating pin tool with FP1-WP pins (V&P Scientific) was used to scrape multiple wells in parallel. Injuries were performed at DIV10–11 for primary mouse cortical neurons, DIV40 for human iPSC-derived cortical neurons, and DIV32 for human iPSC-derived motor neurons. After injury, cultures were returned to the incubator and allowed to regenerate for 3, 8, or 15 days post-injury (dpi), depending on the experiment.

### Pharmacological treatments

The study utilized the following pharmacological agents: Y-27632 dihydrochloride (a ROCK inhibitor from STEMCELL Technologies, cat. no. 72302) and Fasudil hydrochloride (a ROCK inhibitor from Sigma-Aldrich, cat. no. CDS021620). Both Y-27632 and Fasudil hydrochloride were dissolved in sterile DMSO and used at 10 μM unless otherwise specified, within a dose–response range of 0–40 μM. Vehicle controls consisted of an equivalent volume of 0.1% DMSO. For ROCK-2 inhibition, the compounds were administered six hours after injury, and the cultures were maintained until fixation at designated post-injury time points.

### Immunofluorescence staining for ImageXpress-based automated imaging experiments and STED super-resolution confocal imaging experiments

Cultured mouse cortical neurons were fixed at room temperature (RT) for 20 minutes in a solution of 4% formaldehyde (Thermo Fisher Scientific, cat. no. 28906) and 4% D-sucrose (Thermo Fisher Scientific, cat. no. S5-500) in 1X PBS. For human iPSC-derived neurons (both cortical and motor), fixation involved 15 minutes at 37 °C with 4% paraformaldehyde (PFA) in DPBS. After fixation, samples were washed 3-4 times with 1X DPBS. Fixed samples were then blocked in a buffer containing 1% Bovine Serum Albumin (BSA, Sigma-Aldrich, cat. no. A4737) and 1% Triton X-100 (Thermo Scientific^TM^, cat. no. PI85111) for 1 hour at RT. Primary antibodies were incubated overnight at 4 °C in blocking buffer, followed by a 2–2.5-hour incubation with secondary antibodies at room temperature. Nuclei were counterstained with DAPI (1 μg/mL in DPBS) during the ImageXpress automated imaging protocol for 15 minutes at room temperature, then washed three times with DPBS. For STED imaging, nuclear staining was omitted. In 96-well plate imaging, 1X DPBS was added to each well, and plates were covered with aluminum foil. For 35 mm dishes used in STED imaging, ProLong Diamond served as the mounting medium.

### Fluorescence imaging in Image Express and image processing

For automated confocal imaging of intact and injured neuronal cultures in 96-well plates, fixed and immunostained plates were imaged using the ImageXpress® Micro Confocal system (Molecular Devices) with a 10X air objective. Three fluorescence channels were recorded at each site, such as FITC (βIII-tubulin for axons), Cy3 (Rhodamine-phalloidin for F-actin), and DAPI (nuclear stain). Nine overlapping sites per well were imaged to ensure complete coverage, including the scraped zone and nearby intact areas. Exposure, illumination, and focus settings were kept constant across all wells and plates. The images from the nine sites in each well were automatically stitched into a composite montage using a custom Journal script in MetaXpress® software (Molecular Devices) (https://www.moleculardevices.com). The resulting montage images were saved as TIFF files and imported into FIJI (ImageJ; NIH) (https://fiji.sc/) for further quantitative analysis.

### Quantification of axon regrowth

Axon regrowth was assessed by analyzing confocal images of βIII-tubulin immunostaining taken at 10× magnification with the ImageXpress® Micro Confocal system. A rectangular ROI was positioned within the scraped zone for each well. The βIII-tubulin fluorescence signal was thresholded uniformly across all images using Meta Xpress® software, and the percentage of the ROI area containing βIII-tubulin-positive signal was measured. This value, called “% or percentage area of regrowth,” was the main indicator of axon regrowth ability. All analyses were conducted blindly regarding experimental conditions, using FIJI/ImageJ or Meta Xpress®.

### Stimulated emission depletion (STED) super-resolution microscopy

Super-resolution confocal imaging was conducted using a Leica STELLARIS Tau STED microscope (Leica Microsystems CMS GmbH, Model DMi8 automated, SN 537859) (https://www.leica-microsystems.com). The setup included a pulsed white-light laser (WLL; tunable from 440 to 790 nm), a 660 nm STED depletion laser, and Power HyD detectors, all controlled with LAS X software. All STED imaging used an HC PL APO 100X/1.40 NA oil-immersion objective. For dual-color sequential STED imaging of βII-spectrin and F-actin, or MYO5A and ROCK-2, secondary antibodies conjugated with Alexa Fluor 555 (for βII-spectrin) were excited at 553 nm, while Oregon Green 488 (OG488) conjugates (for MYO5A or ROCK-2) or Oregon Green phalloidin (for F-actin) were excited at 488 nm using the WLL. The 660 nm pulsed STED depletion laser was applied to the Alexa Fluor 555 and OG488 channels to achieve resolution below the diffraction limit after their respective point-scanning confocal imaging. Dual-color images were acquired in sequential-imaging mode, with the OG488 and AF555 channels imaged in separate scans sharing the 660 nm depletion laser, ensuring channel separation and valid single-depletion-laser STED for both fluorophores. The detector pinhole was adjusted to 0.8 Airy units. Images were captured as 16-bit datasets at a single focal plane. All STED images were processed using the Leica Lightning adaptive deconvolution feature in LAS X software. This tool uses an iterative, hardware-adaptive algorithm to enhance signal-to-noise ratio and spatial resolution. Laser power, detector gain, pixel size, and line averaging remained consistent across all experimental conditions within each experiment.

### Regional sampling strategy for MPS analysis along regrowing axons

Non-overlapping regions of interest (ROIs) were defined along the regrowing axons depending on their distance from anatomical landmarks. In the intact proximal axon, two regions were designated as R1a (∼50 μm from the soma) and R1b (∼100 μm from the soma). In the regrowing axonal segments extending beyond the injury site, four regions were identified at progressively decreasing distances from the growth cone (GC) as R2 (∼300 μm), R3 (∼200 μm), R4 (∼100 μm), and R5 (∼50 μm).

### Autocorrelation and Fourier transformation analysis of βII-spectrin periodicity

Quantification of βII-spectrin periodicity, or ROCK-2/ MYO5A arrangements, was performed using 1D autocorrelation analysis of fluorescence intensity profiles extracted from STED images and processed in FIJI/ImageJ. For each axonal segment, a line profile of βII-spectrin fluorescence was extracted along the axon shaft. The autocorrelation function (ACF) was calculated using a custom Python script in Google Colab (https://colab.research.google.com/). The ACF amplitude was defined as the difference between its first peak and first trough, reflecting the ∼190 nm fundamental periodicity of the MPS. Segments were classified as having MPS-range periodicity if peaks occurred within 190–400 nm; segments without peaks in this range were assigned an ACF value of 0. This binary classification enabled the creation of stacked bar graphs depicting the proportion of periodic versus aperiodic segments across conditions. Fourier transform (FFT) analysis was also performed as a validation step, with peaks in the spatial frequency spectrum near ∼190 nm indicating periodic organization. All analyses were conducted in a blinded manner to the experimental conditions.

### Western blot analysis

Human or mouse cortical neurons were lysed using ice-cold 1× RIPA buffer (Millipore, cat. no. 20-188), supplemented with protease and phosphatase inhibitors (Halt™ Protease and Phosphatase Inhibitor Cocktail, EDTA-free, 100X, Thermo Fisher Scientific, cat. no. 78441). Protein concentration from the cell lysates was measured using the Pierce^TM^ BCA Protein Assay Kit (ThermoFisher Scientific, cat. no. 23225). For Western blotting, 10 μg of protein per lane was run on 7.5% SDS-PAGE gels (Bio-Rad), targeting βII-spectrin (∼250 kDa) and MYO5A (∼215 kDa). Proteins were transferred to nitrocellulose membranes with the Trans-Blot Turbo Transfer System (Bio-Rad) under semi-dry conditions. Membranes were blocked with 5% BSA in TBS-T (20 mM Tris, 150 mM NaCl, 0.1% Tween-20) for 1 hour at room temperature. Primary antibodies—mouse anti-βII-spectrin (BD Biosciences, 1:200), rabbit anti-MYO5A (Cell Signaling, 1:200), and mouse anti-β-actin (Sigma, 1:2,000)—were incubated overnight at 4°C in 5% BSA/TBS-T. After three washes, membranes were incubated for 1 hour at room temperature with secondary antibodies: donkey anti-mouse Alexa Fluor 555 dye and goat anti-rabbit Oregon Green 488. Bands were visualized using a ChemiDoc Imaging System (Bio-Rad), and band intensities were quantified using FIJI/ImageJ, with normalization to β-actin.

### Co-immunoprecipitation and mass spectrometry

Mass spectrometry following co-immunoprecipitation (CoIP) was used to identify the βII-spectrin interactome in human iPSC-derived cortical neurons under six different conditions, such as S1, WT uninjured + vehicle (0.1% DMSO); S2, WT uninjured + 10 μM Y-27632; S3, WT injured + vehicle; S4, WT injured + 10 μM Y-27632; S5, ROCK-2-CRISPRi-KD uninjured; S6, ROCK-2-CRISPRi-KD injured. The CoIP employed antibodies against βII-spectrin (BD Biosciences, cat. no. 612563; 10μg) and αII-spectrin (Thermo Fisher Scientific, cat. no. PA5-44905; 10μg), with mouse or rabbit IgG (Santacruz Biotechnology, cat. no. sc-02025; 10μg) isotype controls as negative controls. Cell lysates (250-500 μg protein per condition) were pre-cleared and incubated with antibody-bound protein G magnetic Dynabeads (Thermo Fisher Scientific, cat no. 10007D) for 2 hours at 4°C with rotation, then washed four times with lysis buffer (50 mM Tris pH 8.0, 150 mM NaCl, 1% Triton X-100, protease inhibitors). Bound proteins were eluted using Laemmli buffer at 95°C for 5 minutes. The eluates were subjected to SDS-PAGE, and gel slices were excised for mass spectrometry. Protein abundance data were visualized as a z-score heatmap through unsupervised hierarchical clustering. GO enrichment analysis of the identified proteins was performed using the DAVID database, and the STRING platform was employed for protein–protein interaction network analysis to reveal interactions among candidate interactors.

### Mass spectrometry sample preparation, data acquisition, and analysis

Gel slices were cut into small pieces and washed with 1 mL water for 10 min on a tilt table, followed by washing with 1 mL 50% acetonitrile (ACN)/100 mM ammonium bicarbonate (ABC) for 20 min. Samples were reduced with 1 gel volume of 4.5 mM dithiothreitol (DTT) in 100 mM ABC at 37 °C for 20 min. After removal of the DTT solution and cooling, samples were alkylated with 1 gel volume of 10 mM iodoacetamide (IAA) in 100 mM ABC for 20 min at RT in the dark. Gel pieces were washed twice with 1 mL 50% ACN/100 mM ABC and twice with 1 mL 50% ACN/25 mM ABC for 10 min each, then briefly dried by SpeedVac. Gel pieces were resuspended in 1 gel volume of 25 mM ABC containing 2.5 ng/μL sequencing-grade trypsin (Promega, V5111) and incubated at 37 °C for 16 h. Supernatants containing tryptic peptides were transferred to new tubes, and residual peptides were extracted from gel pieces with 3 gel volumes of 80% ACN/0.1% trifluoroacetic acid (TFA) for 15 min, pooled with the initial digests, and dried by SpeedVac. Dried peptides were dissolved in 24 μL MS loading buffer (2% ACN, 0.2% TFA), with 5 μL injected for LC-MS/MS analysis.

Data were acquired on a Thermo Scientific Q Exactive Plus coupled to a Waters nanoACQUITY UPLC system with a binary solvent system (A: 100% water, 0.1% formic acid; B: 100% ACN, 0.1% formic acid). Trapping was performed at 5 μL/min in 99.5% Buffer A for 3 min using a Waters Symmetry® C18 180 μm × 20 mm trap column. Peptides were separated on an ACQUITY UPLC PST (BEH) C18 nanoACQUITY Column (1.7 μm, 75 μm × 250 mm; 45 °C) at 300 nL/min with a gradient from 3% to 90% Buffer B over 180 min. MS spectra were acquired in profile mode over 300–1,700 m/z (70,000 resolution, AGC target 3 × 10^6^, 45 ms maximum injection time). Data-dependent MS/MS was performed on the top 20 precursors per MS scan (17,500 resolution, AGC target 1 × 10^5^, 100 ms maximum injection time, 1.7 m/z isolation window, HCD fragmentation at normalized collision energy 28%, charge states 2–6, intensity threshold 1 × 10^4^, 30 s dynamic exclusion).

All MS/MS spectra were analyzed using Proteome Discoverer v2.5.0.400 (Thermo Scientific) with the Mascot search algorithm v2.8.3 (Matrix Science). Data were searched against the Swiss-Prot database restricted to *Homo sapiens* (version 2024_2; 20,435 sequences). Search parameters included: trypsin digestion with up to 2 missed cleavages; 10 ppm peptide mass tolerance; 0.02 Da MS/MS fragment tolerance; fixed modification of cysteine carbamidomethylation; variable modification of methionine oxidation. Target-decoy database searches were conducted with 95% confidence threshold (*p* < 0.05). Scaffold v5.1.2 (Proteome Software) ( https://www.proteomesoftware.com/products/scaffold-5) was used to validate for MS/MS-based peptide and protein identification. Peptide identifications were accepted at >95.0% probability (Scaffold Local FDR algorithm). Protein identifications required >99.0% probability and ≥2 identified peptides. Proteins sharing similar peptides were grouped to satisfy parsimony principles.

### Primary antibodies used for Imaging, Western blot, and Co-immunoprecipitation experiments

This study employed the following primary antibodies: Mouse anti-βII-spectrin (BD Transduction Laboratories, cat. no. 612563; 1:200 for immunofluorescence; 1:500 for western blot; 10μg for CoIP), Mouse anti-αII-spectrin (Bio Legend; cat. no. 803201; 1:200 for immunofluorescence), Mouse anti-αII-spectrin (Thermo Fisher Scientific, cat. no. PA5-44905; 1:500 for western blot; 10μg for CoIP), Mouse anti-βIII-tubulin (Promega; cat. no. G7121; 1:1,000 for immunofluorescence), Phalloidin-TRITC (ThermoFisher Scientific; cat. no. R415; 1:1,000 for immunofluorescence), Rabbit anti-ROCK-2 middle polyclonal antibody (Proteintech; cat. no. 21645-1-AP; 1:300 for immunofluorescence), Rabbit anti-MYO5A (Cell Signaling Technology; cat. no. 3402; 1:100 for immunofluorescence; 1:1,000 for western blot), Oregon GreenTM 488 Phalloidin (ThermoFisher Scientific; cat. no. O7466; 1:500 for immunofluorescence), Rabbit anti-HA tag (Cell Signaling Technology; cat. no. 3724; 1:1,000 for immunofluorescence), Rabbit anti-Islet-1 (Abcam; cat. no. ab178400; 1:300 for immunofluorescence of human iPSC derived motor neurons), Rabbit anti-FOXP1 (Abcam; cat. no. ab16645; 1:500 for immunofluorescence of human iPSC derived motor neurons), Rabbit anti-ChAT (Sigma Aldrich Millipore; cat. no. AB144P; 1:150 for immunofluorescence of human iPSC-derived motor neurons), Anti-β-actin (Sigma Aldrich Millipore; cat. no. A5441; 1:5,000 for western blot loading control), Chicken anti-MAP-2 (Abcam; cat. no. ab5392; 1:500 for immunofluorescence) and IgG (Santacruz Biotechnology, cat. no. sc-02025; 10μg for CoIP).

### Secondary antibodies for immunofluorescence and western blot experiments

The study used several secondary antibodies: Goat anti-mouse IgG Alexa Fluor 488 (Thermo Fisher Scientific, cat. no. A-11001; 1:1,000 for immunofluorescence in mice and 1:500 for immunofluorescence in human; 1:500 for western blot), Donkey anti-mouse IgG (H+L) Alexa Fluor 555 (Thermo Fisher Scientific; cat. no. A-31570; 1:500 for immunofluorescence; 1:500 for western blot), Donkey anti-rabbit IgG (H+L) Alexa Fluor 555 (Thermo Fisher Scientific; cat. no. A-31572; 1:500; 1:500 for western blot), Goat anti-rabbit IgG (H+L) Oregon Green 488 (Thermo Fisher Scientific, cat. no. O-11038; 1:500), and Donkey anti-rabbit IgG (H+L) Alexa Fluor 647 (Thermo Fisher Scientific; cat. no. A-31573; 1:1,000).

### AAV and lentiviral transduction

For MYO5A knockdown in CRISPRi-i3N iPSCs, AAV vectors encoding MYO5A-targeting sgRNAs under a U6 promoter and mTagBFP2^50^ under a CAG promoter (pAAV CAG>mTagBFP2; U6>[SgRNA]; 6,004 bp each) (Supplementary Fig. 10a-b). Two independent sgRNA sequences were used: sgRNA1, 5′-GGGCGGCCGCCCGAGCGGACT-3′, and sgRNA2, 5′-GGGCCTGGGCGGCCGCCCGAG-3′. Both constructs were flanked by AAV inverted terminal repeats (ITRs). The AAV constructs were designed and obtained from Vector Builder (https://en.vectorbuilder.com/). HEK293T cells were seeded at 1 × 10⁷ cells per 15-cm dish one day prior to transfection and were transfected upon reaching approximately 80% confluency. For AAV production, a plasmid cocktail comprising the pDF6 helper plasmid (18 µg; Addgene, cat. no. 112867), pAAV-CAG driving sgRNAs targeting *MYO5A* (6 µg), and the Rep/Cap packaging plasmid pAAV2/9n (6 µg; Addgene, cat. no. 112865) was prepared in 3 ml of serum-free DMEM. Polyethylenimine (PEI; Polysciences, cat. no. 49553-93-7) was added to a final volume of 150 µl, and the DNA–PEI mixture was incubated for 15 min at room temperature before being applied to the cells. Cells were harvested 96 h after transfection and treated with DNase I (10 U/ml; Sigma-Aldrich, AMPD1) and Benzonase (50 U/ml; Millipore, cat. no. 70746) for 40 min at 37 °C. Cell debris was pelleted by centrifugation at 3,000 × g for 20 min at 4 °C, and the supernatant containing viral particles was further purified by ultracentrifugation over an iodixanol density gradient ^51^. AAV transduction of WT human iPSC-derived cortical neurons was carried out at DIV7 or DIV14; for ROCK-2 CRISPRi KD lines, transduction was performed at DIV14, as specified for each experiment.

For the MYO5A overexpression experiment, two lentiviral vectors were utilized: a control virus expressing mCherry under the EF1A promoter [pLV(Exp)-EGFP/Puro-EF1A-mCherry; 10,462 bp], and a MYO5A-overexpressing virus that encodes MYO5A isoform 1 fused to a 3X HA tag under the neuron-specific SYN1 promoter [pLV(Exp)-SYN1-(MYO5A-Iso-1)/3×GGGGS/HA/HA/HA; 12,729 bp]. The MYO5A-OE lentiviral construct was designed and obtained as a lentiviral vector from Vector Builder. Lentiviral transduction was carried out at DIV14. Successful transduction was confirmed by TagBFP2 fluorescence (for AAV transduction) or mCherry/HA tag (for lentiviral transduction) immunofluorescence at fixation. Knockdown efficiency was evaluated through immunofluorescence quantification of MYO5A normalized to βII-spectrin in transduced versus non-transduced neurons. Overexpression was validated using anti-HA immunostaining (HA-Tag Rabbit Monoclonal Antibody, Cell Signaling Technology, cat. no. 3724; 1:800).

### Lentiviral packaging and transduction for CRISPRi knockdown of ROCK-2 and PTEN

HEK293FT cells were cultured in a six-well plate for a day prior to transfection. The following day, the cells were transfected with lentiviral vectors encoding sgRNAs targeting either ROCK-2, PTEN, or Scramble-2 (as a control). Equal amounts of lentiviral packaging vectors (Thermo Fisher, cat. no. K497500), facilitated by JetPRIME reagent (Polyplus), were used. After six hours of incubation, the culture medium was replaced, and then the cells were again incubated for an additional thirty-six hours. The supernatants were then harvested and filtered through a 0.2-micron filter unit. The supernatants contain the lentiviral particles. For viral transduction, CRISPRi-i3N iPSCs (WT, stably expressing doxycycline-inducible dCas9-KRAB) were plated on Vitronectin-coated plates in Essential 8 Medium with 2 nM Thiazovivin and 5 µg/mL Polybrene (Sigma Aldrich, cat. no. TR-1003). The cells were then infected with clarified lentiviral supernatant encoding ROCK-2- or PTEN-targeting sgRNAs. The following day, the virus-containing medium was replaced with fresh Essential 8 Medium. Starting 48 hours post-transduction, stably transduced cells were selected with 1 µg/mL puromycin (Gibco, cat. no. A1113803) for 5-7 days. The surviving puromycin-resistant cells were expanded to generate polyclonal CRISPRi knockdown (KD) lines. Controls included CRISPRi-i3N iPSCs transduced with a non-targeting (Scramble-2) sgRNA lentivirus (WT-Scramble-2). Knockdown efficiency was confirmed at the mRNA level via quantitative RT-PCR (normalized to GAPDH) and at the protein level by immunofluorescence. All ROCK-2-CRISPRi KD and PTEN-CRISPRi KD lines, along with the WT-Scramble-2 control, were then differentiated into cortical neurons following the previously described protocol ^48^.

### Quantitative reverse-transcription PCR (qRT-PCR)

Total RNA was extracted from human iPSC-derived cortical neurons using the RNeasy Plus Mini Kit (Qiagen, cat. no. 74136). RNA concentration was quantified by NanoDrop spectrophotometry. cDNA was synthesized from 500 ng total RNA using the iScript Reverse Transcription Supermix (Bio-Rad, cat. no. 1708840). Quantitative real-time PCR was performed on a CFX96 Real-Time PCR system (Bio-Rad) using the Universal SYBR Green Supermix (Bio-Rad, cat. no. 1725272). Gene expression levels were calculated using the 2^−ΔΔCt^ method and normalized to GAPDH as the reference gene. PCR was conducted for 40 cycles at 95 °C for 30 s (denaturation), 55 °C for 30 s (annealing), and 72 °C for 30 s (extension). ROCK-2 mRNA knockdown was confirmed in both L1 and L2 CRISPRi KD lines relative to WT-Scramble-2 controls.

### Axon diameter measurement

Axon diameter was measured in intact, uninjured axonal segments roughly 50 μm from the cell body in human iPSC-derived cortical neurons. These neurons were transduced with MYO5A-targeting sgRNA AAVs or MYO5A-overexpression lentivirus. Measurements were taken from STED confocal images of βII-spectrin immunostaining at 100X magnification, using FIJI/ImageJ to manually draw line profiles perpendicular to the axon shaft and measure the full-width at half-maximum of the βII-spectrin signal. A minimum of 20 axons were measured per condition across three independent experiments.

### Growth cone imaging

To evaluate MYO5A localization at growth cones, regrowing axons from human iPSC-derived wild-type cortical neurons were immunostained for MYO5A (Alexa Fluor 555; 1:300) and F-actin (Oregon Green 488 - phalloidin; 1:200). High-resolution confocal images were obtained at 100X magnification using a Leica STELLARIS 8 STED system. The colocalization of MYO5A with F-actin in filopodia and lamellipodia was assessed by visual inspection and validated using Pearson’s correlation analysis within manually selected growth-cone regions of interest.

### Statistical analysis

All statistical analyses were performed using GraphPad Prism (version 10.5) (https://www.graphpad.com/). Data are presented as mean ± SEM, with individual data points shown. For comparisons between two groups, unpaired two-tailed Student’s *t*-tests were used. For comparisons among three or more groups, a one-way ANOVA with Tukey’s post hoc test was used. Significance thresholds were set at *P* ≤ 0.05 (*), *P* ≤ 0.01 (**), *P* ≤ 0.001 (***), and *P* ≤ 0.0001 (****). All experiments were performed with at least three independent biological replicates (*N* ≥ 3 independent batches/experiments). Sample sizes (number of axonal segments, wells, or mice) are reported in individual figure legends. Statistical tests are indicated throughout the text and figure legends. Investigators were blinded to conditions during image acquisition and during data analyses.

### Reproducibility

All experiments were independently replicated at least three times using biologically independent batches of neurons or mice. Representative images shown in all figures were selected from experiments replicated with similar results across independent replicates. No data were excluded from the analysis.

## Supporting information

Supplementary Figures 1-10 and Legend for XLXS files

## ACKNOWLEDGEMENTS

We thank Ines Ingabire for her help with maintaining human cortical neuronal cultures and for assisting with the qPCR method for the CRISPRi line. We thank Koustuv Sinha for his guidance in developing Python code in Colab for Fourier transform and autocorrelation analyses. We recognize Sarah Helena Nies for her guidance in maintaining iPSC cultures. We appreciate Ramakrishnan Kannan for providing reagents and assisting with AAV transfection. We thank Olivia Wan for carefully proofreading the manuscript. This study was supported by NIH grant R35NS097283 awarded to S.M.S.

## AUTHOR CONTRIBUTIONS

Conceptualization, A.B, and S.M.S.; Methodology, A.B., E.M.H., N.K., K.J., T.A., J.K., L.T. and S.M.S.; Investigation, A.B. and E.M.H.; Data Analysis, A.B., and J.K.; Writing—Original Draft, A.B., K.J. and S.M.S.; Writing—Review & Editing, all; Funding Acquisition, S.M.S.; Resources, S.M.S; Supervision, S.M.S.

## COMPETING INTERESTS

The authors declare no competing interests.

## Notes

### Competing Interest Statement

The authors have declared no competing interest.

