## Supplementary Figures 1-10 and Legend for XLXS files for "Myosin 5A mediates membrane-associated periodic skeleton reassembly during axon regeneration in response to ROCK-2 inhibition"

**SUPPLEMENTARY INFORMATION**

- **Supplementary Figures 1-10**
- **Legends for Supplementary Data Files 1-3 (XLSX format)**

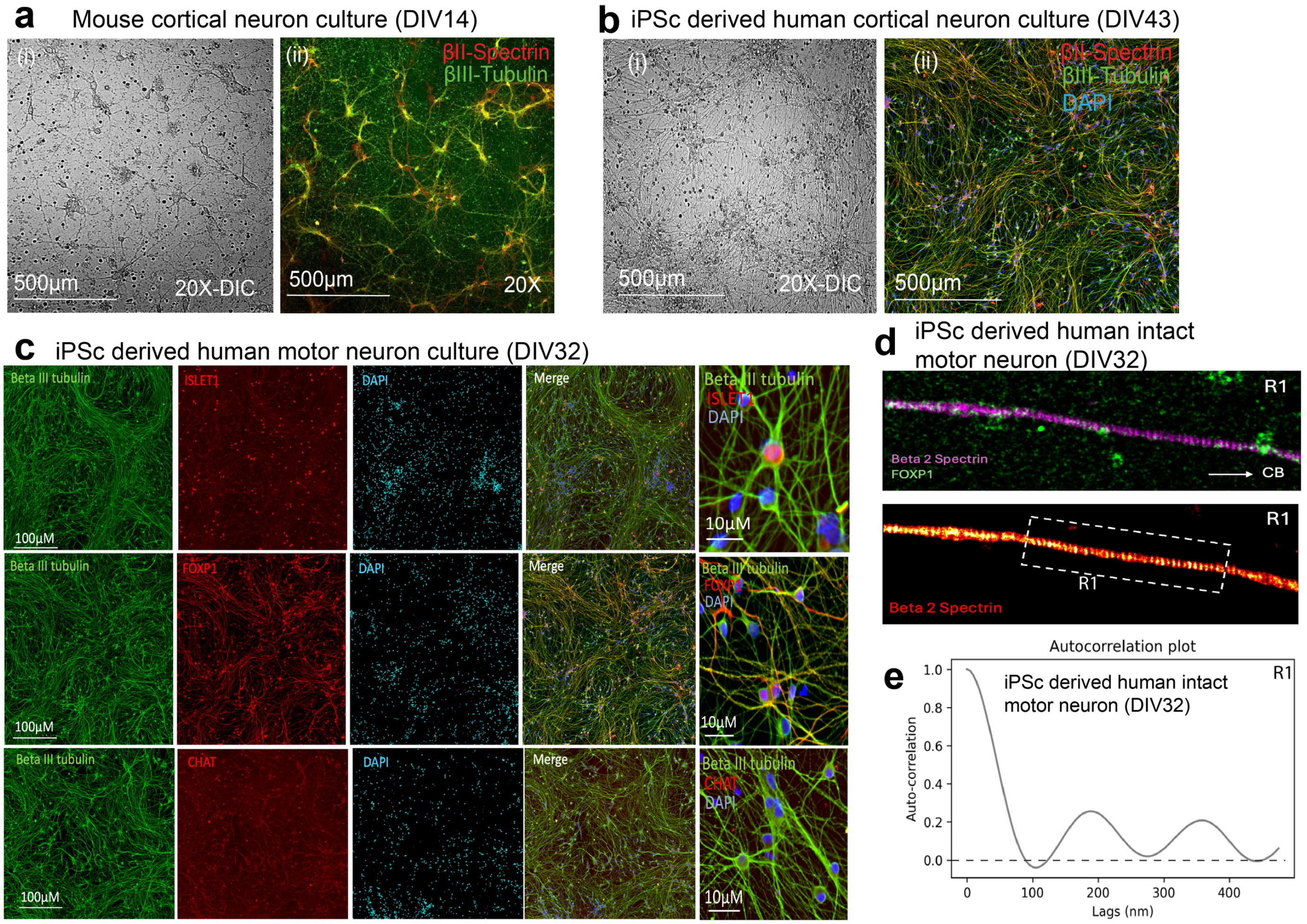


**Supplementary Figure 1: Periodic arrangement of βII-spectrin in intact axons of mouse and human neurons.**

**a.** Representative images of primary mouse cortical neuron cultures at DIV14. (i) DIC image at 20X magnification showing neuronal morphology and network density. (ii) Corresponding 10X fluorescent confocal image immunostained for βII-spectrin (red) and βIII-tubulin (green), demonstrating extensive axonal networks with co-expression of both cytoskeletal markers. Scale bars, 500 μm.

**b**. Representative images of human iPSC-derived cortical neuron cultures at DIV43. (i) DIC image at 20X magnification. (ii) Corresponding 10X fluorescent confocal image immunostained for βII-spectrin (red), βIII-tubulin (green), and DAPI (cyan), confirming mature neuronal identity and dense axonal projections. Scale bars, 500 μm.

**c**. Characterization of human iPSC-derived motor neuron cultures at DIV32. Left panels showing confocal images of neurons immunostained for βIII-tubulin (green) and the motor neuron markers Islet-1 (red, top row), FOXP1 (red, middle row), and ChAT (red, bottom row), together with DAPI (cyan) and merged overlays. Right panels showing higher-magnification views of individual motor neurons co-expressing βIII-tubulin with Islet-1, FOXP1, or ChAT, confirming motor neuron identity. Scale bars for left panels 100 μm; for right panels, 10 μm.

**D**. Representative STED super-resolution images of an intact human iPSC-derived motor neuron at DIV32. The top panel shows an overview of the neuron immunostained for βII-spectrin (green) and FOXP1 (magenta), with the cell body (CB) and axonal region R1 indicated. Bottom panels show magnified views of the R1 axonal region, revealing the periodic arrangement of βII-spectrin along the axon shaft visualized by super-resolution microscopy.

**e**. Autocorrelation analysis of βII-spectrin fluorescence intensity along the axon in region R1 of an intact human iPSC-derived motor neuron at DIV32. The autocorrelation function displays a periodic arrangement with a spacing of approximately 190 nm, consistent with the characteristic periodicity of the membrane-associated periodic skeleton (MPS).

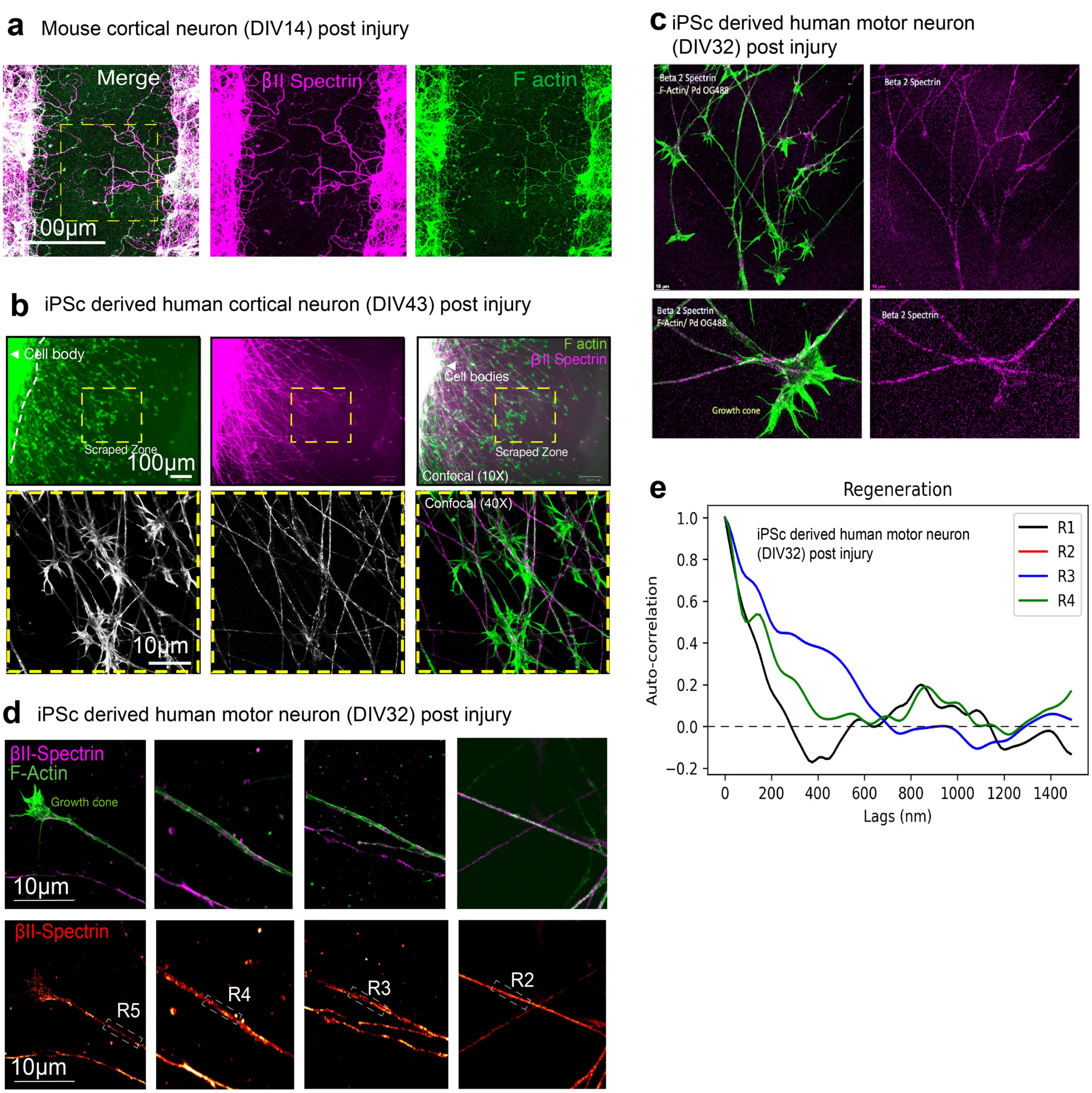


**Supplementary Figure 2: Characterization of βII-spectrin and F-actin distribution in regrowing axons following mechanical or scrape injury in mouse and human cortical and motor neurons.**

**a.** Representative confocal images of primary mouse cortical neurons at DIV14 following scrape injury, immunostained for βII-spectrin (magenta) and F-actin (green), with merged overlay (left). The dashed yellow rectangle indicates the region at the border of the scraped zone where regrowing axons are visible. Dashed yellow rectangles represent areas shown at higher magnification. Scale bar, 100 µm.

**b.** Representative confocal images of human iPSC-derived cortical neurons at DIV43 post-injury. The top row shows a 10× overview of the injury site showing βII-spectrin (magenta), F-actin (green), and a merged overlay, with the scraped zone and cell body regions indicated. Dashed yellow rectangles represent areas shown at higher magnification. The bottom row shows higher-magnification (40×) confocal images of the boxed regions, revealing individual regrowing axons co-stained for βII-spectrin (white, left), F-actin (green, right), and merged (magenta/green). Scale bars: top, 100 μm; bottom, 10 μm.

**c.** Representative confocal images of human iPSC-derived motor neurons at DIV32 post-injury. Top panels show neurons co-stained (left) for βII-spectrin (magenta) and F-actin (green) and βII-spectrin alone (magenta, right). Bottom panels show regrowing motor neurons with prominent growth cones, co-stained for βII-spectrin and F-actin (left) and βII-spectrin alone (right). Growth cones are enriched in F-actin, whereas βII-spectrin distributes along the axon shaft.

**d.** STED super-resolution images of a regrowing human iPSC-derived motor neuron axon at DIV32 post-injury. The top rows show co-immunostaining for βII-spectrin (magenta) and F-actin (green) at progressively greater distances from the growth cone. Bottom row shows βII-spectrin signal visualized in a fire/heat lookup table, with sequential axonal regions R2–R5 indicated by dashed boxes at increasing distances from the growth cone. The periodic arrangement of βII-spectrin becomes progressively more apparent in regions distal to the growth cone. Scale bars, 10 μm.

**e**. Autocorrelation analysis of βII-spectrin fluorescence intensity along regrowing motor neuron axons at regions R1 (black), R2 (red), R3 (blue), and R4 (green), corresponding to increasing distances from the growth cone. The dashed line indicates the zero-correlation baseline.

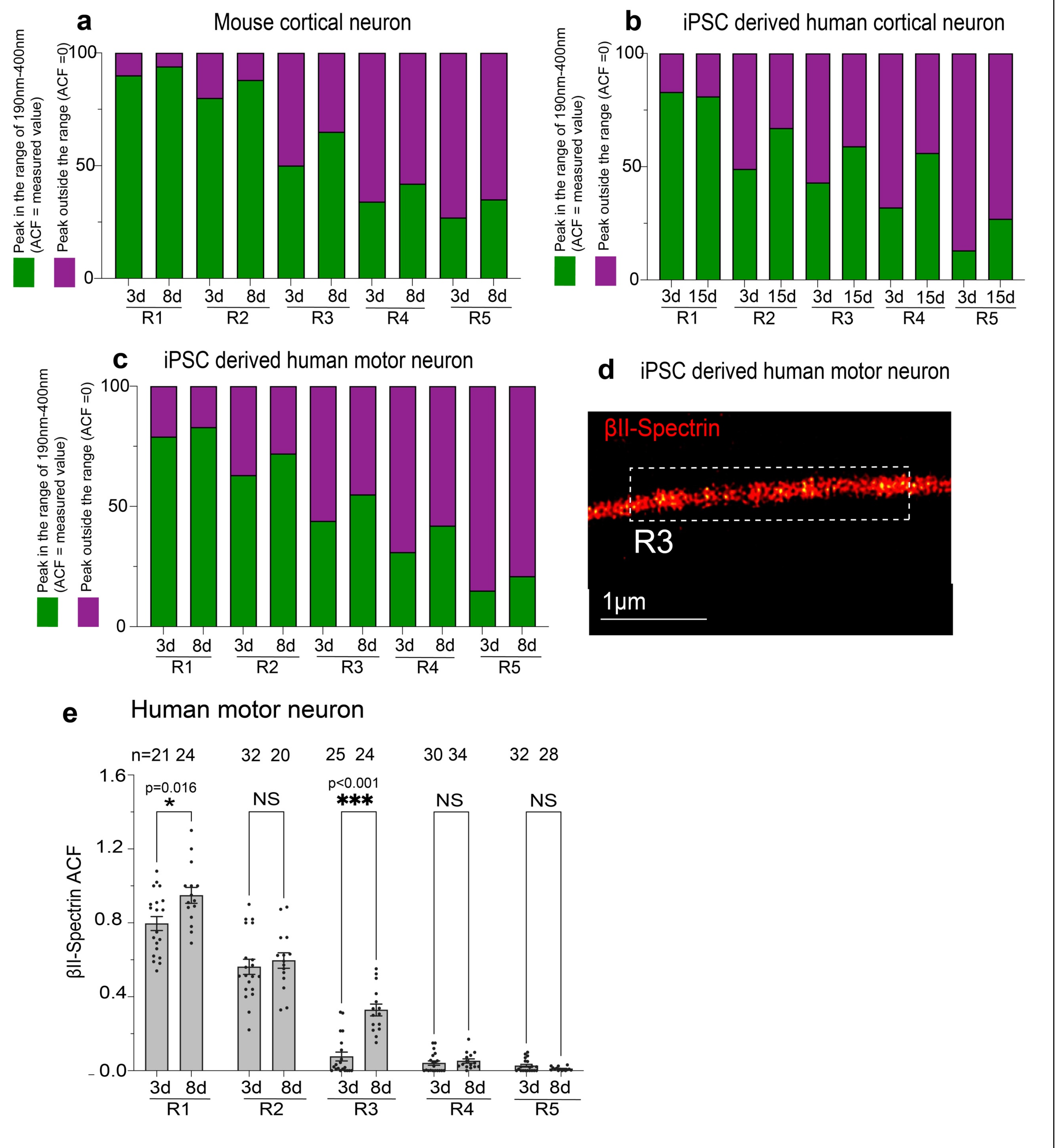


**Supplementary Figure 3: Spectrin periodicity in regenerating axons recovers partially at 8–15 days post-injury in human motor neurons.**

**a.** Stacked bar graphs showing the percentage of axonal segments with autocorrelation function (ACF) peaks in the 190–400 nm range (green, ACF = measured value, indicating periodic βII-spectrin organization) versus peaks outside this range (purple, ACF = 0, indicating loss of periodicity) in mouse cortical neurons at 3 days (3d) and 8 days (8d) post-injury across five axonal regions (R1–R5).

**b.** Stacked bar graphs as in (**A**) but for iPSC-derived human cortical neurons at 3 days (3d) and 15 days (15d) post-injury across R1–R5. A similar pattern of partial recovery is observed, with the proportion of periodic segments increasing between 3d and 15d in R2 and R3, while distal regions R4 and R5 show minimal recovery.

**c.** Stacked bar graphs as in (**A**) but for iPSC-derived human motor neurons at 3 days (3d) and 8 days (8d) post-injury across R1–R5. Partial recovery of periodicity is observed in R1–R3 by 8d, while R4 and R5 remain predominantly aperiodic.
**d.** Representative STED super-resolution image of βII-spectrin (red) in an iPSC-derived human motor neuron axon at region R3. The dashed box indicates the region used for autocorrelation analysis. Scale bar: 1 µm.
**e.** Quantification of βII-spectrin ACF (autocorrelation amplitude) across five axonal regions (R1–R5) in human motor neurons at 3 days (3d, black dots) and 8 days (8d, green dots) post-injury. Sample sizes (number of axonal segments): R1 ($n$ = 21, 24), R2 ($n$ = 32, 20), R3 ($n$ = 25, 24), R4 ($n$ = 30, 34), R5 ($n$ = 32, 28). βII-spectrin ACF significantly increased between 3d and 8d at R1 (P = 0.016) and R3 (P < 0.001), but not at R2, R4, or R5 (NS). Data are presented as mean ± SEM with individual data points shown. All experiments were conducted in $N$ = 3 independent replicates. Statistics: unpaired two-tailed Student's $t$-test for each region. NS, not significant; *P < 0.05; ***P < 0.001. P-values are indicated on the graph.

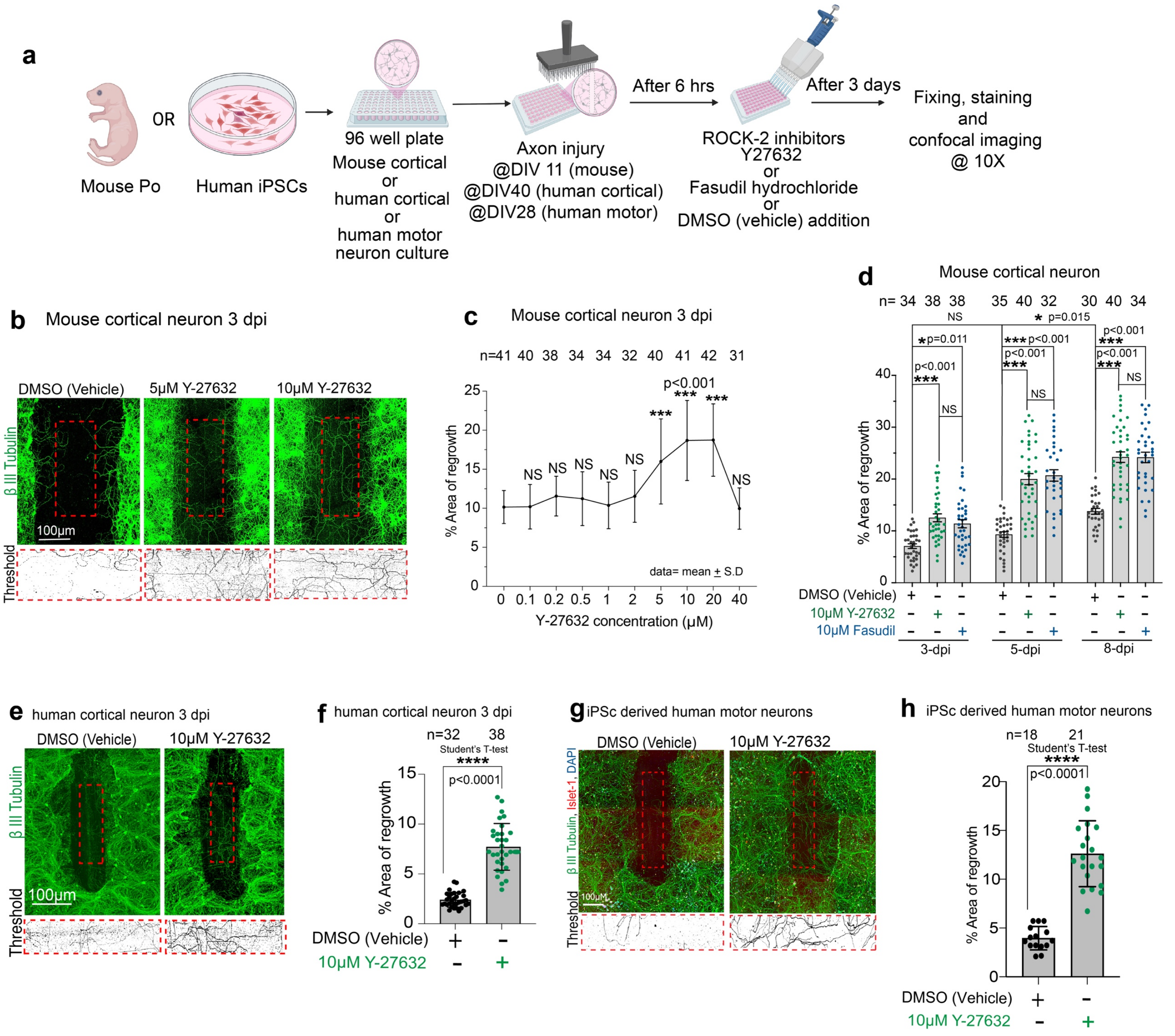


**Supplementary Figure 4: Pharmacological inhibition of ROCK-2 enhances overall axon regrowth in mouse and human cortical neurons.**
**a.** Schematic of the experimental workflow. Mouse P0 cortical neurons, human iPSC-derived cortical neurons, or human iPSC-derived motor neurons were cultured in 96-well plates. Neurons were subjected to axon injury

at DIV 11 (mouse), DIV 40 (human cortical), or DIV 28 (human motor). Six hours post-injury, the ROCK-2 inhibitors Y-27632 or Fasudil hydrochloride, or DMSO vehicle, were added. Neurons were fixed 3 days post-injury for immunostaining and 10X confocal imaging.

**b.** Representative confocal images of βIII-tubulin-stained mouse cortical neurons at 3 days post-injury (dpi) treated with DMSO vehicle (left), 5 μM Y-27632 (middle), or 10 μM Y-27632 (right). Red dashed boxes indicate regions of interest at the injury site. Scale bar: 100 μm.

**c.** Dose-response quantification of percent area of regrowth in mouse cortical neurons at 3 dpi treated with increasing concentrations of Y-27632 (0, 0.1, 0.2, 0.5, 1, 2, 5, 10, 20, and 40 μM). Sample sizes (number of wells analyzed from 96-well plates): 0 μM (*n* = 41), 0.1 μM (*n* = 40), 0.2 μM (*n* = 38), 0.5 μM (*n* = 34), 1 μM (*n* = 34), 2 μM (*n* = 32), 5 μM (*n* = 40), 10 μM (*n* = 41), 20 μM (*n* = 42), 40 μM (*n* = 31). Significant increases in regrowth were observed at 10 μM and 20 μM ($P < 0.001$ for both), whereas concentrations of 0.1-5 μM and 40 μM showed no significant effect relative to the vehicle. Data are presented as mean ± SD. Statistics: one-way ANOVA with Tukey's *post hoc* test for multiple comparisons.

**d.** Quantification of percentage area of regrowth in mouse cortical neurons treated with DMSO vehicle, 10 μM Y-27632, or 10 μM Fasudil at 3, 5, and 8 dpi. Sample sizes (each well of 96-well plate): 3 dpi (*n* = 34, 38, 38), 5 dpi (*n* = 35, 40, 32), 8 dpi (*n* = 30, 40, 34). At 3 dpi, Y-27632 significantly increased regrowth compared to vehicle ($P = 0.011$), while Fasudil showed no significant difference from vehicle; Y-27632 and Fasudil were not significantly different from each other. At 5 dpi, both Y-27632 ($P < 0.001$) and Fasudil ($P = 0.015$) significantly increased regrowth compared to vehicle; Y-27632 and Fasudil were not significantly different. At 8 dpi, both Y-27632 ($P < 0.001$) and Fasudil ($P < 0.001$) significantly increased regrowth compared to vehicle; Y-27632 and Fasudil did not differ significantly. Data are presented as mean ± SEM with individual data points shown. Statistics: one-way ANOVA with Tukey's *post hoc* test for multiple comparisons at each time point.

**e.** Representative confocal images of βIII-tubulin-stained human iPSC-derived cortical neurons at 3 dpi treated with DMSO vehicle (left) or 10 μM Y-27632 (right), with corresponding thresholded binary images for βIII-tubulin are shown below. Red dashed boxes indicate regions of interest. Scale bar: 100 μm.

**f.** Quantification of the percentage area of regrowth in human cortical neurons. Sample sizes: vehicle (*n* = 32), 10 μM Y-27632 (*n* = 38). Treatment with 10 μM Y-27632 significantly increased regrowth compared to vehicle ($P < 0.0001$). Data are presented as mean ± SEM with individual data points shown. Statistics: unpaired two-tailed Student's *t*-test.

**g.** Representative confocal images of human iPSC-derived motor neurons at 3 days post injury stained for βIII-tubulin (green), Islet-1 (red), and DAPI (blue), treated with DMSO vehicle (left) or 10 μM Y-27632 (right). Corresponding thresholded images for βIII-tubulin are shown below. Red dashed boxes indicate regions of interest. Scale bar: 100 μm.

h. Quantification of the percentage area of regrowth in human iPSC-derived motor neurons. Sample sizes: vehicle (*n* = 18), 10 μM Y-27632 (*n* = 21). Treatment with 10 μM Y-27632 significantly increased regrowth compared to vehicle ($P< 0.0001$). Data are presented as mean ± SEM with individual data points shown. Statistics: unpaired two-tailed Student's *t*-test. All experiments were conducted in *N* = 3 independent replicates. NS, not significant; *$P < 0.05$; ***$P < 0.001$; ****$P < 0.0001$. P-values are indicated on the graphs.

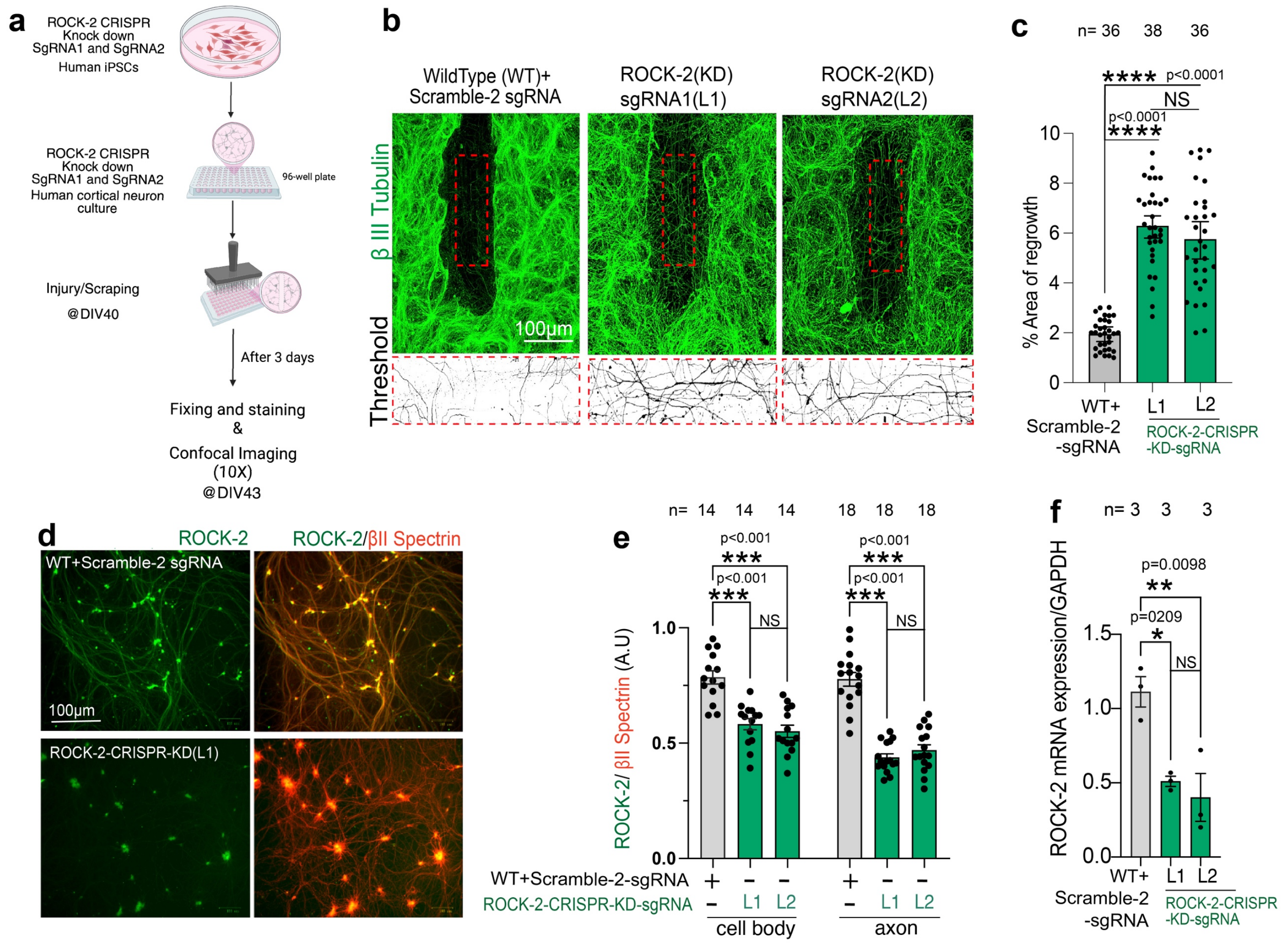


**Supplementary Figure 5: CRISPR interference of ROCK-2 enhances overall axon regrowth in human cortical neurons.**

**a.** Representative confocal images of wild-type (top) and ROCK-2 KD sgRNA1 (L1, bottom) human cortical neurons immunostained for ROCK-2 (red, left) and co-stained for ROCK-2 and βII-spectrin (right, merged). ROCK-2 protein expression is visibly reduced in the knockdown line compared to WT in both cell bodies and axons. Scale bar: 100 µm.

**b.** Quantification of ROCK-2 fluorescence intensity normalized to βII-spectrin intensity in cell bodies (left) and axons (right) of wild-type (WT+Scramble-2 sgRNA) and ROCK-2 KD lines (ROCK-2-CRISPR-sgRNA1; L1 and sgRNA2; L2) neurons. Sample sizes: $n$ = 14 per group for cell bodies and $n$ = 18 for axons. ROCK-2 KD neuron lines showed significantly reduced ROCK-2 protein levels in both cell bodies ($P < 0.01$) and axons ($P < 0.001$) compared to WT. Statistics: one-way ANOVA with Tukey's *post hoc* test for multiple comparisons. All experiments were conducted in $N$ = 3 independent replicates. NS, not significant; **P < 0.01; ***P < 0.001; ****P < 0.0001. P-values are indicated on the graphs.

**c.** Quantification of ROCK-2 mRNA expression normalized to GAPDH in wild-type (WT+Scramble-2 sgRNA), ROCK-2 KD Line 1 (ROCK-2-CRISPR-sgRNA1), and ROCK-2 KD Line 2 (ROCK-2-CRISPR-sgRNA2) neurons. $N$ = 3 independent replicates per group. Both Line 1 ($P < 0.01$) and Line 2 ($P < 0.001$) showed significantly reduced ROCK-2 mRNA expression compared to WT. Data are presented as mean ± SEM with individual data points shown. Statistics: one-way ANOVA with Tukey's *post hoc* test for multiple comparisons.

**d.** Schematic of the experimental workflow. Human iPSCs with ROCK-2 CRISPR knockdown (sgRNA1 and sgRNA2) were differentiated into cortical neurons and cultured in 96-well plates. At DIV 40, neurons were subjected to a scrape injury. Neurons were fixed 3 days post-injury (DIV 43) for immunostaining and 10X confocal imaging.

**e.** Representative confocal images of βIII-tubulin-stained human cortical neurons at 3 days post-injury from wild-type (WT, left), ROCK-2 KD sgRNA1 (L1, middle), and ROCK-2 KD sgRNA2 (L2, right) lines, with corresponding βIII-tubulin thresholded binary images below. Red dashed boxes indicate regions of interest at the injury site. Both ROCK-2 KD lines display visibly enhanced axonal regrowth into the injury zone compared to WT. Scale bar: 100 μm.

**f.** Quantification of the percentage area of regrowth. Sample sizes (number of wells of 96-well plate analyzed): WT (*n* = 36), ROCK-2 KD L1 (*n* = 38), ROCK-2 KD L2 (*n* = 36). Both L1 (*P* < 0.0001) and L2 (*P* < 0.0001) significantly increased regrowth compared to WT. No significant difference was observed between L1 and L2. Data are presented as mean ± SEM with individual data points shown. Statistics: one-way ANOVA with Tukey's *post hoc* test for multiple comparisons. All experiments were conducted in *N* = 3 independent replicates. NS, not significant; **P < 0.01; ***P < 0.001; ****P < 0.0001. P-values are indicated on the graphs.

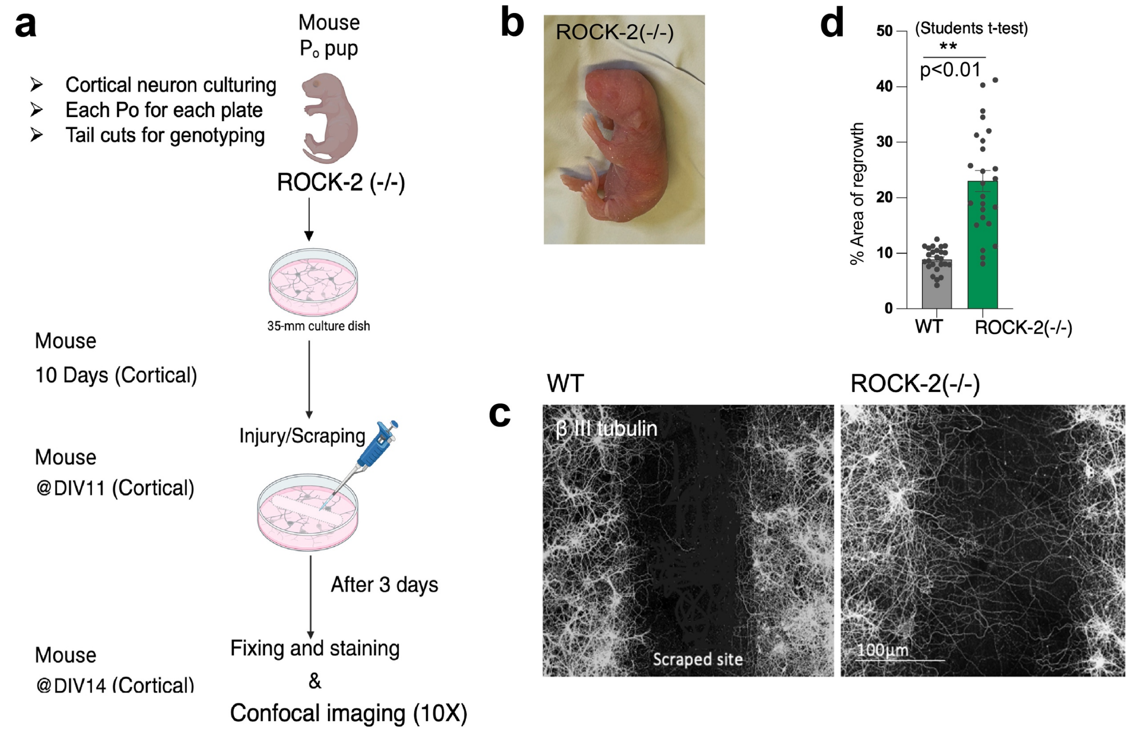


**Supplementary Figure 6: Genetic deletion of ROCK-2 enhances axon regrowth in mouse cortical neurons.**

**a.** Schematic of the experimental workflow. Cortical neurons were cultured from individual P0 mouse pups, with each pup plated on a separate 35-mm culture dish and tail cuts collected for genotyping. ROCK-2(−/−) Knockout pups were identified by genotyping, and their cortical neurons were cultured for 10 days. At DIV 11, neurons were subjected to a scrape injury. Neurons were fixed 3 days post-injury (DIV 14) for immunostaining and 10X confocal imaging.

**b.** Photograph of a ROCK-2(−/−) KO P0 mouse pup.

**c.** Representative confocal images of βIII-tubulin-stained cortical neurons at 3 days post-injury from wild-type (WT-littermate, left) and ROCK-2(−/−) KO (right) mice. The scraped injury site is indicated. ROCK-2(−/−) KO neurons display visibly enhanced axonal regrowth into the injury zone compared to WT neurons. Scale bar: 100 μm.

**d.** Quantification of the percentage area of regrowth in WT and ROCK-2(−/−) KO mouse cortical neurons at 3 days post-injury. ROCK-2(−/−)-KO neurons showed significantly greater regrowth compared to WT-littermate (P < 0.01). Data are presented as mean ± SEM with individual data points shown. Statistics: unpaired two-tailed Student's *t*-test. **P < 0.01. P-values are indicated on the graph. Data for (**D**) are available in the Source Data file.

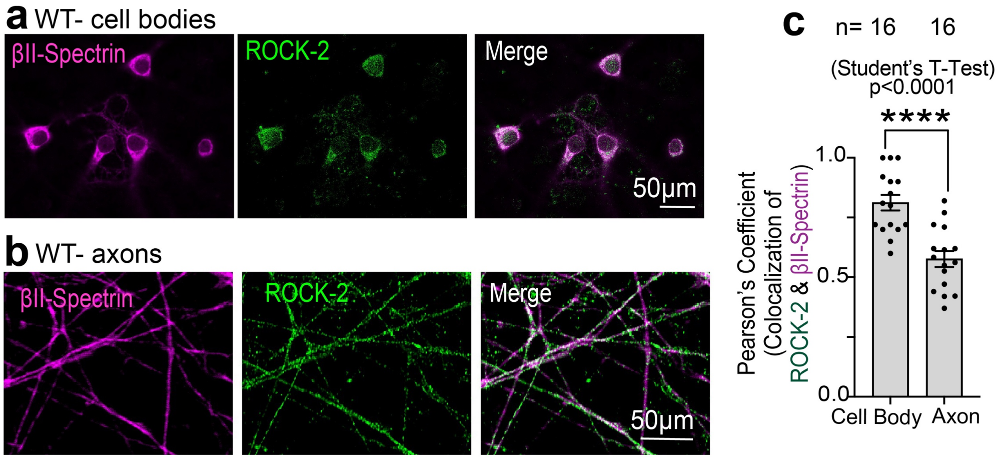


**Supplementary Figure 7: ROCK-2 kinase shows periodic organization and co-localizes with βII-spectrin in intact axons of control (GMK2) iPSC-derived human cortical neurons.**
**a-b.** Confocal images of wild-type human cortical neuron cell bodies (**a**) and axons (**b**) immunostained for βII-spectrin (magenta), ROCK-2 (green), and merged channels. ROCK-2 is expressed in both compartments and partially co-localizes with βII-spectrin. Scale bars, 50 μm.
**c.** Pearson's correlation coefficient for ROCK-2/βII-spectrin co-localization in cell bodies and axons. Co-localization was significantly higher in cell bodies than in axons (P < 0.0001). Sample sizes: *n* = 16 cell bodies, *n* = 16 axons. Data are mean ± SEM, with individual data points shown. Unpaired two-tailed Student's *t*-test. ****P < 0.0001.

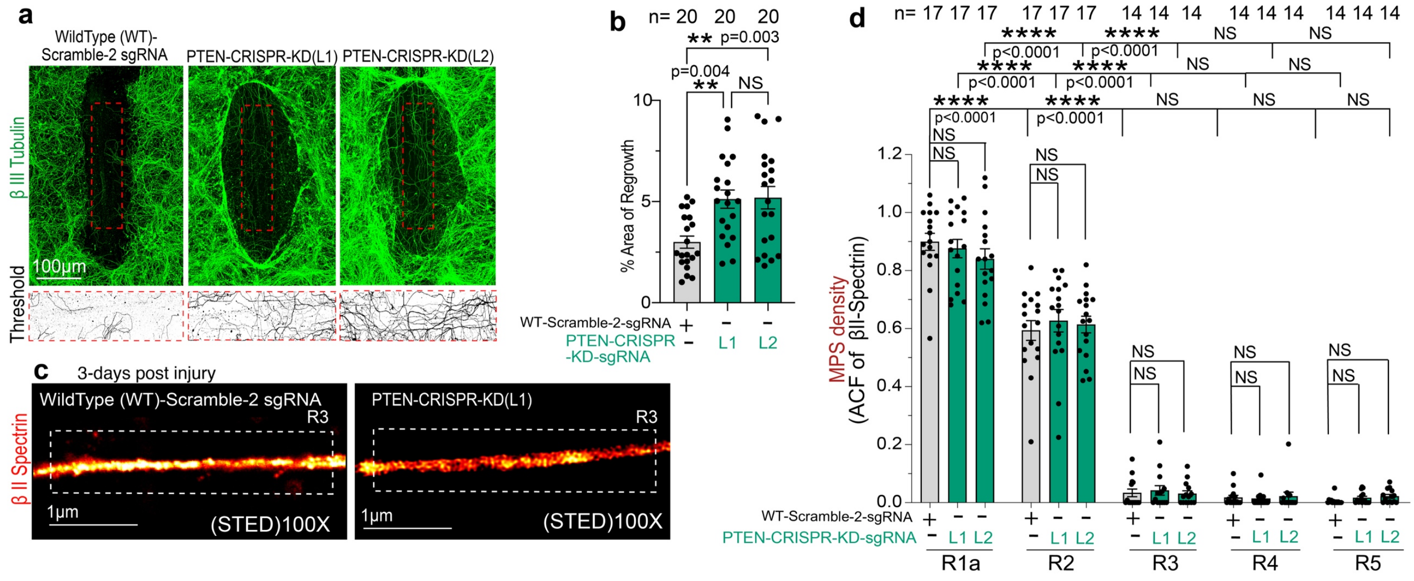


**Supplementary Figure 8: PTEN inhibition by CRISPRi enhances regrowth capacity but cannot restore MPS periodic arrangement after 4 days of injury.**
**a.** Confocal images of the scrape injury site at 3 dpi in wild-type (WT + Scramble-2 sgRNA), PTEN-CRISPR-KD (L1), and PTEN-CRISPR-KD (L2) human cortical neurons immunostained for βIII-tubulin (green). Bottom panels show thresholded binary images of βIII-tubulin within the indicated regions (red dashed boxes). Both PTEN-KD lines display enhanced regrowth compared with WT. Scale bar, 100 μm.
**b.** Percentage area of regrowth. Both PTEN-KD lines showed significantly increased regrowth compared with WT (L1, *P* = 0.004; L2, *P* = 0.003), with no difference between L1 and L2 (NS). n (number of wells in 96 well plate) = 20 per group. Data are mean ± SEM, with individual data points shown. One-way ANOVA with Tukey's *post hoc* test. **P < 0.01.
**c.** STED super-resolution images of βII-spectrin (red hot) at R3 in WT (left) and PTEN-CRISPR-KD (L1) (right) axons at 3 dpi. Despite enhanced regrowth, PTEN-KD neurons show no improvement in βII-spectrin periodicity compared with WT. Dashed boxes indicate segments used for autocorrelation analysis. Scale bars, 1 μm.

**d.** MPS density (ACF of βII-spectrin) across regions R1a–R5 in WT, PTEN-KD (L1), and PTEN-KD (L2) neurons at 3 dpi. At R1a, MPS density was comparable across genotypes (NS). All genotypes showed reduced periodicity at R2 relative to R1a ($P < 0.0001$), but no differences among genotypes within any region (NS at R2–R5). MPS density was near zero at R3, R4, and R5 in all conditions. Sample sizes: R1a (*n* = 17, 17, 17), R2 (*n* = 17, 17, 17), R3–R5 (*n* = 14, 14, 14 each). Data are mean ± SEM with individual data points; *N* = 3 independent replicates. One-way ANOVA with Tukey's *post hoc* test. NS, not significant; ****$P < 0.0001$.

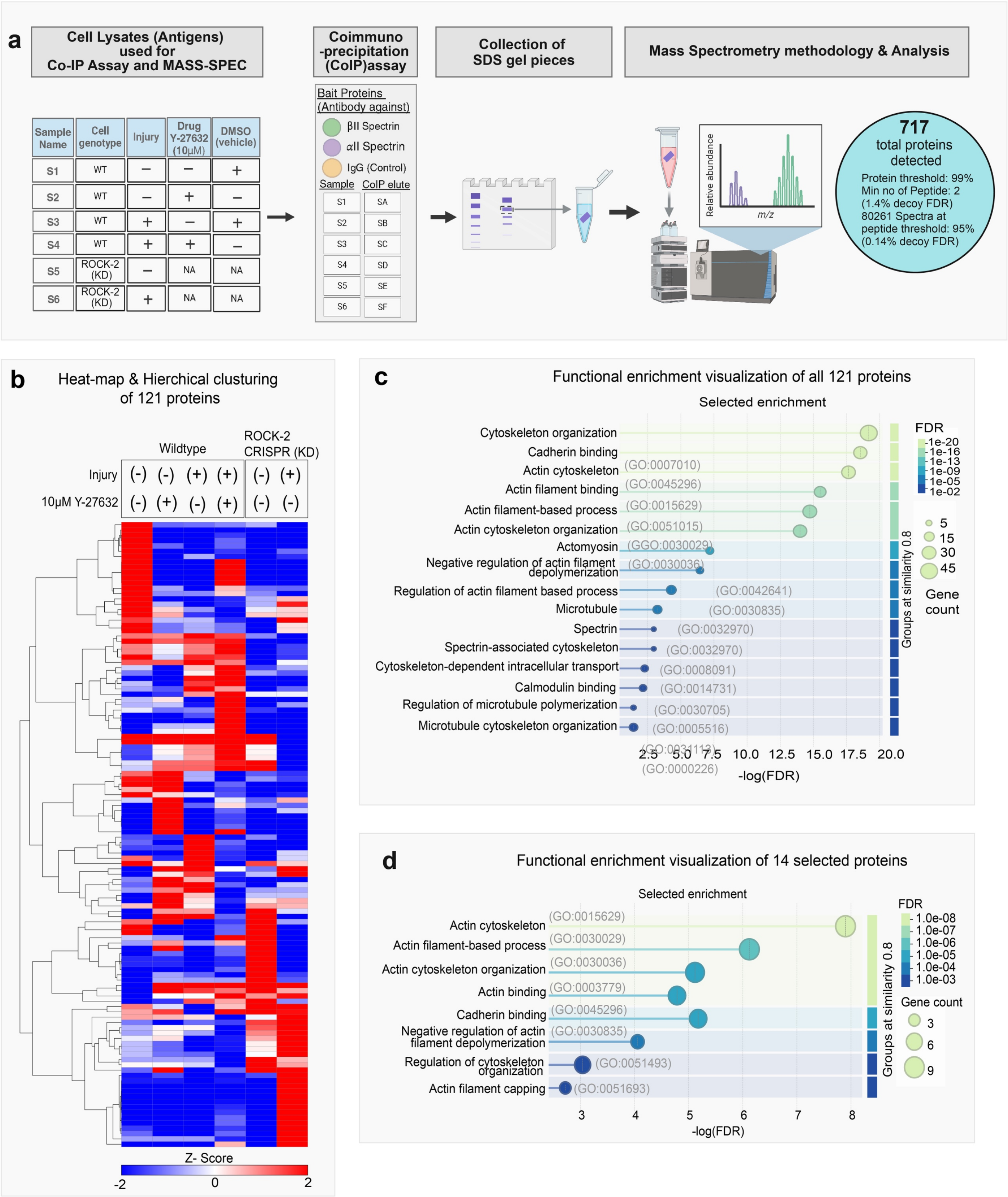


**Supplementary Figure 9: Proteomic mass spectrometry analysis and GO term enrichment analysis reveal upregulation of MYO5A protein during injury and ROCK-2 inhibition.**

**a.** Schematic of the experimental and proteomic workflow. Cell lysates from human iPSC-derived cortical neurons were prepared under six conditions: S1, wild-type (WT) uninjured with vehicle (1× PBS); S2, WT uninjured with 10 μM Y-27632; S3, WT injured with vehicle; S4, WT injured with 10 μM Y-27632; S5, ROCK-2 CRISPR KD line (l1) uninjured (NA); S6, ROCK-2 CRISPR KD L1injured (NA). Lysates were subjected to co-immunoprecipitation (CoIP) using antibodies against βII-spectrin and αII-spectrin as bait proteins, with IgG as a control. CoIP eluates (SA–SF) were separated by SDS-PAGE, gel pieces were collected, and 717 proteins were identified by mass spectrometry.

b. Heatmap illustrating hierarchical clustering of Z-score-normalized 121 protein abundance across six conditions. Each row represents an individual protein, and each column represents a condition. This clustering reveals distinct protein expression patterns between uninjured vehicle-treated, Y-27632-treated, injured, and ROCK-2 CRISPR KD L1 conditions. The color scale represents Z-scores ranging from −2 (red, indicating lower abundance) to +2 (blue, indicating higher abundance).

c. Functional enrichment analysis of all 121 proteins identified in the βII-spectrin interactome. Selected Gene Ontology (GO) terms are shown, with circle size representing gene count and color representing false discovery rate (FDR). The most enriched categories include cytoskeleton organization (GO:0007010), cadherin binding (GO:0045296), actin cytoskeleton (GO:0015629), actin filament binding (GO:0051015), and actin filament-based process (GO:0030029). Spectrin (GO:0008091) and spectrin-associated cytoskeleton (GO:0014731) terms confirm the specificity of the pull-down. The x-axis shows −log (FDR).

**d.** Functional enrichment analysis of 14 selected proteins that were differentially expressed upon injury and/or ROCK-2 inhibition. Top enriched GO terms include actin cytoskeleton (GO:0015629), actin filament-based process (GO:0030029), actin cytoskeleton organization (GO:0030036), actin binding (GO:0003779), and cadherin binding (GO:0045296). The x-axis shows −log(FDR); circle size represents gene count, and color represents FDR. GO term enrichment analysis was performed using STRING tool with FDR correction.

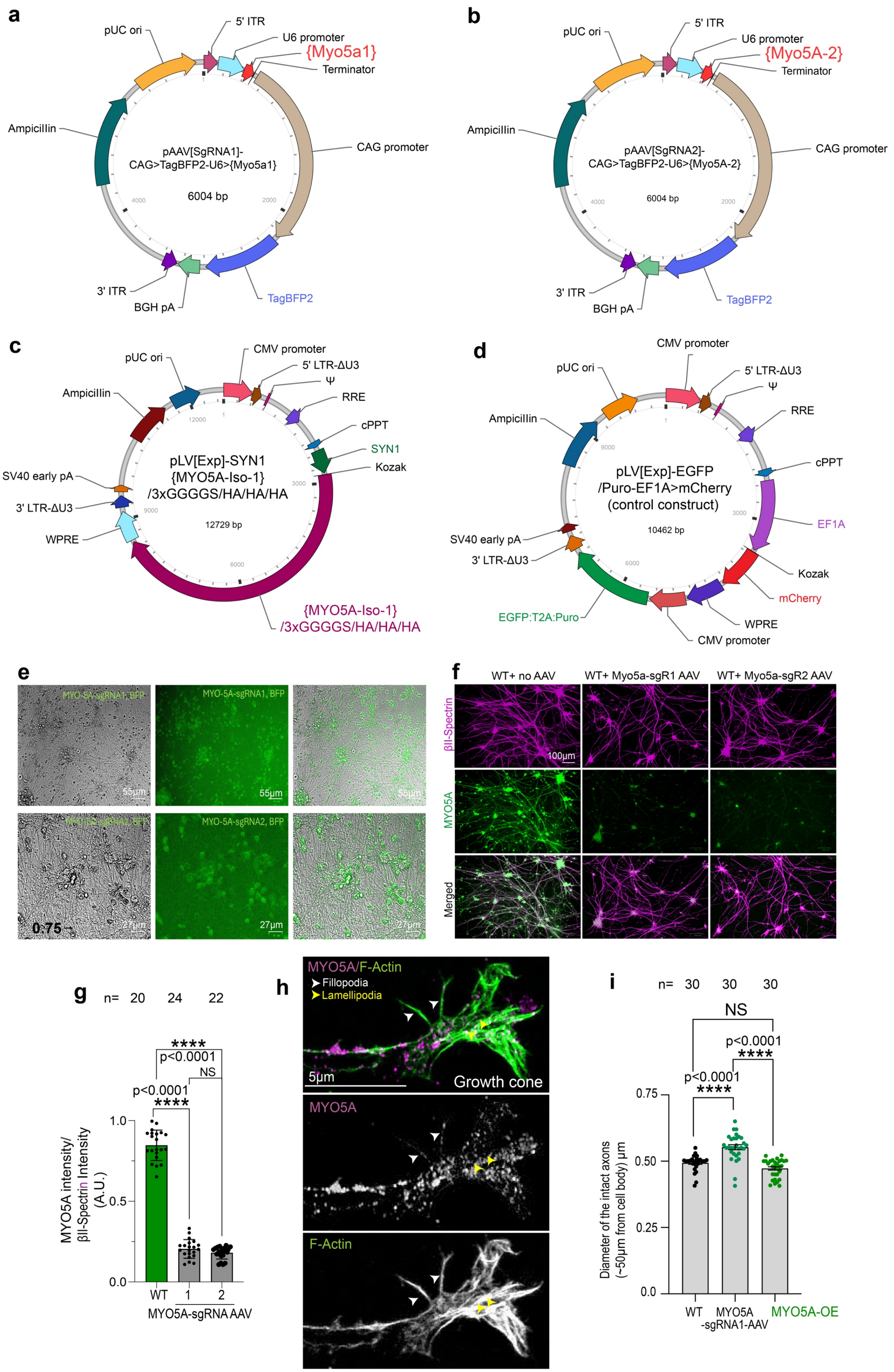
a
pUC ori
5' ITR
U6 promoter
{Myo5a1}
Terminator
Ampicillin
pAAV[SgRNA1]-
CAG>TagBFP2-U6>{Myo5a1}
6004 bp
4000
2000
CAG promoter
3' ITR
BGH pA
TagBFP2
b
pUC ori
5' ITR
U6 promoter
{Myo5A-2}
Terminator
Ampicillin
pAAV[SgRNA2]-
CAG>TagBFP2-U6>{Myo5A-2}
6004 bp
CAG promoter
3' ITR
BGH pA
TagBFP2
c
pUC ori
CMV promoter
5' LTR-ΔU3
Ψ
RRE
Ampicillin
cPPT
SYN1
Kozak
SV40 early pA
3' LTR-ΔU3
WPRE
pLV[Exp]-SYN1
{MYO5A-Iso-1}
/3xGGGGS/HA/HA/HA
12729 bp
{MYO5A-Iso-1}
/3xGGGGS/HA/HA/HA
d
CMV promoter
pUC ori
5' LTR-ΔU3
Ψ
Ampicillin
RRE
cPPT
pLV[Exp]-EGFP
/Puro-EF1A>mCherry
(control construct)
10462 bp
EF1A
SV40 early pA
3' LTR-ΔU3
Kozak
mCherry
EGFP:T2A:Puro
WPRE
CMV promoter
e
MYO-5A-sgRNA1, BFP
55µm
MYO-5A-sgRNA2, BFP
27µm
f
WT+ no AAV
WT+ Myo5a-sgR1 AAV
WT+ Myo5a-sgR2 AAV
βII-Spectrin
MYO5A
Merged
100µm
g
n= 20 24 22
****
p<0.0001
NS
p<0.0001
****
MYO5A intensity/
βII-Spectrin Intensity
(A.U.)
1.0
0.5
0.0
WT
1
2
MYO5A-sgRNA AAV
h
MYO5A/F-Actin
➤ Fillopodia
➤ Lamellipodia
5µm
Growth cone
MYO5A
F-Actin
i
n= 30 30 30
NS
p<0.0001
****
p<0.0001
****
0.75
0.50
0.25
0.0
Diameter of the intact axons
(~50µm from cell body) µm
WT
MYO5A
-sgRNA1-AAV
MYO5A-OE

**Supplementary Figure 10: Construct design, validation of MYO5A knockdown, MYO5A localization at growth cones, and effect of MYO5A perturbation on axon diameter.**

**a-b.** Vector maps of MYO5A-sgRNA1 (**a**) and MYO5A-sgRNA2 (**b**) AAV constructs (pAAV[SgRNA]-CAG>TagBFP2-U6>; 6004 bp each), containing U6-driven sgRNAs and CAG-driven TagBFP2 reporter, flanked by ITRs. **c-d.** Vector maps of the MYO5A-overexpression lentiviral construct (pLV[Exp]-SYN1-(MYO5A-Iso-1)/3×GGGGS/HA/HA/HA; 12729 bp)

**c-d**. Vector Maps of the lentiviral constructs used in MYO5A gain-of-function experiments. (c) The overexpression vector (pLV [Exp]-SYN1-{MYO5A-iso-1}/3×GGGGS/HA/HA/HA; 12,729 bp) features the neuron-specific SYN1 promoter driving MYO5A isoform 1, fused at its C-terminus via a 3×GGGGS linker to a 3×HA tag. d: The control vector (pLV [Exp]-EGFP/Puro-EF1A>mCherry; 10,462 bp) utilizes EF1A to drive mCherry expression and a CMV promoter to control an EGFP: T2A:Puro reporter/puromycin-selection cassette.

**e.** Phase-contrast (left) and fluorescence (right) images of human cortical neurons transduced with MYO5A-sgRNA1 (top) or MYO5A-sgRNA2 (bottom) AAV, confirming successful transduction via TagBFP2 expression. Scale bars, 55 µm (top) and 27 µm (bottom).

**f.** Confocal images of WT neurons with no AAV, MYO5A-sgRNA1 AAV, or MYO5A-sgRNA2 AAV, immunostained for βII-spectrin (magenta), MYO5A (green), and merged channels. MYO5A signal is markedly reduced in both sgRNA conditions. Scale bar, 100 µm.

**g.** MYO5A fluorescence intensity normalized to βII-spectrin. Both sgRNAs significantly reduced MYO5A levels ($P < 0.0001$), with no difference between them (NS). $n$ = 20, 24, 22. Data are mean ± SEM with individual data points; $N$ = 3 independent replicates. One-way ANOVA with Tukey's *post hoc* test. ****$P < 0.0001$.

**h.** Confocal images of a growth cone immunostained for MYO5A (green) and F-actin (magenta). MYO5A co-localizes with F-actin in filopodia (white arrowheads) and lamellipodia (yellow arrowheads). Scale bar, 5 µm.

**i.** Axon diameter measured ~50 µm from the cell body in intact axons. MYO5A knockdown significantly increased diameter compared with WT ($P < 0.0001$), while MYO5A-OE did not alter diameter (NS). Data are mean ± SEM with individual data points; $N$ = 3 independent replicates. One-way ANOVA with Tukey's *post hoc* test. ****$P <$ 0.0001.

**Legends for Supplementary Data Files 1-3 (XLSX format)**

**Supplementary Data 1: Complete co-immunoprecipitation–mass spectrometry (CoIP-MS) dataset of the βII-spectrin interactome in human iPSC-derived cortical neurons.** The full Scaffold proteomic report underlying Fig. 4 and Supplementary Fig. 9 details 717 proteins identified across all eluates, organized into various protein clusters. The report header outlines the search parameters: spectra were analyzed with Mascot (v2.8.3, within Proteome Discoverer) against the Swiss-Prot Homo sapiens database, using trypsin digestion (up to 2 missed cleavages), a 10-ppm parent ion tolerance, and a 0.02 Da fragment tolerance. Carbamidomethylation of cysteine was set as a fixed modification, while methionine oxidation was variable. Proteins were filtered for >99% probability, at least 2 peptides identified, and >95% peptide probability. Each row lists the protein cluster/group ID, the identified protein, the UniProt accession number, the gene symbol, the molecular weight, and the quantitative, normalized spectral counts for 18 LC-MS/MS runs. These runs represent six experimental conditions (S1–S6), each with three CoIP baits (A: βII-spectrin; B: αII-spectrin; C: IgG control): S1-WT uninjured + vehicle; S2-WT uninjured + 10 μM Y-27632; S3-WT injured + vehicle; S4-WT injured + 10 μM Y-27632; S5-ROCK-2-CRISPR-KD (line 1) uninjured; S6-ROCK-2-CRISPR-KD (line 1) injured. IgG-control runs show minimal detection of spectrin complex proteins (e.g., SPTBN1, SPTAN1, ANK2, MYO5A), confirming pull-down specificity. Decoy hits are marked "-DECOY" in the accession field.

**Supplementary Data 2: Curated 121-protein βII/αII-spectrin interactome with normalized spectral counts across conditions.** The 121 high-confidence interactome proteins, identified with their UniProt accession, protein name, and normalized total spectral counts for βII-spectrin (A) and αII-spectrin (B) baits across six conditions: S1, wild-type (WT) uninjured without Y-27632; S2, WT uninjured with 10 μM Y-27632; S3, WT injured without Y-27632; S4, WT injured with 10 μM Y-27632; S5, ROCK-2-CRISPR-KD (line 1) uninjured; S6, ROCK-2-CRISPR-KD (line 1) injured. These proteins, co-precipitated by both spectrin baits after IgG-control background subtraction, form the basis of Fig. 4 b,c and Supplementary Fig. 9b–d.

**Supplementary Data 3: The 14 spectrin interactors regulated by injury or ROCK-2, normalized to βII-spectrin.** The 14 candidate interactors, along with their UniProt accession, protein, and gene names, are listed, including the βII-spectrin bait (SPTBN1) used as a normalization reference. For each of the six conditions—(S1) WT uninjured, no Y-27632; (S2) WT uninjured + 10 μM Y-27632; (S3) WT injured, no Y-27632; (S4) WT injured + 10 μM Y-27632; (S5) ROCK-2-CRISPR-KD line 1 uninjured; (S6) ROCK-2-CRISPR-KD line 1 injured—normalized total spectral counts are provided for the βII-spectrin (A) and αII-spectrin (B) baits, each expressed as a ratio to βII-spectrin (/βIISpec). These values underlie the z-score heatmap and STRING network shown in Fig. 4 c,d, the MYO5A quantification in Fig. 4e, and Supplementary Fig. 9d.
